# Regulation of cell proliferation by a novel feedback system on Cdk function

**DOI:** 10.1101/2025.10.28.685054

**Authors:** Joseph C. Ryan, Baptiste Leray, Akanksha Jain, Nelson Coehlo, Maria Rosa Domingo-Sananes, Theo Aspert, Gilles Charvin, Matthias Simoes Krockenberger, Nathalia Chica Blaguera, Sandra Lopez Aviles, Luca Takacs, Frank Uhlmann, Ludong Yang, Jia-Xing Yue, Gianni Liti, Victor Cochard, Guillaume Chevreux, Hironori Sugiyama, Yuhei Goto, Kazuhiro Aoki, Pei-Yun Jenny Wu, Cameron Mackereth, John Tyson, Béla Novák, Damien Coudreuse

## Abstract

The proliferation of eukaryotic cells is regulated by a complex network of regulatory systems that promotes efficient cell cycle progression and ensures proper responses to the environment. Despite this complexity, the core inputs that are necessary and sufficient for robust alternation of DNA replication and mitosis are surprisingly simpler than anticipated. Indeed, fission yeast cells operating with an engineered minimal cell cycle network that lacks the highly conserved Wee1+Cdc25 feedback loops on Cdk1 function are viable, although slow growing. This provides a unique entry for evaluating how such simplified cells can evolve and improve their proliferation potential while exploring unknown mechanisms modulating cell cycle progression. Taking advantage of this model, we applied laboratory evolution assays to minimal fission yeast backgrounds and selected for the emergence of faster growing populations. We found that loss of the small disordered protein Spo12 brings about enhanced population growth in cells lacking the Wee1+Cdc25 mitotic switch. Importantly, we demonstrate that Spo12 defines a new and conserved family of inhibitors of the Cdk-counteracting phosphatase PP2A that are directly regulated by Cdk-dependent phosphorylation. Our results also reveal a trade-off associated with Spo12-dependent regulation, which may have implications for our understanding of the principles underlying the evolution of cell cycle control. Finally, our study highlights how combining simplified circuits with experimental evolution allows for uncovering regulatory elements that may be obscured by network complexity.

## Introduction

Regulation of the eukaryotic cell cycle shapes the behavior of both uni- and multicellular organisms. However, despite considerable knowledge of the multi-layered control of cell proliferation^1,2^, our understanding of the core mechanisms that are necessary and sufficient to ensure robust cell division and maintain cellular homeostasis remains surprisingly incomplete. Substantial evidence supports the idea that qualitative changes in the activities of different cyclin-dependent kinases (Cdks), through their association with distinct cyclins, are central to the alternation of DNA synthesis and mitosis during eukaryotic cell proliferation^1–4^. Nevertheless, a number of studies have demonstrated that quantitative modulation of the activity of a single, oscillating Cdk can drive faithful progression through the cell cycle^5–9^ (Fig. 1a).

**Figure 1.**
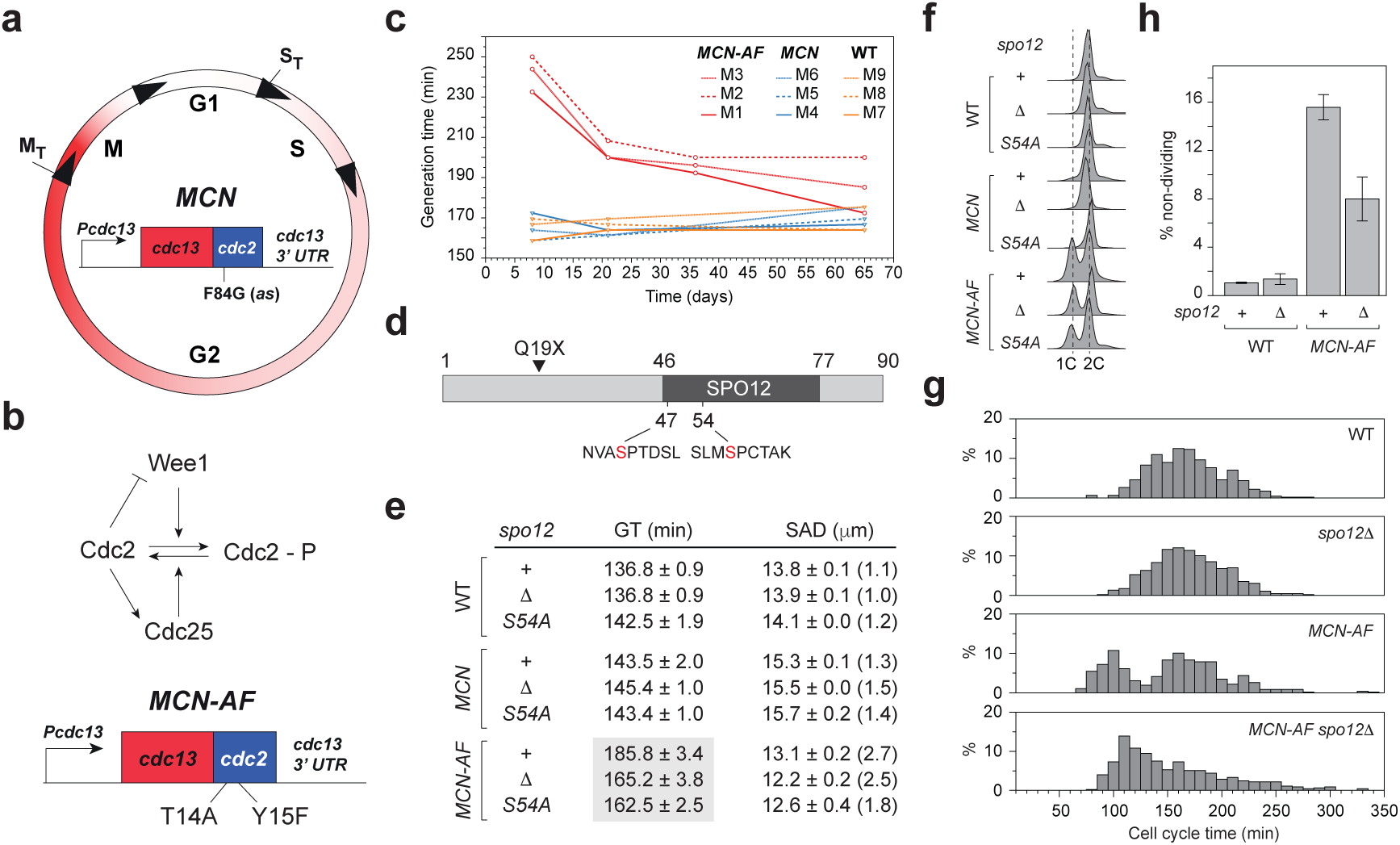
Laboratory evolution assays identify Spo12 as a regulator of Cdc2 function. **a.** Schematic of the Minimal Cell cycle Network (*MCN*). Fission yeast cell proliferation can be driven by oscillations of a single quantitative Cdk activity between a low S phase threshold (S_T_) and a high M phase threshold (M_T_)^6^. In *MCN* cells, cell cycle progression solely relies on a fusion between Cdc13 (cyclin B) and Cdc2 (Cdk1). Expression of the *MCN* module is controlled by the *cdc13* regulatory elements (*Pcdc13*: *cdc13* promoter and *cdc13* 3’UTR). In these cells, the endogenous copies of *cdc13*, *cdc2* and the cell cycle cyclin expressing genes *cig1*, *cig2* and *puc1* are deleted. Red gradient represents the changes in activity of the Cdc13-Cdc2 fusion. A substitution in the Cdc2 moiety of the fusion protein (F84G) renders the MCN module sensitive to reversible and dose-dependent inhibition by non-hydrolysable ATP analogs (*e.g.*, 3-MBPP1; *as*: analog-sensitive)^6^. **b.** *Top panel*: mitotic entry is controlled by the conserved Wee1+Cdc25 feedback loops on Cdc2 activity. The Wee1 kinase inhibits Cdc2 by phosphorylation on Y15 throughout G2, a modification that is counteracted by the phosphatase Cdc25. Cdc2 in turn inhibits Wee1 and activates Cdc25. This system generates a bistable switch that underlies the burst in Cdk activity that triggers mitotic onset. Note that Wee1 also phosphorylates Cdc2 on T14, although the function of this modification remains unclear^15^. *Bottom panel*: in *MCN-AF* cells, substitution of T14 (T14A) and Y15 (Y15F) uncouples the activity of the Cdc13-Cdc2 fusion from Wee1+Cdc25 regulation. While these alterations are lethal in a wild-type (WT) background, *MCN-AF* cells are alive although slow growing (see *e*). **c.** Changes in population generation times of the different populations throughout the evolution experiment (Extended data Fig. 1). Generation times were determined by inoculating batch cultures with a sample of each evolving population. An improvement in population growth was observed for all *MCN-AF* cultures. Note that all experiments using samples or clones from the evolving populations (Extended data Fig. 1) were performed in the evolution conditions (supplemented minimal medium, 3% galactose, 28 °C), leading to slower doubling times compared to those reported in *e, g* and Extended data Fig. 6 (32 °C) as well as Fig. 3 and Extended data Fig. 5 (30 °C). This lower temperature was used to limit flocculation in the evolution assay^17^, and similar conditions were applied for all initial characterizations. **d.** Spo12 is a short protein of 90 amino acids. It contains a SPO12 domain (46-77) as well as two consensus Cdk target sites (S47 and S54). An early stop codon (Q19X) in Spo12 was identified in one of the evolved *MCN-AF* populations (M3). **e.** Population generation time (GT) and cell size at division (SAD) for cells lacking Spo12 function (*spo1211* and *spo12(S54A)*) in the WT, *MCN* and *MCN-AF* genetic backgrounds. Averages of three independent replicates (*n*>100 for each replicate for SAD) with standard errors (standard deviations over the pooled datasets for SAD are shown in parentheses). GT corresponds to the culture doubling time based on optical density measurements, taking into account both proliferating cells and cells that do not divide. It therefore differs from cell cycle time determination (as in *g*), which is restricted to dividing cells. **f.** DNA content analysis of cells in *e*. In fission yeast, the appearance of a 1C DNA content reflects an abnormal extension of G1. Loss of Spo12 reduces the size of the 1C peak in the *MCN-AF* background, suggesting a change in cell cycle phase organization, in particular a shortening of G1. **g.** Single-cell analyses from time-lapse experiments of cell cycle time for the indicated genotypes. Pooled datasets from three independent replicates (*n*>150 for each replicate). Average and standard deviation (pooled datasets): WT, 168 ± 34 min; *spo1211*, 170 ± 33 min; *MCN-AF*, 154 ± 53 min; *MCN-AF spo1211*, 154 ± 50 min. See Extended data Fig. 2e for accompanying size at division measurements. **h.** Percentage of cells that did not divide over the course of 10 h time-lapse experiments for the indicated genotypes. Averages of three independent experiments with standard errors. *n*>80 for each replicate.

In the fission yeast *Schizosaccharomyces pombe*, the cell cycle Cdk-cyclin circuit can be replaced by a single fusion protein consisting of Cdc13/cyclin B and Cdc2/Cdk1^6^ (Minimal Cell cycle Network, *MCN*, Fig. 1a). As is the case in many eukaryotes, commitment to mitosis in fission yeast also largely relies on a conserved positive feedback control of Cdk function that is mediated by the Cdk-inhibitory kinase Wee1 and its counteracting phosphatase Cdc25^10–13^ (Fig. 1b). These dual feedback loops lie at the heart of the proposed bistability and switch-like behavior of the cell cycle network^13^. Surprisingly, while the timely modulation of Cdc2 phosphorylation by Wee1+Cdc25 is essential for the proliferation and viability of *S. pombe* cells, it is dispensable in cells operating with a simplified cell cycle system^6,14^. Indeed, *MCN*-derived cells in which the Wee1+Cdc25 loops are disabled through alteration of the Wee1 target sites in the Cdc2 moiety of the fusion protein (T14A and Y15F^12,15^, *MCN-AF*, Fig. 1b) are alive, although slower growing^6^.

From an evolutionary perspective, the demonstration that a simplified, genetically-engineered system can effectively sustain cell division highlights the plasticity in the regulation of cell proliferation. It also raises intriguing and largely unexplored questions on the way cells can adapt and enhance their division potential. In particular, this provides a unique experimental model for evaluating the strategies that may have been employed by natural cells operating with cell cycle networks simpler than those of present-day cells to cope with selective environments. Would proliferation improvements be systematically brought about by establishing *de novo* complexity in cell cycle control? Alternatively, phases of simplification through loss of existing functions might shift the balance between different elements in a given cell cycle circuit and thereby promote cell proliferation. Assessing these possibilities using experimental evolution may bring new insights into the way cell growth and division can evolve and help us understand different network topologies that may drive cell cycle progression in distinct eukaryotic species. In addition, surveying the genetic and mechanistic space that cells can explore to increase population growth may uncover important regulatory features of the cell cycle that have so far been obscured in laboratory conditions.

To address these questions, we subjected *MCN-AF* populations, which lack the mitotic switch, to laboratory evolution assays. To this end, cultures were maintained under constant selective pressure for efficient proliferation, promoting the emergence of faster-growing, evolved clones. Our results led us to identify the small disordered protein Spo12 as a founding member of a new and conserved family of cell cycle regulators. First, using population and single-cell analyses, we show that loss of Spo12 in the *MCN-AF* background improves population growth. Importantly, *in vivo* Cdk activity measurements, proteomic approaches, *in silico* protein structure prediction, and mathematical modeling demonstrated that Spo12 is a stoichiometric inhibitor of the conserved Cdk-counteracting phosphatase PP2A and part of a novel feedback system on Cdk function. In contrast to known PP2A inhibitors, Spo12 is directly targeted by Cdk1-dependent phosphorylation, defining a distinct class of regulators of this phosphatase complex. Our findings also indicate that the Spo12 control loop comes with a trade-off, highlighting the complexity of the opposing forces that shape the evolution of the cell cycle network.

### Experimental evolution of *MCN-AF* cells

We set out to isolate evolved *MCN-AF* clones that show increased population growth rates using laboratory evolution assays. To this end, triplicates of *MCN-AF,* as well as of *MCN* and wild-type (WT) controls, were maintained in a continuous proliferative state and followed for 70 days in a turbidostat system^16,17^ (Extended data Fig. 1a). In this setup, which operates at constant culture volume and cell density, slow perfusion of fresh medium is compensated by an equivalent efflux of cells, thereby imposing a selection pressure for faster proliferating clones. All *MCN-AF* cultures and individual tested clones showed significant growth improvement over time, in contrast to *MCN* and WT populations (Fig. 1c, Extended data Fig. 1b). This was accompanied by changes in cell proliferation-associated phenotypes, including 1) cell cycle phase distribution, with a shortening of the long G1 phase observed in the parental strain^6^ (Extended data Fig. 1c, d), and 2) cell size at division, which tended to both decrease and become more homogenous in two of the cultures (Extended data Fig. 1e, f).

Whole-genome sequencing of both heterogenous evolved populations and individual clones isolated from the different cultures revealed complex genetic trajectories throughout the evolution assay (Supplementary Table 1). Indeed, distinct sub-genotypes clearly emerged within each of the *MCN-AF* turbidostats, with relatively limited mutational overlap between replicates but significant clonal interference (Extended data Fig. 1g). In one out of the three *MCN-AF* populations (M1), a reversion of the Y15F alteration in Cdc2, but not of the T14A substitution, was detected, demonstrating the strong selective advantage of the Wee1+Cdc25 feedback loops on Cdc2 function. We next selected the M3 culture for further study due to a greater phenotypic stability between the mid-point and the end of the evolution assay compared to the M1 and M2 populations. Interestingly, we identified the emergence of an evolved allele of *spo12* in this culture. *spo12* was previously found as a multicopy suppressor of the cell cycle arrest induced by the temperature-sensitive (*ts*) allele *mcs3-12^ts^*in the *wee1-50^ts^ cdc25-22^ts^* background^18^. Although the identity of the *mcs3* gene remains unknown, this hinted at a genetic interaction between Spo12 and the Wee1+Cdc25 loops and a potential role for Spo12 in the cell cycle control network.

Spo12 is a relatively small protein (90 amino acids) whose only well-defined feature is a SPO12 domain, shared with *Saccharomyces cerevisiae* Spo12. Budding yeast Spo12 is part of the FEAR network and contributes to the release of the Cdc14 phosphatase from the nucleolus at mitotic exit through an unknown molecular mechanism^19–22^. In contrast, the function and regulation of *S. pombe* Spo12, which does not contribute to regulating the localization of the Cdc14 homolog Clp1^23^, remained elusive. The mutation that we identified in our evolved population generates an early stop codon (Q19X, Fig. 1d). Furthermore, this loss-of-function allele of *spo12* co-segregated with the improved growth phenotype upon backcross of an M3 clone with the parental, non-evolved strain (Extended data Fig. 1h). Altogether, these data implicated Spo12 as a strong candidate for modulating the proliferation of cells lacking the Wee1+Cdc25 mitotic switch.

### Loss of Spo12 function enhances the proliferation of *MCN-AF* cells

While a full deletion of the *spo12* open reading frame in WT and *MCN* cells did not result in any apparent cell cycle phenotypes, it significantly improved the population growth rate of *MCN-AF* cells (Fig. 1e; Extended data Fig. 2a). As anticipated from the characterization of the evolving populations, loss of Spo12 function was sufficient to partially revert the prolonged G1 in *MCN-AF* cells (Fig. 1f) and had a modest effect on their size at division (Fig. 1e; Extended data Fig. 2b). Consistent with a previous report, it did not impact meiotic progression^23^ (Extended data Fig. 2c). Interestingly, deletion of *spo12* also partially rescued the hypersensitivity of *MCN-AF* cells to various replication stresses (Extended data Fig. 2d), despite the absence of checkpoint signaling through Cdc2/Cdk1 in this background^24,25^.

**Figure 2.**
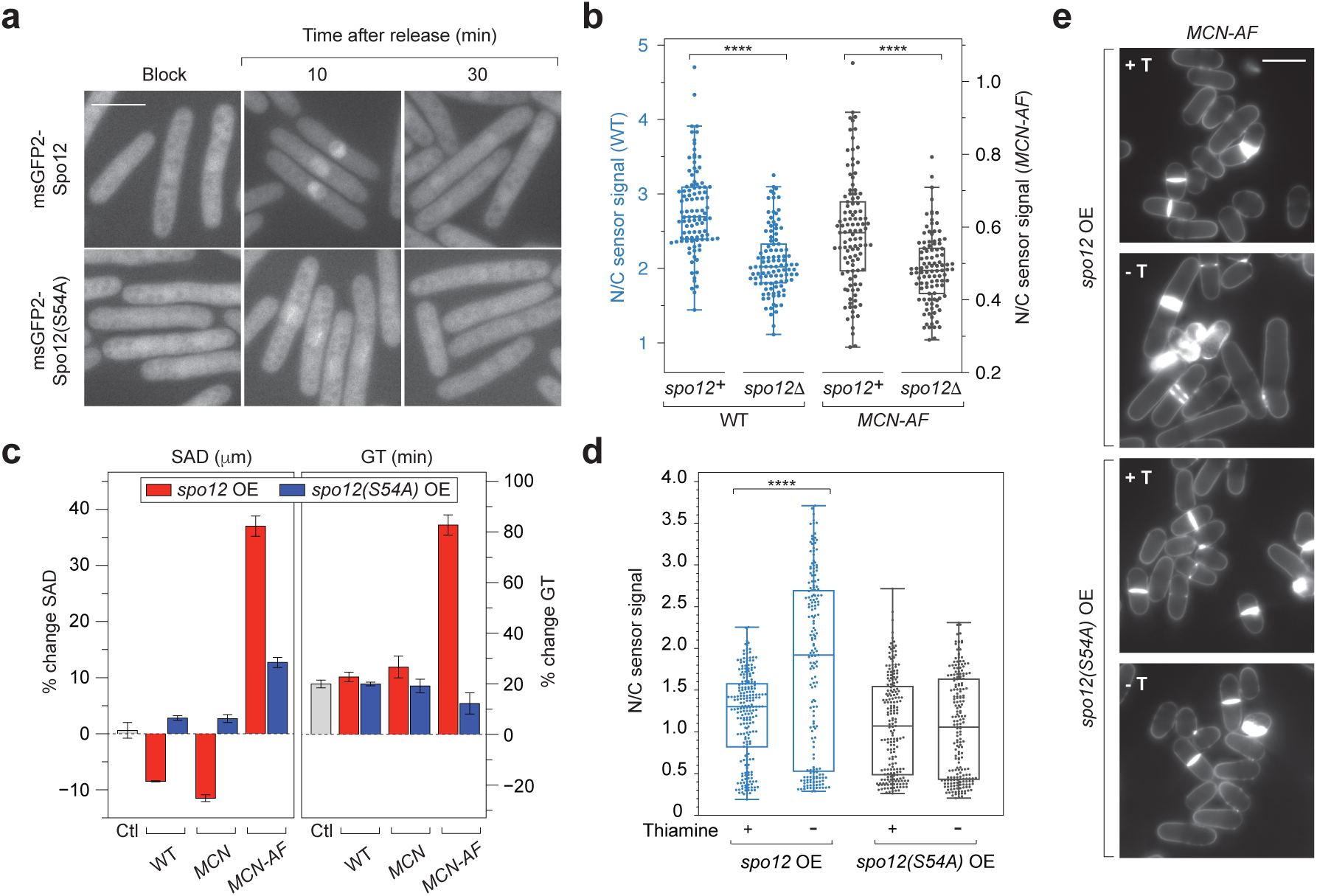
Spo12 is part of a novel feedback system on Cdc2 function. **a.** *MCN* cells expressing msGFP2-tagged versions of Spo12 or Spo12(S54A) were arrested in G2 (Block) using the 3-MBPP1 analog (Fig. 1a). Cells were then synchronously released into the cell cycle upon inhibitor wash-off (Extended data Fig. 3b-d). Images were acquired at different times after the release (full dataset in Extended data Fig. 3f). msGFP2-Spo12 accumulates in the nucleus at G2/M (T=10 min) and becomes diffuse after mitotic exit (T=30 min). This is not observed for msGFP2-Spo12(S54A), which only shows a mild and transient signal on the mitotic spindle (see Extended data Fig. 3f). Scale bar = 10 μm. **b.** Analysis of the peak nuclear/cytoplasmic signal ratio (N/C) of the synCut3-mCherry Cdk1 activity sensor^27^ in cells of the indicated genotypes. Time-lapse experiments were performed in microfluidic devices to maintain optimal growth (Extended data Fig. 4a). For each cell, the N/C ratio of mCherry intensity was determined at the time point where the maximum nuclear signal was detected. Pooled datasets of two independent replicates (*n*>50 per replicate). Box and whiskers plots indicate the min, Q1, Q3 and max, with outliers determined by 1.5 IQR (interquartile range). Note that the peak N/C ratio obtained using this specific biosensor is significantly lower in *MCN-AF* cells compared to WT, consistent with previous reports^27^. ****: *p*<0.0001 (Mann-Whitney U test for pooled datasets). **c.** Changes in cell size at division (SAD) and population generation time (GT) upon *spo12* or *spo12(S54A)* overexpression (OE, see Methods) in the indicated genetic backgrounds (see Extended data Fig. 4b, c for overexpression levels). The difference in SAD or GT in the absence of thiamine (overexpression) was determined and represented as a percentage change compared to the corresponding sample kept in the presence of thiamine (repression). Averages of three independent replicates with standard errors (*n*>100 for each replicate for SAD). Ctl: control strain expressing endogenous *spo12*. Upon *spo12* overexpression, the population doubling time of *MCN-AF* cells was 260 ± 6 min (standard error). **d.** N/C ratio of synCut3-mCherry signal in asynchronous WT cultures upon *spo12* OE or *spo12(S54A)* OE treated as in *c*. Only cells in which nuclear mCherry signal was detected were analyzed. Cells already undergoing mitosis (as identified by changes in nuclear morphology) were excluded. Pooled datasets of 2 independent replicates (*n*>85 per replicate). Box and whiskers plots indicate the min, Q1, Q3 and max, with outliers determined by 1.5 IQR (interquartile range). ****: *p*<0.0001 (Mann-Whitney U test for pooled datasets). Note that as expected, the N/C values obtained from these asynchronous cultures are lower and more spread than in panel *b*, which focuses on the maximum nuclear sensor intensity in each cell. *spo12* OE induced a G1 extension specifically in the *MCN* background (Extended data Fig. 4d), which is likely due to the excessive levels of Spo12 in these cells (Extended data Fig. 4b, c) together with the distinctive regulation of cell cycle progression in *MCN vs* WT^6,14^. *spo12* OE also led to strong defects in *MCN-AF* cells (see *c*, *e* and Extended data Fig. 4d). Given these alterations and their potential impact on the quantification and interpretation of the synCut3-mCherry signal in asynchronous *MCN* and *MCN-AF* cultures, we restricted our analyses to *spo12* OE in WT cells. **e.** Blankophor images of *MCN-AF* cells overexpressing *spo12* or *spo12(S54A)* as in *c*. +T: presence of thiamine. -T: absence of thiamine. Increased levels of Spo12 but not Spo12(S54A) lead to a broad range of defects. Scale bar = 10 μm.

At the single-cell level, *MCN-AF* cells that were maintained in optimal growth conditions within perfused microfluidic devices showed a bimodality in their cycle time distribution (Fig. 1g). In fact, a fraction of these cells displayed even shorter cycle times than wild-type cells. In contrast, in *MCN-AF* cells lacking Spo12, this distribution was unimodal, peaking at division times that were intermediate to the two subpopulations observed in *MCN-AF* (Fig. 1g). As a result, the average division time of *MCN-AF spo12Δ* cells was similar to that of *MCN-AF* and even lower than those of WT and *spo12Δ* cells (see legend of Fig. 1g). However, while a fraction of *MCN-AF* cells (∼16%; Fig. 1h) did not undergo division during our 10 h time-lapse experiments, this was substantially reduced by the loss of *spo12* (∼8%), contributing to the improved growth rate of the population. Finally, we observed that *MCN-AF* cells show increases in the length of mitosis, the residency time of the Polo kinase Plo1 at the spindle pole body (SPB) during cell division, and the duration of chromosome segregation (Extended data Fig. 2f). Only the latter was rescued by deletion of *spo12*, making it more comparable to WT and *spo12Δ* cells (Extended data Fig. 2f, iii).

Altogether, these observations demonstrate that loss of Spo12 function in cells lacking the Wee1+Cdc25 feedback loops promotes population growth and cell-to-cell homogeneity in cell cycle progression, while impacting a number of phenotypes associated with the division process. This highlighted a potential role for Spo12 in tuning the cell cycle control network.

### Spo12 modulates Cdc2 function in a Cdc2-dependent manner

Spo12 harbors two canonical Cdk target sites (S47 and S54, Fig. 1d), one of which (S54) was previously identified in a phosphoproteomic study of Cdc2 substrates in fission yeast^26^. This suggested that Spo12 function may be directly regulated by Cdk-mediated phosphorylation. Substitution of Spo12 S54 by an alanine (S54A) phenocopied the full deletion of *spo12*, with *MCN-AF spo12(S54A)* populations showing an improved growth rate, reduction of G1 duration and resistance to replication stresses similar to *MCN-AF spo12Δ* (Fig. 1e, f; Extended data Fig. 2a, b, d). This was not the result of differences in protein concentration, as Spo12 and Spo12(S54A) were detected at comparable levels in the WT, *MCN* and *MCN-AF* genetic backgrounds (Extended data Fig. 2g). Interestingly, in asynchronous cultures, we observed nuclear accumulation of msGFP2-Spo12 in long mono-nucleated cells in all tested backgrounds, while msGFP2-Spo12(S54A) signal was barely detectable in the nucleus (Extended data Fig. 3a). We then further probed the relationship between Cdc2 phosphorylation of Spo12 and its subcellular localization throughout the cell cycle. Taking advantage of the sensitivity of the Cdc13-Cdc2 fusion protein to reversible inhibition by non-hydrolysable ATP analogs^6^ (see legend of Fig. 1a), we synchronized cells expressing either *msGFP2-spo12* or *msGFP2spo12(S54A)* in G2 and subsequently released them into the cell cycle (Extended data Fig. 3b-d). First, we did not observe major differences in protein levels throughout the cell cycle in either population (Extended data Fig. 3e). Strikingly, while msGFP2-Spo12 was significantly enriched in the nucleus at the G2/M transition and became more diffuse upon mitotic exit (Fig. 2a; Extended data Fig. 3f), this was not the case for msGFP2-Spo12(S54A). This variant of Spo12 was only faintly detected at the mitotic spindle during mitosis (Extended data Fig. 3f). These results indicate that S54 phosphorylation by Cdc2 is important for the transient nuclear accumulation of Spo12 and its associated function. Moreover, the strict timing in Spo12 subcellular behavior supports a role in the mitotic process.

**Figure 3.**
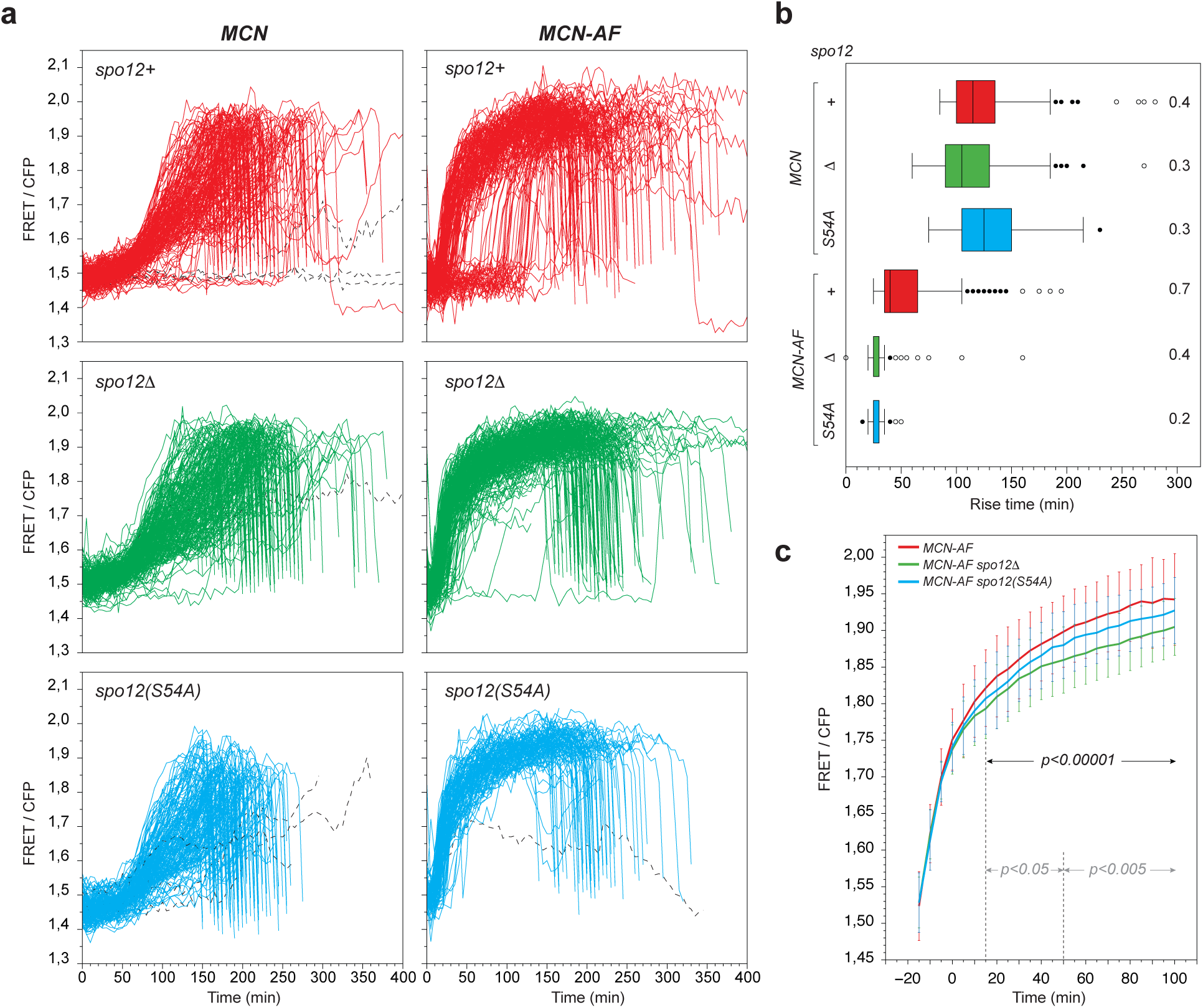
Loss of Spo12 alters Cdk activity dynamics in the absence of Wee1+Cdc25 regulation. **a.** Single-cell profiles of FRET/CFP signal for the indicated genotypes using the Eevee-spCDK biosensor^29^. All traces are aligned to the previous division for each cell (T=0 min). Analyses include all measurements, even for cells that did not undergo a full cell cycle over the duration of the time-lapse experiments. Dashed lines represent aberrant profiles. For WT (see Extended data Fig. 5a) and *MCN* cells, the profiles are as previously reported^29^. *MCN*: *n*=187; *MCN spo1211*: *n*=225; *MCN spo12(S54A)*: *n*=173; *MCN-AF*: *n*=165; *MCN-AF spo1211*: *n*=247; *MCN-AF spo12(S54A)*: *n*=128. For display purposes, the *x*-axis is restricted to the 0-400 min window. Only a few cells were measured beyond 400 min (*n*=5 in *MCN*; *n*=1 in *MCN spo1211*; *n*=3 in *MCN-AF*; *n*=4 in *MCN-AF spo1211*). Expression of *spo12(S54A)* has a similar effect on the FRET/CFP trajectories as deletion of *spo12*. **b.** Analysis of the time at which the FRET/CFP ratio rises for each cell in the profiles in *a*. The rise time corresponds to the first time point at which the smoothened FRET/CFP value (see Methods) is above a threshold of 1.65. Box and whiskers plots indicate the min, Q1, Q3 and max, with outliers determined by 1.5 IQR (interquartile range). For display purposes, the *x*-axis was restricted to the 0-300 min window, excluding 2 outliers in *MCN* (rise time > 450 min). Coefficients of variation are indicated. **c.** Comparison of the average FRET/CFP signals during the Cdk activity rise (up to 100 min after the rise time) between the indicated strains. The FRET/CFP traces in *a* were aligned to T = -15 min before the activity rise time (see *b*). At each time point, averages and standard deviations were determined. Statistical tests (Mann-Whitney U test) were performed and temporal windows of significance are shown (black: *MCN-AF vs. MCN-AF spo1211*; grey: *MCN-AF vs. MCN-AF spo12(S54A)*).

In eukaryotes, orderly progression through the key cell cycle events relies on an intricate network of feedback systems at virtually all steps of the process^13^. The effects of *spo12* deletion on *MCN-AF* cells, together with the tight Cdc2-dependent control of Spo12 nuclear localization just prior to mitosis, suggested that Spo12 might in turn regulate Cdc2. We therefore assessed peak Cdc2 activity in the presence and absence of Spo12 in both WT and *MCN-AF* cells. To this end, we used a previously established *in vivo* marker for Cdc2 activity that relies on the Cdc2-mediated accumulation of the biosensor in the nucleus^27^ (Extended data Fig. 4a). While such sensors are also affected by the function of Cdk counteracting phosphatases, their output will be referred to as “Cdk activity” for simplicity. Strikingly, in both WT and *MCN-AF*, loss of Spo12 was associated with a reduction in nuclear translocation of the marker (Fig. 2b), suggesting that Spo12 may be a positive regulator of Cdc2. Consistent with this possibility, in WT and *MCN* cells, overexpression of *spo12* but not *spo12(S54A)* led to a lower average cell size at division (Fig. 2c; Extended data Fig. 4b-d), a standard indicator of advances in mitosis in fission yeast. Using the synCut3-mCherry marker in the WT background, we demonstrated that this size phenotype was accompanied by an increase in nuclear biosensor signal, indicating increased Cdk activity (Fig. 2d). In *MCN-AF* cells, in which Cdc2 is deregulated, *spo12* overexpression brought about a significantly longer population generation time together with strong morphological and cell cycle defects (Fig. 2c, e; Extended data Fig. 4d). If Spo12 indeed regulates Cdc2, this may result from excessive Cdc2 activation that is not compensated by Wee1+Cdc25 regulation in this background. Collectively, these observations suggest that Spo12 may be part of a novel positive feedback loop on Cdc2 function.

**Figure 4.**
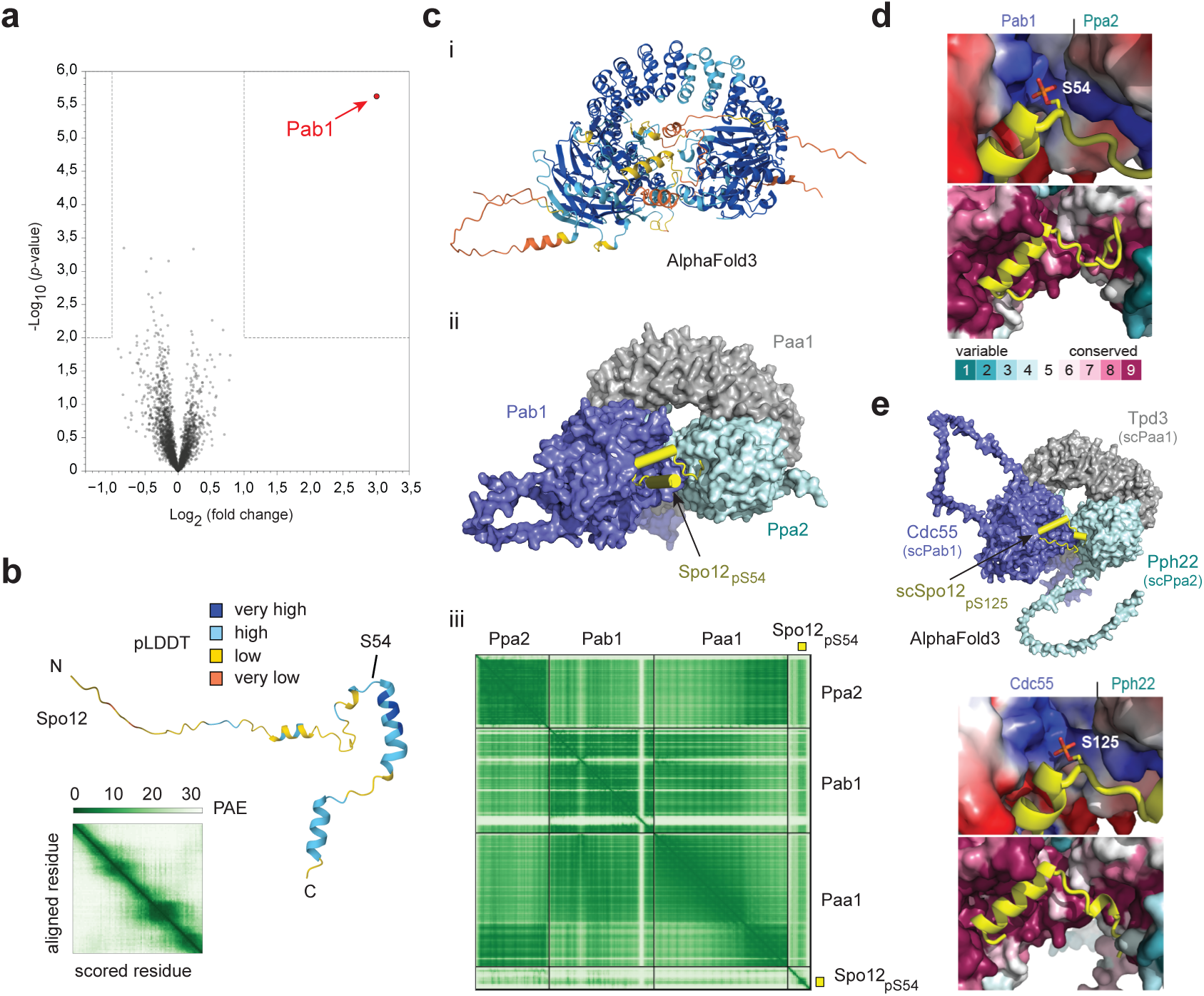
Spo12 defines a new family of PP2A inhibitors. **a.** Volcano plot of the Turbo-ID results comparing strains expressing *spo12 vs. spo12(S54A)* with Turbo-ID tags. Values are from averages of five independent replicates (Supplementary Table 2). *x*-axis: log_2_ (fold change); *y*-axis: -log_10_ (*p*-value). Proteins for which the averaged quantity of all 10 samples was lower than 25 were excluded. Among the remaining candidates, those that were not detected in up to 2 of the Spo12 samples and/or of the Spo12(S54A) samples were then additionally excluded. Using these filters, ∼96% of the proteins identified in the Turbo-ID experiment are plotted. Pab1/B55 (red arrow) was the sole significant factor interacting with Spo12 but not Spo12(S54A) that was detected. **b.** Predicted Spo12 structure from the AlphaFold Protein Database (https://alphafold.ebi.ac.uk/) for entry AF-Q10189-F1-v4. The results of the pLDDT (predicted Local Distance Difference Test) are mapped onto the predicted model and show generally poor confidence in keeping with a disordered protein, apart from two putative helical segments. The PAE (Predicted Aligned Error) plot is also consistent with a lack of intermolecular contacts, which is common for intrinsically discorded proteins. **c.** i: cartoon representation of an AlphaFold3 (https://alphafoldserver.com) model of the Spo12(pS54):Pab1:Ppa2:Paa1 complex, with pLDDT values colored as in *b*. Pab1, Ppa2 and Paa1 are the B-regulatory, C-catalytic and A-scaffolding subunits of the PP2A phosphatase complex. ii: surface representation of the complex, with Spo12(pS54) in yellow. iii: PAE plot for the complex above. Yellow rectangles correspond to the conserved region of Spo12 (dashed square in Extended data Fig. 7a). See Extended data Fig. 7 and 8 for more prediction details. **d.** *Top panel*: cartoon representation of Spo12(pS54) with predicted helices (yellow) shown on a surface representation of Pab1, Ppa2 and Paa1 (colored based on their ConSurf conservation as in Extended data Fig. 7a). *Bottom panel*: close-up view of the Spo12 conserved region around S54 (yellow, aa 43-74) with Pab1 and Ppa2 surfaces colored based on charge (from positive in blue to negative in red). **e.** *Top panel*: surface representation of an AlphaFold3 model of the *S. cerevisiae* Spo12(pS125):Cdc55/Pab1:Pph22/Ppa2:Tpd3/Paa1 complex. See Extended data Fig. 9 for more prediction details. *Middle panel*: cartoon representation of *S. cerevisiae* Spo12(pS125) on a surface representation of Cdc55/Pab1, Pph22/Ppa2 and Tpd3/Paa1 as in *d*. *Bottom panel*: close-up view of the *S. cerevisiae* Spo12 conserved region around S125 as in *d*.

### Spo12 function shapes the profile of Cdc2 activity in the absence of the mitotic switch

The distinctive temporal profile of Cdk activity throughout the division cycle of wild-type cells is thought to rely strongly on the Wee1+Cdc25 loops^13,14,28^. During G2, Cdk activity is kept low by Wee1-mediated inhibition. However, when sufficient active cyclin-Cdk complexes accumulate as a result of cell growth, Wee1 becomes inhibited while Cdc25 is activated, resulting in a Cdk activity burst that triggers mitosis. Cdk activity then returns to a low level at mitotic exit through cyclin B degradation, which is part of a negative feedback loop on Cdk function. Cells subsequently return to G1, during which specific Cdk inhibitors take charge (*e.g.*, Rum1 in fission yeast). The nature, control and significance of the temporal pattern of Cdk activity during the transition to S phase remains more elusive. To assess accurately the impact of Spo12 on the dynamics of Cdc2 function, we used the recently-developed Eevee-spCDK biosensor^29^. This system takes advantage of intra-molecular Förster Resonance Energy Transfer (FRET) upon Cdc2-dependent phosphorylation rather than nuclear translocation of a modified substrate, allowing us to monitor changes in Cdc2 activity with higher temporal resolution.

The FRET profiles that we obtained in WT and *MCN* cells were as previously reported for fission yeast cells expressing Eevee-spCDK^29^ (Fig. 3a, Extended data Fig. 5a). Loss of Spo12 in these backgrounds did not bring about noticeable changes, consistent with the absence of detectable phenotypes in these cells (Fig. 3a, Extended data Fig. 5a). In contrast, our FRET biosensor revealed unanticipated temporal patterns of Cdc2 activity in *MCN-AF* cells (Fig. 3a, b). First, the timing of Cdc2 activation was highly heterogenous between cells. Around 50% of the population showed a rapid increase in Cdk activity shortly after the preceding mitosis, much earlier than in WT and *MCN* cells, while the other half exhibited a long G1 phase with low Cdk activity. Second, in all cells, the activity rise was strikingly abrupt, even more than what we observed in the WT and *MCN* backgrounds, which have a fully functional Wee1+Cdc25-mediated mitotic switch. Finally, Cdc2 activity remained high for extended periods of time before cells entered mitosis to eventually drop at mitotic exit. The reason why *MCN-AF* cells do not enter mitosis rapidly after the switch to high Cdc2 activity remains unclear, but it may be linked to the observed slowdown in the increase in FRET/CFP signal after the initial burst (from T≈20 min in Fig. 3c), with Cdk activity reaching a potentially higher mitotic threshold only at later time points.

**Figure 5.**
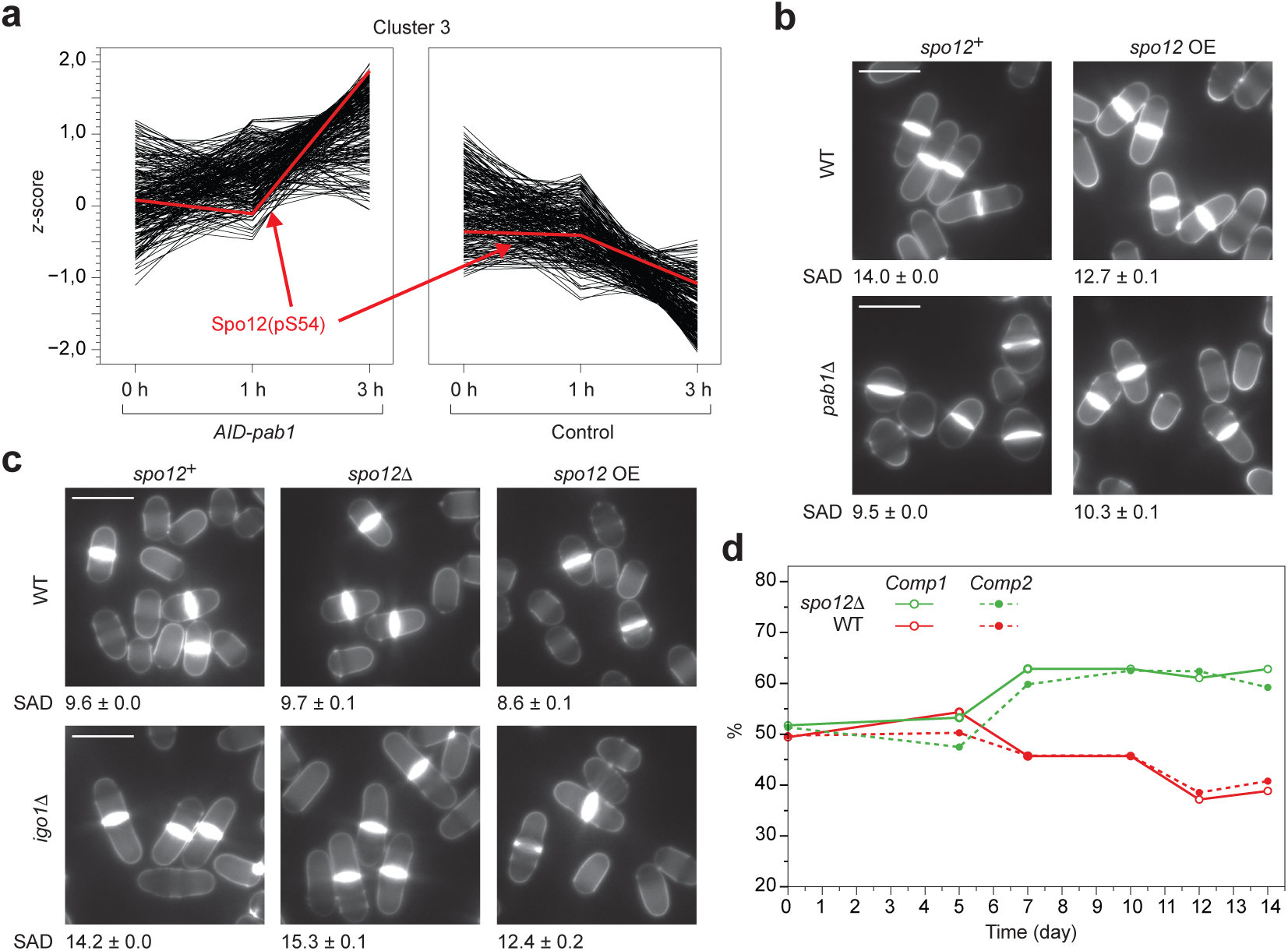
Functional analysis of the regulation of PP2A by Spo12. **a.** Loss of Pab1 leads to an increase in Spo12 S54 phosphorylation. Kinetic profiles for Cluster 3 from phosphoproteomic analysis upon Pab1 depletion (Extended data Fig. 6c-g). Individual phosphopeptides are shown as black lines, with Spo12(pS54) highlighted in red. The *x*-axis represents time (h) after thiamine + NAA treatment in control or *AID-pab1* strains. **b.** Blankophor images and cell size at division (SAD) for WT or *pab1*11 cells overexpressing *spo12* (*spo12* OE). Averages of three independent replicates with standard errors (*n*>100 for each replicate). **c.** Blankophor images and SAD upon nitrogen depletion for WT or *igo1*11 cells harboring the indicated *spo12* alleles (OE: overexpression). Cells were grown in the absence of thiamine for 48 h. Cultures were then washed by filtration using minimal medium without a nitrogen source and kept in these conditions for 2 h; SAD was then determined. Averages of three independent replicates with standard errors (*n*>80 for each replicate). For *b* and *c*, scale bar = 10 μm. **d.** Competition experiments between WT and *spo1211* cells in an eVOLVER setup^48,49^ (see Methods). Both strains carried a deletion of the flocculin-encoding gene *gsf2*, a critical alteration for the continuous culture system^48^. In the WT background, *gsf2* was deleted using a hygromycin resistance cassette. In the *spo1211* background, both *spo12* and *gsf2* were deleted using a kanamycin resistance cassette. At each time point, a defined number of cells was plated on rich medium followed by replica plating on medium containing either kanamycin or hygromycin. The percentages of cells of each genotype were then determined relative to the total number of colonies growing on the non-selective rich medium. Two independent experiments are shown (Comp1 and Comp2).

In line with our assays using the synCut3-mCherry sensor (Fig. 2b), loss of Spo12 function through either *spo12* deletion or S54A substitution lowered the plateau level of Cdc2 activity (Fig. 3c). Furthermore, the heterogeneity in Cdc2 activity profiles was significantly reduced in both cases (Fig. 3a, b), reminiscent of the single-cell cycle time measurements in *MCN-AF spo12Δ vs. MCN-AF* (Fig. 1g). The improved cell-to-cell homogeneity was noticeable even when restricting our analyses to the most rapidly proliferating cells (*i.e.*, cells that underwent a full cell cycle within the time frame of the FRET time-lapse experiments, Extended data Fig. 5b). These phenotypes may contribute to the observed differences in the fraction of non-dividing cells between the two genetic backgrounds (Fig. 1h). Altogether, our results reveal an unexpected temporal pattern of Cdc2 activity in cells lacking the mitotic switch and suggest that in this context, modulation of Cdc2 function through alteration of Spo12 has a strong impact on cell cycle dynamics.

### Spo12 defines a novel family of negative regulators of the PP2A phosphatase

While Spo12 may be part of a direct positive feedback loop that activates Cdc2 in a Cdc2-dependent manner, it may also operate through modulating negative regulators of Cdc2. Alternatively, Spo12 may control factors that do not directly target Cdc2 activity but rather affect the phosphorylation state of Cdc2 substrates. Indeed, as mentioned earlier, *in vivo* dephosphorylation of the biosensors certainly plays a role in the measured signals, which may for instance reflect changes in activity of Cdk1-counteracting phosphatases. To gain insight into the mechanisms underlying Spo12 function, we used a Turbo-ID (TID) approach to identify proteins that interact with or are located in close proximity to Spo12^30^. For this purpose, we compared cells expressing *3HA-TID-spo12 vs. 3HA-TID-spo12(S54A)* alleles integrated at the endogenous *spo12* locus (Extended data Fig. 6a, b). This strategy identified Pab1/B55, a regulatory subunit of the conserved Cdc2-counteracting phosphatase complex PP2A^31^, as the sole factor that was significantly and specifically enriched in cells expressing the TID-tagged wild-type Spo12 (Fig. 4a; Supplementary Table 2). This interaction was more recently confirmed in a proteomic study for Pab1-binding partners^32^. Collectively, this suggested that Spo12 may not be a direct regulator of Cdc2 activity but rather promote Cdc2 function and modulate cell cycle progression through stoichiometric inhibition of the B55:PP2A complex.

**Figure 6.**
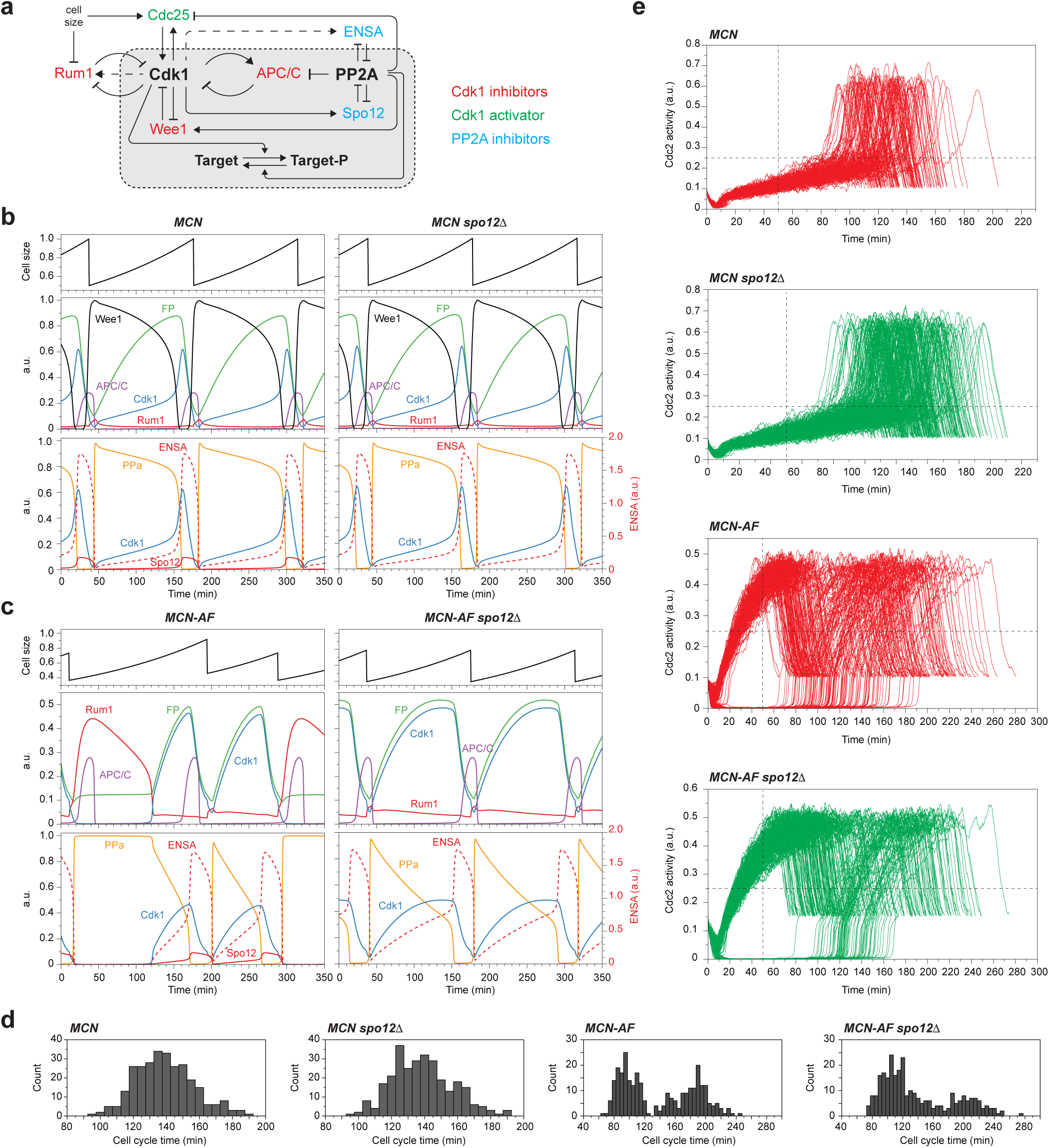
Mathematical modeling of a cell cycle network integrating Spo12 function. **a.** Model network of the regulation of Cdk1 and PP2A in cell cycle progression. Spo12 defines a novel family of PP2A inhibitors that are directly regulated by Cdk1. Spo12 operates at both mitotic entry and exit and is involved in an intricate network of feedback loops on Cdk1 function. This network was used as the basis for mathematical modeling of the impact of Spo12 alteration in cell cycle progression. **b, c.** Deterministic simulations using the model in *a* (see Supplementary Model, part A) for the indicated genotypes. FP: concentration of the MCN or MCN-AF fusion protein; Cdk1: active form of the Cdk1 kinase (dephosphorylated on T14 and Y15); Rum1: concentration of Rum1, a key inhibitor of Cdc2 in G1; Wee1: active form of the Wee1 kinase (dephosphorylated); APC/C: active form of the APC/C (Anaphase Promoting Complex; phosphorylated by Cdc2); Spo12: active form of Spo12 (phosphorylated by Cdc2); PPa: active form of the PP2A phosphatase; ENSA: active form of the PP2A inhibitor ENSA (Igo1 in fission yeast). Values for cell size are relative to the size at division of *MCN* cells (1 a.u., arbitrary unit). **d.** Distribution of cell cycle time for the indicated genotypes using stochastic simulations of the model in *a* (see Supplementary Model, part C and Extended data Fig. 11a, b; *n*>100 for each genotype). The *in vivo* behaviors of the different genotypes are recapitulated by the model, including the bimodal distribution of cell cycle time in *MCN-AF* cells and its partial rescue in the absence of Spo12 (Fig. 1g). Average and standard deviation for cell cycle time: *MCN*, 139 ± 18 min; *MCN spo1211*, 139 ± 18 min; *MCN-AF*, 139 ± 47 min; *MCN-AF spo1211*, 139 ± 47 min. The accompanying distributions for cell size at division are shown in Extended data Fig. 11c. **e.** Simulated single-cell Cdk activity traces for the indicated genotypes based on the stochastic simulations of the model (see *a*) as in Extended data Fig. 11a, b. These plots reflect the FRET/CFP traces in Fig. 3a. Traces are aligned to the previous division for each cell. For each genotype, *n*>200. The dashed lines indicate the thresholds in both Cdc2 activity (0.25) and Time (50 min) that were used to distinguish between cells with an early *vs.* late rise in Cdc2 activity (see Extended data Fig. 11d). a.u.: arbitrary unit. Also see Supplementary Model, part C.

To evaluate this hypothesis, Spo12 structure and interactions were analyzed *in silico*. Fission yeast Spo12 is predicted to be conformationally disordered (Fig. 4b; Extended data Fig. 7), with two noticeable potential helical regions located within the most conserved area of the protein, in the vicinity of the S54 phosphorylation site. Importantly, an AlphaFold3 model predicted with high confidence the formation of a large protein complex consisting of the heterotrimeric (ABC) PP2A assembly Pab1:Ppa2:Paa1 with phosphorylated Spo12(pS54) (Fig. 4c). The greatest confidence score in these analyses suggests a docking of the conserved region of Spo12 between Pab1 and Ppa2 (Fig. 4c). Consistent with our experimental results, the absence of Cdc2-dependent phosphorylation of Spo12 on S54 significantly reduced the confidence of this complex (Extended data Fig. 8a). The role of S54 phosphorylation in Spo12 function was also supported by the association of this modified serine with a large basic patch formed by the neighboring surfaces of Pab1 and Ppa2 (Fig. 4d).

While Spo12 shares no specific sequence similarity with known stoichiometric inhibitors of PP2A in fission yeast or in more complex eukaryotes (*e.g.*, Igo1, ENSA, ARPP19, FAM122A), its structure and mode of interaction with the PP2A complex is reminiscent of those described from cryo-EM approaches for some of these factors (Extended data Fig. 8b-f)^33^. However, the SPO12 domain (Fig. 1d) is relatively well-conserved between fission and budding yeast (Extended data Fig. 9a, b). Moreover, S118 and S125, which are located within this domain of *S. cerevisiae* Spo12, are both consensus Cdk target sites and are essential for Spo12 function^19^. Interestingly, AlphaFold3 predicted with high confidence the formation of a complex between *S. cerevisiae* PP2A and Spo12 phosphorylated on S125 (pS125) (Fig. 4e; Extended data Fig. 9c-e), analogous to the *S. pombe* complex. Furthermore, pS125 is strikingly located against the same charged regions of scPab1/Cdc55 and scPpa2/Pph22, as was predicted for pS54 in fission yeast (compare Fig. 4d with 4e). In addition, the other putative Cdk target residue of Spo12 (S47 in fission yeast and S118 in budding yeast) appears to be directly situated in the active site of the PP2A complex and is exactly 7 amino acids away, N-terminal to the primary Cdk target in both species (S54 in *S. pombe* and S125 in *S. cerevisiae*; Extended data Fig. 10). This reinforces the idea that at the molecular level, Spo12 functions similarly in fission and budding yeast. In both organisms, double phosphorylation of Spo12 increased the confidence of the predicted complex with PP2A (Extended data Fig. 10b, d), suggesting that phosphorylation of Spo12 S47 and S118 in fission and budding yeast, respectively, may contribute to Spo12 binding. Finally, scSpo12 was previously shown to also harbor a LxF cyclin docking motif for Cdk1 substrates^34^, providing further evidence that Cdk1 directly targets Spo12.

The structural prediction of a Spo12(pS54):Pab1:Ppa2:Paa1 complex and the positioning of the conserved area of Spo12 in the PP2A active site, similarly to other PP2A inhibitors, supports the hypothesis that Spo12 positively promotes Cdc2 function through stoichiometric inhibition of PP2A activity. As such, Spo12 may also be a target of PP2A. To test this possibility, phosphoproteomic analyses were performed using cells in which degradation of Pab1 was induced by an auxin-degron system^35^, together with its transcriptional silencing by means of a thiamine-repressible promoter^36^ (Extended data Fig. 6c). This approach identified a complete set of potential B55:PP2A targets in fission yeast, which was subdivided in four distinct clusters based on the dynamic changes in their phosphorylation status upon loss of Pab1 (Extended data Fig. 6d, e; Supplementary Table 3). Notably, Spo12 S54 was detected in cluster 3, showing a significant increase in its phosphorylation state in the absence of Pab1 (Fig. 5a; Extended data Fig. 6e, f). In contrast, Spo12 S47 phosphorylation did not change (Extended data Fig. 6e). However, its central position within the PP2A active site (Extended data Fig. 10a, c) indicates that it may be very efficiently dephosphorylated and thus difficult to detect.

We then tested *in vivo* the function of Spo12 in cell cycle control as an inhibitor of the PP2A complex. First, the reduction in cell size at division induced by *spo12* overexpression in WT was fully suppressed by deletion of *pab1*, despite the overall smaller size of the *pab1Δ* cells (Fig. 5b). Next, as Spo12 and the ENSA/ARPP19 counterpart Igo1^37^ both bind the catalytic pocket of PP2A (Extended data Fig. 8), we surmised that alteration of Spo12 levels may modulate the well-characterized behavior of *igo1Δ* cells in nutrient-limited conditions^37^. Consistent with this idea, loss of Spo12 enhanced the size phenotype of *igo1Δ* cells deprived of nitrogen. Conversely, overexpression of *spo12* partially rescued the response of *igo1Δ* to this nutritional stress (Fig. 5c).

Altogether, these results demonstrate that Spo12 functions in the cell cycle regulatory network through binding and inhibiting the B55:PP2A complex. Importantly, Spo12 defines a novel, distinct and conserved family of PP2A inhibitors. First, in contrast to Igo1, ENSA or AARP19, which are targets of the Greatwall kinase^38–40^, our results and previous reports in budding yeast^19^ demonstrate that Spo12 is itself positively regulated by Cdk-dependent phosphorylation, representing a direct feedback system on Cdc2 function. Second, while previous evidence suggested that the cellular role of fission yeast Spo12 differs from that of budding yeast Spo12^23^, our structural predictions indicate that the two proteins act similarly at the mechanistic level. Not only does this highlight the conservation of the Spo12-dependent loop, but it also elucidates the way in which Spo12 contributes to the FEAR network in *S. cerevisiae*, which remained an open question.

Finally, although loss of *spo12* appeared to have no detectable impact on the growth of WT and *MCN* populations in laboratory conditions (Fig. 1e), we found that it surprisingly provides a proliferative advantage over WT in competition experiments, which are highly sensitive to small differences in generation times (Fig. 5d). This suggests that the Spo12-dependent regulation of PP2A is a critical process that has evolved despite being associated with a trade-off on population growth.

### Dynamic role of the new Spo12 feedback on Cdc2 function at mitotic entry and exit

Our experimental and structural data suggest a model in which Spo12 - via stoichiometric inhibition of PP2A - acts at both mitotic entry and exit (Fig. 6a), impacting the phosphorylation state of Cdk substrates, including critical cell cycle regulators. In this simple regulatory system, Spo12 and Igo1/ENSA participate in several feedback and feed-forward loops on Cdc2 function. To evaluate whether the dynamics of this control network could recapitulate our experimental observations and shed light on unexpected properties of different cell cycle circuits, we generated a mathematical model based on the proposed wiring diagram.

First, we examined a deterministic model based on a set of non-linear ordinary differential equations (ODEs, see Supplementary Model, part A). This explicitly includes cell growth, which, for simplicity, is assumed to be exponential. This is an essential element of the approach, as the increase in size is critical for driving changes in concentration of the key regulators of the cell cycle, in particular those of the MCN and MCN-AF fusion proteins. Note that the model describes changes in Cdk1 activity *per se*, while our experimental results using biosensors integrate the inputs from both Cdk1 and its counteracting phosphatases. This may underlie potential quantitative differences between model behavior and *in vivo* data. Deterministic simulations predicted stable cell cycles in *MCN* and *MCN spo12Δ* cells, with similar cycle times and cell size at division (Fig. 6b). Significantly, the behavior of the network in the *MCN-AF* background recapitulated the surprisingly abrupt and early post-mitotic increase in Cdc2 activity, as well as the extended period of high activity compared to *MCN* cells (Fig. 6c). The model also predicted that the absence of Cdc2 regulation by Wee1+Cdc25 in *MCN-AF* cells leads to variability in cell cycle time, size at division and G1 duration, with the occurrence of long G1 phases associated with persistently high levels of Rum1, the G1 inhibitor of Cdc2 (*e.g.* compare the two successive cycles in Fig. 6c). Loss of Spo12 in the *MCN-AF* context was sufficient to reduce this overall heterogeneity in model simulations and bring about more regular profiles of Cdc2 activity from one cycle to the next, thus mirroring experimental observations (Fig. 6c).

Next, we performed stochastic simulations by introducing noise terms to the ODEs describing the accumulation of the cyclin B-Cdk1 fusion protein and of Rum1 (Extended data Fig. 11a, b). The distributions of interdivision time and cell size at division that were derived from these simulations (Fig. 6d, Extended data Fig. 11c) were strikingly similar to those obtained from experimental measurements (Fig. 1g, Extended data Fig. 2b). In particular, the bimodal distribution of cell cycle duration in *MCN-AF* cells, which results from the alternation of long and short division cycles in the model (Extended data Fig. 11b) and is reminiscent of the quantized cell cycles in *wee1-50^ts^ cdc2511* cells^41^, was also predicted to decrease in *MCN-AF spo12Δ* (compare Fig. 1g with Fig. 6d). Moreover, single-cell trajectories of Cdc2 activity computed from the stochastic simulations (Fig. 6e) also recapitulated the experimental FRET data (Fig. 3a). Notably, the fraction of cells with a long G1 and late rise in Cdc2 activity was significantly reduced in the *MCN-AF spo12Δ* simulations compared to *MCN-AF*, although not to the same extent as the *in vivo* results (Fig. 6e, Extended data Fig. 11d). Even among cells with a long G1 phase, loss of Spo12 resulted in a more homogenous timing of Cdc2 activation (Extended data Fig. 11e).

Consistent with the single-cell experimental data using the *MCN-AF* and WT genetic backgrounds, the average cycle times of *MCN-AF* and *MCN-AF spo12Δ* cells obtained from the stochastic simulations were comparable to that of *MCN* (see legends of Fig. 1g and Fig. 6d, Supplementary Table 4). However, these experiments and simulations restrict the analyses to proliferating cells. Cells that do not divide in the *MCN-AF* and *MCN-AF spo12Δ* backgrounds (Fig. 1h), which contribute to the differences in population generation time between the different strains (Fig. 1e), are excluded. To account for this, we mathematically derived the number of viable dividing cells while integrating a probability of cell cycle arrest (Supplementary Model, part C). This predicted generation times and fractions of non-dividing cells that are well in line with our experimental observations (Supplementary Model, part C and Fig. 1e, h). The model proposed in Fig. 6a therefore also recapitulates the occurrence of cell cycle exit in the different tested strains.

To understand why certain genotypes exhibit a significant probability that cells terminally exit the cell cycle, we explored the dynamics of the mutant cell cycle control networks using bifurcation diagrams, which illustrate how the different dynamic states of the control circuit depend on increasing size of growing cells (Extended data Fig. 12, Supplementary Model, part B). These diagrams identified a non-canonical steady state in large cells of some of the genetic backgrounds. This state corresponds to a cell cycle arrest that occurs when cells fail to fully inactivate APC/C at the end of mitosis, resulting in an aborted cycle with partially active APC/C and low — but not negligible — Cdc2 activity. The onset of this state of permanent arrest in *MCN-AF* appears at a more “physiological” size that is 9-fold smaller than what is observed in *MCN* (Extended data Fig. 12a, b). Moreover, in contrast to *MCN*, the stable limit cycles disappear abruptly in large *MCN-AF* cells. This predicts that in the absence of the mitotic switch, cells that show alterations in their cell cycle and grow larger will be attracted to the ‘cell-cycle block’ state. Loss of Spo12 in the *MCN-AF* background leads to a shift of the arrested state to even larger cell size and an extension of the range of the stable limit cycles. This implies a reduction in the probability of cell cycle arrest and accounts for the improved population growth of *MCN-AF spo12Δ* cells. These conclusions are confirmed in additional stochastic simulations (Supplementary Model, part C).

Finally, we probed the effects of *spo12* overexpression in the model. Simulations revealed that the proposed network (Fig. 6a) predicts a decrease in cell size at division in *MCN spo12* OE (Extended data Fig. 13a-d, Supplementary Table 4), consistent with our experimental data (Fig. 2c). Interestingly, as suggested by previous studies, the model also indicates that ectopic inhibition of PP2A, here achieved through *spo12* overexpression, resets the mitotic entry threshold to lower Cdk activity^42^ (compare Extended data Fig. 13 with Fig. 6b, c, Extended data Fig. 11a, b and Extended data Fig. 12). Taking into account the differences in mitotic entry thresholds and in Cdk activity between *MCN* and *MCN-AF* cells overexpressing *spo12* (Extended data Fig. 13d), the model also predicts that excess Spo12 is more deleterious in *MCN-AF* than in *MCN* cells (Supplementary Model, part C).

Collectively, the mathematical model brings valuable insight into the role of the Wee1+Cdc25 feedback loops in modulating the temporal profile of Cdk1 activity throughout the cell cycle. It demonstrates that a simple cell cycle network with a limited number of key switches is sufficient to account for the unexpected dynamics of substrate phosphorylation in cells lacking this critical regulation. Importantly, the remarkable similarities between model simulations and virtually all of the experimental data establish that the Spo12 loop, in which Cdc2 directly phosphorylates and activates the PP2A-inhibitor Spo12, is an integral component of the control of Cdk function at both mitotic entry and exit.

## Discussion

Eukaryotic cell proliferation is governed by a complex network of interacting control systems and feedback loops that modulates the phosphorylation dynamics of key substrates at different stages of the cell division cycle. This complexity has made it difficult to distinguish between central components of the network that are necessary and sufficient to drive the core division cycle and secondary regulation that may shape cell cycle progression in different environments. Furthermore, our current understanding of cell proliferation is largely derived from studies in standard laboratory conditions, which may have obscured regulatory systems that are critical in natural environments. We set out to explore unknown mechanisms that control cell proliferation and gain new insight into the way the cell cycle circuit can evolve. To this end, we combined laboratory evolution assays with an engineered simplified cell cycle network in fission yeast^6^. This approach allowed us to identify Spo12 as a novel stoichiometric inhibitor of the Cdk1-counteracting phosphatase PP2A. In contrast to other known regulators of PP2A activity such as Igo1/ARPP19/ENSA, Spo12 is directly regulated by Cdk1. Thus, we demonstrate that Spo12 is at the center of a novel feedback system on Cdk1 function and defines a new class of PP2A inhibitors. It is tempting to speculate that the Spo12 loop may have evolved in the context of a simpler cell cycle circuit, with the stress-sensitive Igo1 pathway emerging as a more complex mechanism that contributes to the response of cells to alteration of their environment.

Interestingly, our findings indicate that this new cell cycle regulatory element is conserved. First, although key aspects of cell cycle control differ between budding and fission yeast, our analyses strongly suggest that *S. cerevisiae* Spo12 (scSpo12) functions similarly to *S. pombe* Spo12 at the mechanistic level. In budding yeast, both scPP2A localization and activity were proposed to be modulated by the Greatwall-ENSA orthologues Rim15-Igo1/2^43,44^. Moreover, scPP2A activity becomes downregulated at anaphase onset by separase and the FEAR network, of which scSpo12 is a member^19,20,22^. Our results and structure predictions using budding yeast proteins indicate that the molecular mechanism underlying FEAR function in the *S. cerevisiae* cell cycle, which remained elusive, is linked to the function of scSpo12 as a stoichiometric inhibitor of PP2A. Second, in more complex eukaryotes, the role and regulation of the recently-described PP2A inhibitor FAM122A do not overlap with ENSA/ARPP19, and the contribution of FAM122A to cell cycle progression is only beginning to be understood^45–47^. FAM122A is not a substrate of Greatwall^45^ and harbors Cdk1 consensus sites. While alanine substitution of all serines and threonines did not alter FAM122A function at high ectopic doses in Xenopus egg extracts^45^, it may be directly regulated by Cdk1-mediated phosphorylation *in vivo*. Thus, FAM122A may belong to the ‘Spo12 family’ of PP2A regulators, despite some differences in the predicted structure of the complexe that it forms with PP2A (Extended data Fig. 8). Notably, as intrinsically disordered proteins such as Spo12 are fast-evolving at the sequence level, making it difficult to identify functional homologs by sequence comparison, this family may be more broadly found among eukaryotic organisms.

Our approach using simplified cells and experimental evolution assays also provides us with a unique entry to understanding how cell proliferation might have evolved. Perhaps surprisingly, the improvement in population growth of cells lacking the Wee1+Cdc25 mitotic feedback loops in our study was not brought about by rewiring of existing functions or the emergence of novel mechanisms. It rather occurred by further reduction of network complexity via the loss of a feedback system on Cdk1 function. This suggests that an effective and simple strategy for promoting proliferation in the context of evolutionary processes may be to fine-tune the balance between the activities of existing core cell cycle regulatory pathways. In the case of the *MCN-AF* cells, loss of the Spo12 loop was central to compensating for the perturbation in the timely modulation of cell cycle substrate phosphorylation dynamics due to the absence of the Wee1+Cdc25 feedbacks. Complexity in cell cycle regulation may therefore have emerged in part through incremental changes in coordinated mechanisms that did not drastically unbalance the equilibrium among cell cycle control pathways. Along these lines, the loss of pre-existing regulatory elements may also have played a role in the evolution of the cell cycle network.

Finally, the functional advantages provided by complexity in the control of cell proliferation may also come with trade-offs, and the balancing of such trade-offs may have contributed to the cell cycle network topologies of present-day cells and how they have emerged. While cells lacking *spo12* do not display detectable phenotypes in batch cultures, they surprisingly outcompete wild-type cells in co-cultures, even in optimal laboratory conditions (Fig. 5d). Thus, although the acquisition of the Spo12 feedback system may have been beneficial during fission yeast evolution, it appears to come at a cost for population growth. This further underscores the intricate balance between the multiple layers that modulate cell cycle progression: each individual element of the system may not by itself promote cell proliferation. Deciphering the fine equilibrium between the different branches of the cell cycle circuit and their respective trade-offs will be central to understanding the histories and structures of eukaryotic cell cycle networks.

## Supporting information

Supplementary Model

## Acknowledgement

We would like to thank Daniel Jeffares (University of York, UK) for his help with the analysis of the whole genome sequencing data, and Helen Flynn from the Crick Proteomics Science Technology Platform (London, UK) for her help with the proteomics analyses described in Fig. 5a and Extended data Fig. 6c-g. We would like to thank Andrea Ciliberto (IFOM, Milan, Italy) for critically reading the manuscript. This work is dedicated to our collaborator and friend Béla Novák.

JCR was supported by a PhD fellowship from the University of Rennes 1 (Ministère de l’Enseignement Supérieur et de la Recherche) and a 4^th^ year PhD grant from the Ligue contre le Cancer. BL was funded by a research grant from the Région Nouvelle Aquitaine to DC (program CHESS, grant agreements 15963520 and 15964420). AJ was supported by a grant from the Agence Nationale de la Recherche (PRC eVOLve, ANR-18-CD13-0009) to DC, a research grant from the Région Nouvelle Aquitaine to DC (program CHESS, grant agreements 15963520 and 15964420) and by the French Government in the framework of the France 2030 Programme IdEx Université de Bordeaux. NC and MRDS were supported by an ERC starting grant to DC (SyntheCycle, Grant Agreement no. 310849). MSK was supported by a PhD grant and an “Internacionalization support” grant from the University of Oslo. LT and FU were supported by the Francis Crick Institute, which receives its core funding from Cancer Research UK, the UKRI Medical Research Council and the Wellcome Trust (cc2137). JY and GL were supported by a grant from the Agence Nationale de la Recherche (PRC eVOLve, ANR-18-CD13-0009). JY also received funding from the National Natural Science Foundation of China (32470663). HS was supported by JSPS KAKENHI grants (nos. 21J01354 and 22K15115) and the Dr. Yoshifumi Jigami Memorial Fund, The Society of Yeast Scientists. YG was supported by JSPS KAKENHI grants (no. 22K15110) and JST ACT-X (no. JPMJAX22B8). KA was supported by JSPS KAKENHI Grants JP24H01416, Joint Research of the Exploratory Research Center on Life and Living Systems (ExCELLS) (ExCELLS program No. 25EXC603). PYJW was supported by grants from the Région Nouvelle Aquitaine (program CHESS, grant agreements 15963520 and 15964420). This work was also supported by research grants from the Ligue contre le Cancer (comités départementaux Gironde and Dordogne) to DC.

## Author contributions

NC and MRDS performed the experimental evolution assay and the initial characterization of the evolved populations. JCR, BL and AJ performed all other experiments, except for the phosphoproteomic analyses upon loss of Pab1, which were performed and analyzed by MSK, NCB, SLA, LT and FU, and the FRET experiments, which were performed by HS, YG and KA. TA and GC designed, optimized and characterized the microfluidic system for single-cell studies and produced the mold for the microsystems. JXY, LY and GL analyzed the whole genome sequencing data from the evolution experiment. VC and GC performed the mass spectrometry analysis for the TurboID assay. PYJW contributed to the design of the project and experiments and to the analysis of the genomic and Cdk activity data. CMK performed the *in silico* structural studies. JT and BN generated the mathematical model and performed the simulations and associated analyses. DC designed and supervised the project, analyzed the data and wrote the manuscript.

