## Supplementary material for "Regulation of cell proliferation by a novel feedback system on Cdk function": Methods

### Fission yeast strains and methods

Standard methods and media were used[1]. All strains used in this study are detailed in Supplementary Table 6. The *MCN*-derived backgrounds were re-engineered compared to the initial strains[2], with the fusion modules being integrated at the endogenous *cdc13* locus. The S54A substitution and N-terminal tagging (3HA, msGFP2, TurboID) of Spo12 were generated by CRISPR using the SpEDIT system[3]. All other modifications and deletions of the open reading frames were obtained by homologous recombination. The synCut3::mCherry and Eevee-spCDK biosensors were as described[4,5]. All experiments in Fig. 1 (except panel *c*), 2, 5 and Extended data Fig. 2 (except panel *c*), 3, 4 and 6 were carried out in supplemented minimal medium (EMM6S) at 32 ºC. The laboratory evolution assays were performed in a ministat setup[6] in EMM6S + 3% galactose at 28 ºC (Extended data Fig. 1 and Fig. 1c). Initial characterization of the evolved populations and clones was carried out in the ministat conditions. The assessment of meiotic progression (Extended data Fig. 2c) was performed at 34 ºC (restrictive temperature for the *pat1-114* allele). The FRET experiments (Fig. 3 and Extended data Fig. 5**)** were carried out in YE supplemented with adenine (225 mg/L) at 30 ºC. The drop assay in Extended data Fig. 2d was performed on rich medium (YE4S) supplemented with hydroxyurea or camptothecin at the indicated concentrations. To synchronize *MCN* cells in G2 (Fig. 2a and Extended data Fig. 3b-f), the 3-MBPP1 non-hydrolysable ATP analog (Toronto Research Chemicals, Canada) was dissolved in DMSO at a stock concentration of 10 mM and added to liquid cultures at a final concentration of 1 µM. Assessments of the percentage of binucleated cells (Extended data Fig. 3c) and of meiotic progression (Extended data Fig. 2c) were carried out using heat fixed samples on microscope slides (70 ºC for 5 min) stained with DAPI (1 µg/mL).

For analysis of *spo12* overexpression in Fig. 2c-e and Extended data Fig. 4b-d, wild-type and mutant alleles were placed under the control of the thiamine-repressible *nmt1* promoter. Cells were grown in minimal medium in the presence of 60 µM thiamine for 36 h. Half of each culture was then washed three times in thiamine-free medium, while the other half was washed in medium containing 60 µM thiamine. Cells were then allowed to grow for 36 h in the corresponding medium prior to characterization.

### Laboratory evolution

#### *Ministat experiments*

Experiments were performed using the ministat setup in ~25 mL cultures as previously described[6]. Specific conditions (EMM6S + 3% glycerol at 28 °C; deletion of the flocculin-encoding gene *gsf2*) were used to prevent flocculation and biofilm formation, making it possible to keep cells exponentially growing over long periods of time[6]. Cultures were sampled and frozen during the course of the experiment. All initial characterizations (Extended data Fig. 1 and Fig. 1c) were performed in the ministat conditions. For whole genome sequencing, batch cultures were grown from frozen samples, and genomic DNA was purified using the QIAGEN pureGene Yeast/Bact kit B.

#### *Whole genome sequencing: read mapping and variant calling*

For whole genome sequencing of evolved populations and clones, genomic DNA was sequenced using Illumina Hiseq2500 PE150. Read mapping and variant calling were performed using an early development version of the Varathon pipeline (https://github.com/yjx1217/Varathon). Briefly, the raw sequencing reads were processed by Trimmomatic[7] (v0.33; options: ILLUMINACLIP:adapters.fa:2:30:10 SLIDINGWINDOW:5:20 MINLEN:36) to trim off sequencing adaptors and regions with low sequencing quality. The trimmed reads were mapped to the *S. pombe* reference genome (ASM294v2) using BWA[8] (v0.7.12) with a mapping quality cutoff of 20. The read alignments were further processed by SAMtools[9] (v1.2), Picard tools (v1.131) (https://broadinstitute.github.io/picard/), and GATK[10] (v3.5-0) for indexing, sorting, realignment, and duplicates removal. Variant calling was subsequently performed using two different tools, GATK's HaplotypeCaller[11] (v3.5-0; option: -ploidy 1) and FreeBayes[12] (v1.0.1-2; options: -p 1 --genotype-qualities) respectively, based on the read alignment file of each sample. The called variants were further processed by vt[13] (v0.5772) and vcflib[14] (v1.0.0-rc1) for variant normalization and filtering. The variant filtering option for the GATK call is QUAL > 20 & QUAL / AO > 10 & SAF > 0 & SAR > 0 & RPR > 1 & RPL > 1, while that for the FreeBayes call is QUAL > 20. For each sample, a consensus call set (*i.e.*, variants called by both variant calling tools) was further generated. The called variants were also manually inspected in IGV[15] (v2.3.70). Finally, Ensembl-VEP[16] (v83) was used to evaluate the potential functional impacts of called variants.

For the population-based pooled sequencing samples, the SNV calls were annotated with VEP (v115.2; options: --distance 500 --pick)[16] based on the *S. pombe* reference genome assembly (version ASM294v2). The non-intergenic SNV calls that have population allele frequencies ≥0.01 and are also detected in clone samples were used for the subclonal analysis. The clonal structure and clonal prevalence within each analyzed population were inferred using PyClone-VI (v0.1.6; options: -d beta-binomial -r 10 --seed 123)[17]. Clonal phylogenies originating from the same ancestor were reconstructed using PhyClone (v0.7.1; options: --outlier-prob 0.001 --num-chains 4 --seed 123)[18]. Finally, the temporal dynamics of the major clones within each population were visualized as Muller diagrams (Extended data Fig. 1g) generated using the R package fishplot (v0.5.2)[19].

**Generation time, cell size at division and DNA content analysis**

Population doubling times, referred to as population generation times, were determined in batch cultures. Cell size at division was determined from images of blankophor-stained cells (1 mg/mL) using the ImageJ PointPicker plugin. For blankophor images in Fig. 2e, Fig. 5b-c and Extended data Fig. 2a, the brightness and contrast were adjusted for display purposes. For DNA content analyses, cells were fixed in 70% cold ethanol, washed in 50 mM sodium citrate, treated with RNAse A (0.1 mg/mL) and stained with propidium iodide (2 mg/mL). DNA content was determined using a BD Accuri C6 flow cytometer. Note that the fission yeast cell cycle has a short G1, and S phase occurs prior to cytokinesis. Therefore, during most of the cell cycle, cells have a 2C DNA content. The appearance of a 1C peak reflects an extension of G1 duration, with cytokinesis occurring prior to DNA replication.

**Induction of synchronous meiosis in diploid *pat1-114* cells**

Diploid cells carrying the *pat1-114* temperature-sensitive allele were grown at 25 ºC and switched to minimal medium without nitrogen (EMM-N) for 16 h. The cultures were then shifted to the restrictive temperature of 34 °C and $NH_4Cl$ was added as a nitrogen source (T=0). After 200 min, samples were collected and heat fixed, and the number of nuclei was assessed by DAPI staining as above.

**Microfluidic chips, live-cell imaging and single-cell analyses**

The microfluidic device designs were generated in AutoCAD (Autodesk) and transferred onto chrome photomasks (JD Photo Data). The complete design file can be downloaded at https://github.com/TAspert/FissionYeast_Traps. Fabrication of the master molds was carried out using two successive rounds of conventional photolithography. The first layer, consisting of an array of 2000 traps, was formed by depositing a 5.25 µm layer of SU-8 2005 negative photoresist (MicroChem, USA). This was achieved by spin-coating 3 mL of the resist at 2500 rpm for 30 s on a 3” wafer (Neyco, France) using a WS650 spin coater (Laurell, USA). The coated wafer underwent a 3 min soft bake at 95 °C, followed by UV exposure at 365 nm with an energy dose of 120 mJ/cm² using a UV-KUB3 mask aligner (Kloé, France), after ensuring a perfect contact between the mask and the resist. A post-exposure bake, identical to the soft bake, was then performed prior to development in SU-8 developer (MicroChem, USA). For the second layer containing the channel structures, a 30 µm layer of SU-8 2025 (MicroChem, USA) was spin-coated at 2500 rpm for 30 s. The soft bake consisted of 3 min at 65 °C followed by 6 min at 95 °C. The wafer was aligned with the second layer mask and exposed to 120 mJ/cm² UV light. Post-exposure baking conditions also matched those of the soft bake. After processing each layer, a hard bake at 150 °C for 15 min was carried out to seal microcracks and enhance the stability of the photoresist. Finally, the completed master molds were passivated by treating them with chlorotrimethylsilane by chemical vapor deposition.

For live-cell imaging using these microfluidic devices (Fig. 1g, h; Fig. 2b; Extended data Fig. 2e, f), PDMS microfluidic chips were prepared following standard procedures[20]. Briefly, a 10:1 PDMS:hardener mix (Sylgard 184, Dow Corning, USA) was degassed for ~45 min at room temperature in a vacuum chamber, poured on the mold and allowed to harden for 2 h at 70 °C. The chips were then detached, inlets were made using a biopsy punch, and the devices were bond to microscopy-grade coverslips using plasma treatment (3 min; Harrick Plasma, USA). The chips were left at room temperature for 15 min prior to use. Cells were then injected in the chips and imaging experiments were performed at 32 ºC with a constant flow of medium (5 µL/min) to maintain optimal growth conditions using flow and pressure controllers (Elvesys, France). The Nhp6-mCherry and Plo1-GFP markers were used to monitor cell cycle events. The determination of the single-cell criteria for Fig. 1g and Extended data Fig. 2e, f are provided in their respective legends. All imaging experiments were performed using either an inverted Zeiss Axio Observer (Carl Zeiss Microscopy LLC, Germany) equipped with a Lumencor Spectra X illumination and an Orca Flash 4.0V2 sCMOS camera (Hamamatsu Photonics, Japan) or a similar system equipped with a Yokogawa CSU-W1 spinning disc module coupled with both an Orca Flash 4.0V2 and a Quest camera (Hamamatsu Photonics, Japan). Acquisition was performed using the Visiview software (Visitron Systems GmbH, Germany).

**Western blot analyses**

For Western blot analyses of whole-cell extracts (Extended data Figs. 2g, 3e, 4b-c, 6b), 10 mL of culture at 0.4-0.5 OD were collected and resuspended in 1.5 mL IPP50 (10 mM Tris-HCl pH 8, 150 mM NaCl, 0.1% NP-40) + protease inhibitors (PMSF; Complete Protease Inhibitor Cocktail, Sigma Aldrich). Total protein extracts were prepared using glass beads and a Precellys 24 homogenizer (2 cycles of 20 s at 8000 rpm). Sample concentrations were determined using the BCA protein assay kit (Thermofisher) and appropriate amounts of proteins were then diluted using 5X sample buffer (300 mM Tris HCl pH 6.8, 20% beta-mercaptoethanol, 20% SDS, 0.05% bromophenol blue, 25% glycerol). After boiling for 5 min at 99 ºC, samples were loaded in equal amounts (except otherwise noted) on 10% SDS-polyacrylamide gels. Antibodies were used at the following dilutions: anti-HA-HRP (1:10000; Abcam ab1190), Anti-Tat1 (tubulin; 1:20000; a kind gift from K. Gull), anti-GFP (1:1000; Roche 11814460001). Biotinylated proteins (Extended data Fig. 6b) were detected using HRP-streptavidin (1:1000, RPN1231, Sigma Aldrich).

***In vivo* analysis of Cdk activity**

*synCut3-mCherry biosensor*

The synCut3-mCherry biosensor was previously described[4]. For the analyses in Fig. 2b, time-lapse experiments in single-cell trap microfluidic chambers were performed as described above. *z*-stacks (0.30 µm steps) were acquired at different time points to quantify the synCut3-mCherry signal: for each cell, sum projections were generated in ImageJ throughout the time lapse and used to identify the time point at which the nuclear intensity was maximal (as determined by intensity quantification along a line scan throughout the nucleus). This time point, which may differ between cells, was then used to evaluate the peak nuclear/cytoplasmic (N/C) ratio of synCut3-mCherry signal. To this end, the contours of both the cell and the nucleus were manually drawn. The total intensities for the nucleus *vs*. the cytoplasm (total cell intensity – nucleus intensity) were then determined. The nuclear and cytoplasmic areas (in pixels) were also extracted. Next, the average background signal per pixel was determined at the same time point using an area outside of the cell. This allowed for background subtraction from both the nuclear and cytoplasmic synCut3-mCherry signals using the respective areas of each compartment. The N/C ratio for synCut3-mCherry was then calculated using these corrected values. For the analyses in Fig. 2d (asynchronous cultures), *z*-stacks (0.30 µm steps) were acquired from cells mounted between slide and coverslip. Cells that showed detectable nuclear signal and had not undergone nuclear division (single round nucleus) were analyzed as above. Note that in this case, the analyses were not restricted to the time of peak N/C ratio.

*Eevee-spCDK FRET biosensor*

The Eevee-spCDK FRET biosensor was previously described[5]. Live-cell imaging for all FRET experiments was performed using an epifluorescence inverted microscope (IX83; Olympus) equipped with a Prime sCMOS camera (Photometrix), a Lumencor Spectra X illumination and an oil-immersion lens (UPLXAPO 60X). Illumination and fluorescence filters were as follows: excitation filter: 438/24; excitation dichroic mirror: FF458 Di-02; emission filters (Semrock): 483/32-25 (CFP) and 520/28 (FRET). Vegetatively growing fission yeast cells were concentrated by centrifugation (3000 rpm, 1 min) and resuspended in 100 µL of YE supplemented with adenine (225 mg/L). Cells were then loaded in an ONIX microfluidic platform (Merck; trapping chamber Y04C-04) by imposing a pressure of 8 psi for 15 s. During imaging, growth conditions were maintained through constant media perfusion (1 psi) at a temperature of 30 °C. All images were analyzed using Fiji/ImageJ. The background was subtracted by the rolling-ball method (radius, 50.0 pixels). Single cell tracking was performed using an optimized tracking plugin for Fiji, LIM Tracker[21]. CFP images were used to define regions of interest (ROIs) corresponding to the nucleus. Fluorescence intensities in the CFP and FRET channels were quantified for each cell as the mean within the ROI. The FRET/CFP ratio was then determined and used as an index of Cdk activity. Cell cycle times were determined as the timing between two nuclear divisions (Extended data Fig. 5b). To compare data, the FRET/CFP traces for each cell were aligned either to the preceding nuclear division (Fig. 3a, Extended data Fig. 5a) or to the Cdk activity rise time (Fig. 3c). The Cdk activity rise time was obtained using specific thresholds for the smoothened FRET/CFP values (see legend of Fig. 3; smoothing was performed by using moving averages of window size 3).

**TurboID assays and mass-spectrometry analyses**

*Sample preparation and pull-down of biotinylated proteins*

50 mL of culture at OD 0.4-0.5 were spun down 2500 rpm for 5 min at 4 ºC. Pellets were then lysed in 500 µL of cold RIPA buffer (50 mM Tris-HCl pH 7.5, 150 mM NaCl, 1.5 mM MgCl2, 1 mM EGTA, 0.1% SDS, 1% NP-40, 0.4% sodium deoxycholate, 1 mM DTT, 1 mM PMSF, 1× PLAAC, 1× cOmplete) using glass beads and a Precellys 24 homogenizer (4 cycles of 20 s at 8000rpm). The sample volume was then increased to 1000 µL with cold RIPA buffer, and the lysates were sonicated three times for 10 s at 20% intensity (Branson SSE-1). DNA and RNA were then digested using 500 units of benzonase for 1 h at 4°C. Protein concentrations were determined and normalized using the BCA protein assay kit (Thermofisher, USA). 3 mg of total proteins were incubated with 30 µL of streptavidin-sepharose beads (Thermofisher, USA) in 1 mL of RIPA buffer (with 0.4% SDS) for 3 h at 4 °C. Samples were subsequently washed twice with wash buffer (50 mM Tris-HCl pH 7.5, 2% SDS), three times with RIPA buffer containing 1 mM DTT, and five times with 20 mM ammonium bicarbonate. For Western blot analysis (Extended data Fig. 6b, right panel), beads were incubated in sample buffer as above at room temperature for 15 min. Samples were then boiled for 15 min at 95 ºC prior to loading on gel. Membrane stripping was performed by three washes of 5 min in stripping buffer (15 g glycine, 1 g SDS, 10 mL Tween 20 for 1 L, adjusted to pH 2.2 with HCl).

*Materials for mass spectrometry*

MS grade Acetonitrile (ACN), MS grade $H_2O$, ammonium bicarbonate ($NH_4HCO_3$), MS grade formic acid (FA), were from ThermoFisher Scientific. Sequencing-grade Trypsin/Lys C mix was from Promega. Ammonium bicarbonate ($NH_4HCO_3$) was from Sigma-Aldrich. Evotips Pure were from Evosep. Aurora Elite CSI analytical column (ref. AUR3-15075C18-CSI) was from Ionopticks.

*Samples preparation prior to LC-MS/MS analysis*

40 µL of 50 mM $NH_4HCO_3$ buffer containing 1 µg of sequencing-grade Trypsin/Lys-C mix were added to each sample, which were then incubated overnight at 37 °C and 600 rpm. The next day, samples were placed on a magnetic rack, the supernatant was transferred to another fresh tube and evaporated dry, before adding 80 µL of $H_2O$ containing 0.1% of FA. 5 µL of this mixture was loaded and desalted on Evotips for each sample, according to manufacturer's procedure, prior to LC-MS/MS analysis.

*LC-MS/MS acquisition*

Samples were analyzed on a timsTOF Pro 2 mass spectrometer (Bruker Daltonics, Germany) coupled to an Evosep one system (Evosep, Denmark) operating with the Whisper Zoom 40SPD method developed by the manufacturer. Briefly, the method is based on a 31-min gradient and a total cycle time of 38 min with a C18 Aurora Elite CSI analytical column (15cm x 75 µm, 1.7µm beads) from Ionopticks, equilibrated at 50 °C and operated at a flow rate of 200 nL/min. $H_2O$/0.1 % FA was used as solvent A and ACN/0.1 % FA as solvent B. The timsTOF Pro 2 was operated with a DIA-PASEF method comprising 12 pydiAID frames with 3 mass windows per frame resulting in a cycle time of 0.975 s as described in Bruker application note LCMS 218. Collisional energy was ramped stepwise as a function of ion mobility.

*Data analyses*

MS raw files were processed using Spectronaut version 19.4.241104.62635. Data were searched against the *Schizosaccharomyces pombe* UniProt database (downloaded 2024_06, 5117 entries). Parent mass tolerance was set to 20 ppm, with fragment mass tolerance at 0.05 Da. Specific tryptic cleavage was selected and a maximum of 2 missed cleavages was authorized. For identification, the following post-translational modifications were included: Acetyl (Protein N-term), Oxidation (M), Phosphorylation (STY) and Deamidation (NQ) as variables and Cys-Cys (C) as fixed. Identifications were filtered based on a 1% Q-value threshold at both precursor and protein levels. Quantification was performed using Spectronaut Quantification Module, with all default parameters. Proteins were inferred using the automatic features of Spectronaut, with the algorithm IDPicker. Multivariate statistics on protein or peptide measurements were performed using Qlucore Omics Explorer 3.9 (Qlucore AB, Sweden). A positive threshold value of 1 was specified to enable a log2 transformation of abundance data for normalization *i.e.* all abundance data values below the threshold will be replaced by 1 before transformation. The transformed data were finally used for statistical analysis *i.e.* evaluation of differentially present proteins or peptides between two groups using a Student's bilateral t-test.

**Phosphoproteomic analyses following Pab1 depletion**

*Sample preparation*

Early exponential cultures of control cells (containing the auxin-inducible degron background, *Padh15-skp1-At-Tir1-2NLS-Padh15-sk1-Os-Tir1*) and *nmt41-3PK-miniAID-pab1* cells (referred to as *AID-pab1;* see Supplementary Table 6 for the complete genotypes) were treated with 15 µM thiamine and 0.5 mM naphthaleneacetic acid (NAA). 40 mL samples were then collected at the indicated times by filtration and fixed in 20 % trichloroacetic acid (TCA). Once all samples were collected, cells were washed with acetone and resuspended in lysis buffer (50 mM ammonium bicarbonate, 5 mM EDTA pH 7.5, 8 M urea). Subsequently, cells were broken by glass bead beating and extracts were cleared by centrifugation. Protein concentration was determined and 200 µg protein equivalent was further processed for TMT-plex mass tag labeling and mass spectrometry as described[22].

*Western blot analysis*

For the Western blot in Extended data Fig. 6c, samples were prepared by diluting 30 µg of each cell extract above with 2X concentrated SDS-PAGE loading buffer. Protein extracts were resolved by SDS-polyacrylamide gel electrophoresis, followed by transfer to nitrocellulose membranes. AID-Pab1 was detected by means of its N-terminal 3PK tag (mouse anti-V5l, BioRad clone SV5-Pk1, 1:1000) and Cdc2 served as a loading control (anti-PSTAIR, Abcam ab10345, 1:1000). HRP-conjugated secondary antibodies were used at 1:10000 dilution.

**Competition experiments**

Competition experiments were performed in the eVOLVER continuous culture system[23]. *gsf2Δ::HYG* and *gsf2Δ::KAN spo12Δ::KAN* cells were separately grown in batch cultures. At day 0 of the competition experiment, eVOLVER vials were inoculated with a 1:1 ratio of each genotype (total volume = 25 mL). At the indicated time points (Fig. 5d), each culture was sampled and ~ 200 cells were plated on non-selective YE4S plates. Plates were incubated at 32 °C until colonies appeared. Plates were then replica-plated on YE4S + hygromycin (200 µg/mL) and YE4S + kanamycin (100µg/mL) to determine the percentage of each genotype in the cultures.

**Model generation using Alphafold3**

The predicted structures of various phosphatase complexes without and with Spo12 and other protein inhibitors were created by using the AlphaFold3[24] server interface (alphafoldserver.com) with the default parameters. Protein and ion combinations used in this work are summarized in Supplementary Table 5. UniProt protein sequence accession numbers are: spSpo12 (Q10189), spPaa1 (Q9UT08), spPab1 (Q12702), spPpa2 (P23636), spIgo1 (P79058), scSpo12 (P17123), scPaa1 (P31383), scPab1 (Q00362), scPpa2 (P23595), hsPPP2R1A (P30153), hsPPP2R2A (P63151), hsPPP2CA (P67775), and hsARPP19 (P56211). Structure-based images were prepared by using PyMOL (Schrödinger).

**Mathematical modelling**

All the details, ordinary differential equations, simulation parameters and additional simulations for the model in Fig. 6 and Extended data Figs. 11-13 are provided in the Supplementary Model file.

**Statistical analyses**

For Fig. 2b, c and Fig. 3c: two-tailed Mann-Whitney U test. For Extended data Fig. 11e: two-tailed independent t-test for normal distribution. For the Turbo-ID experiments (Supplementary Table 2), two-tailed Student's t-test. For the phosphoproteomic study upon Pab1 depletion (Supplementary Table 3): linear regression t-test with Benjamini–Hochberg FDR correction.

**Data availability**

The whole genome sequencing data are available on the NCBI Sequence Read Archive database under the BioProject accession number PRJNA1344658 (https://www.ncbi.nlm.nih.gov/bioproject/PRJNA1344658).

The mass spectrometry data for the TurboID and Pab1 depletion experiments are available upon request.
