## Supplementary Data for "Regulation of cell proliferation by a novel feedback system on Cdk function"

### Extended data Fig. 1

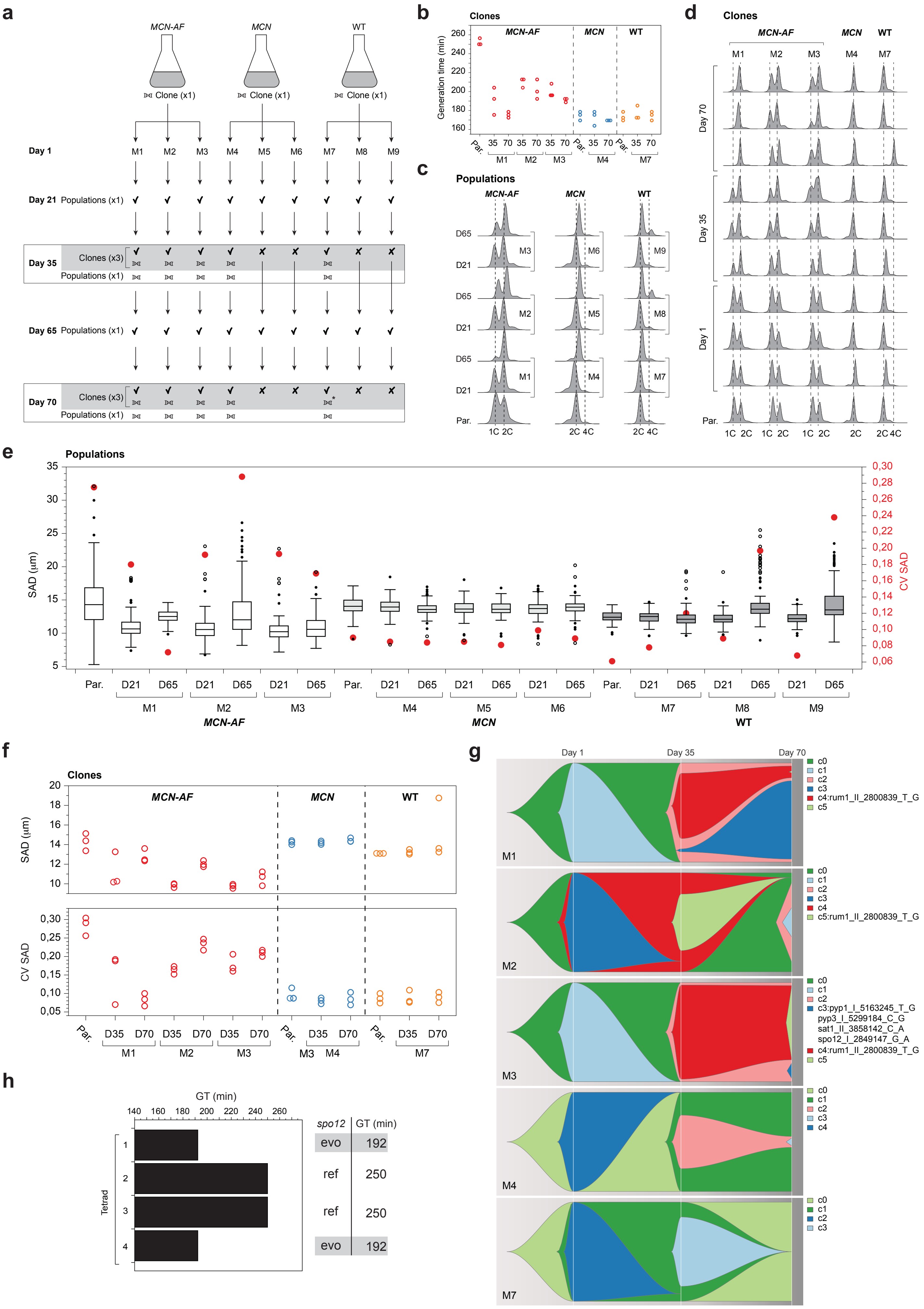

### Extended data Fig. 2

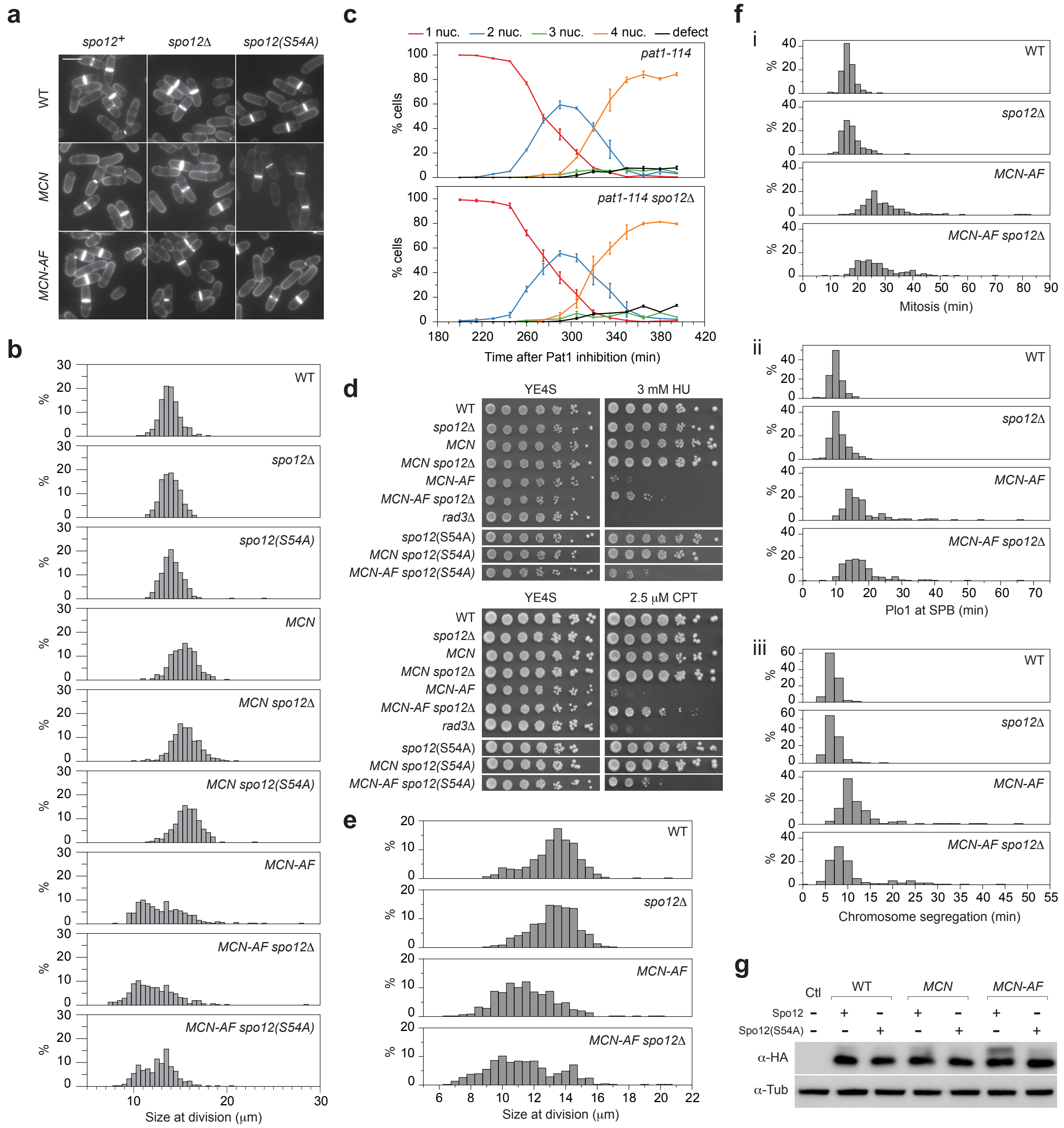

### Extended data Fig. 3

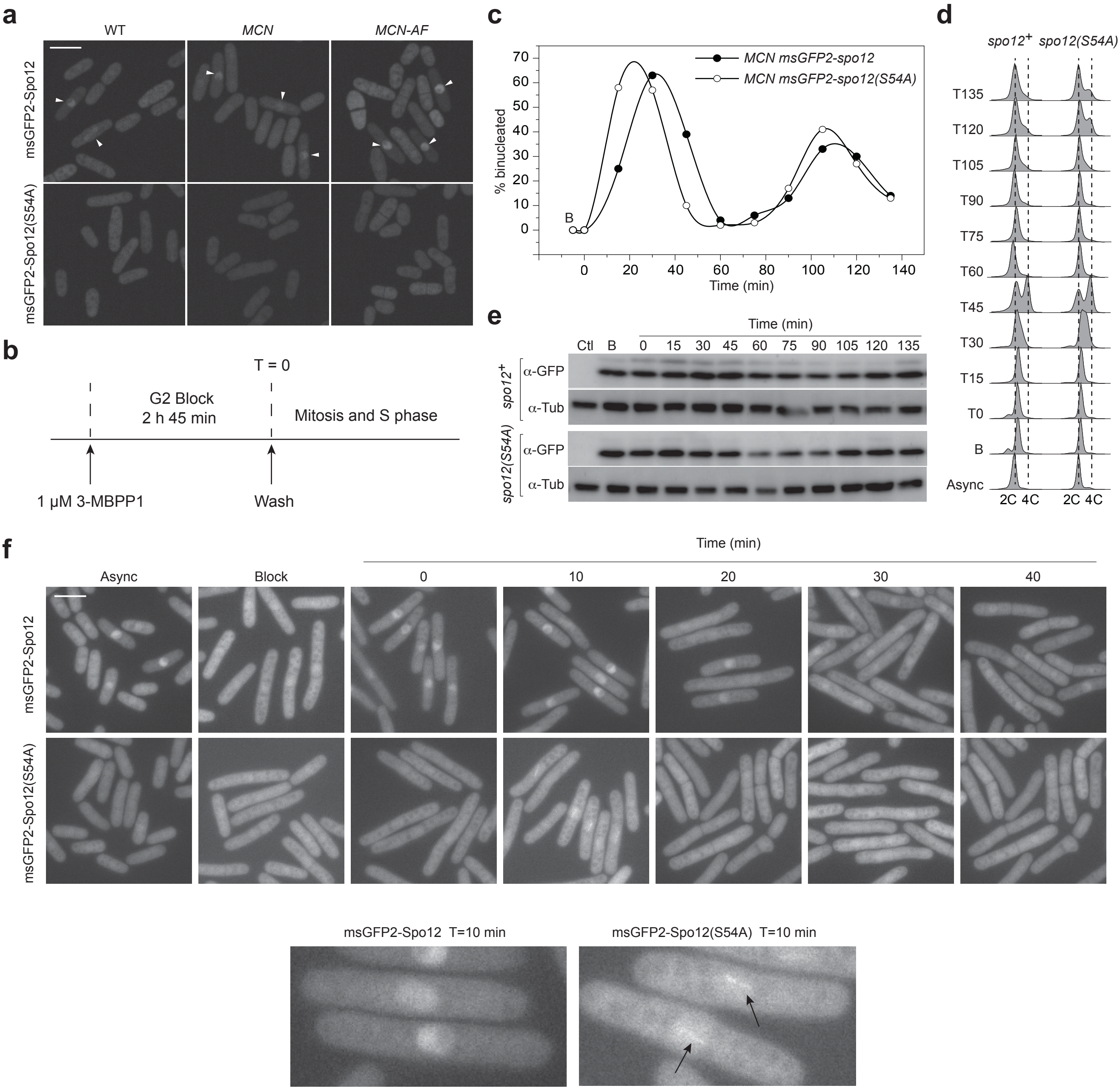

### Extended data Fig. 4

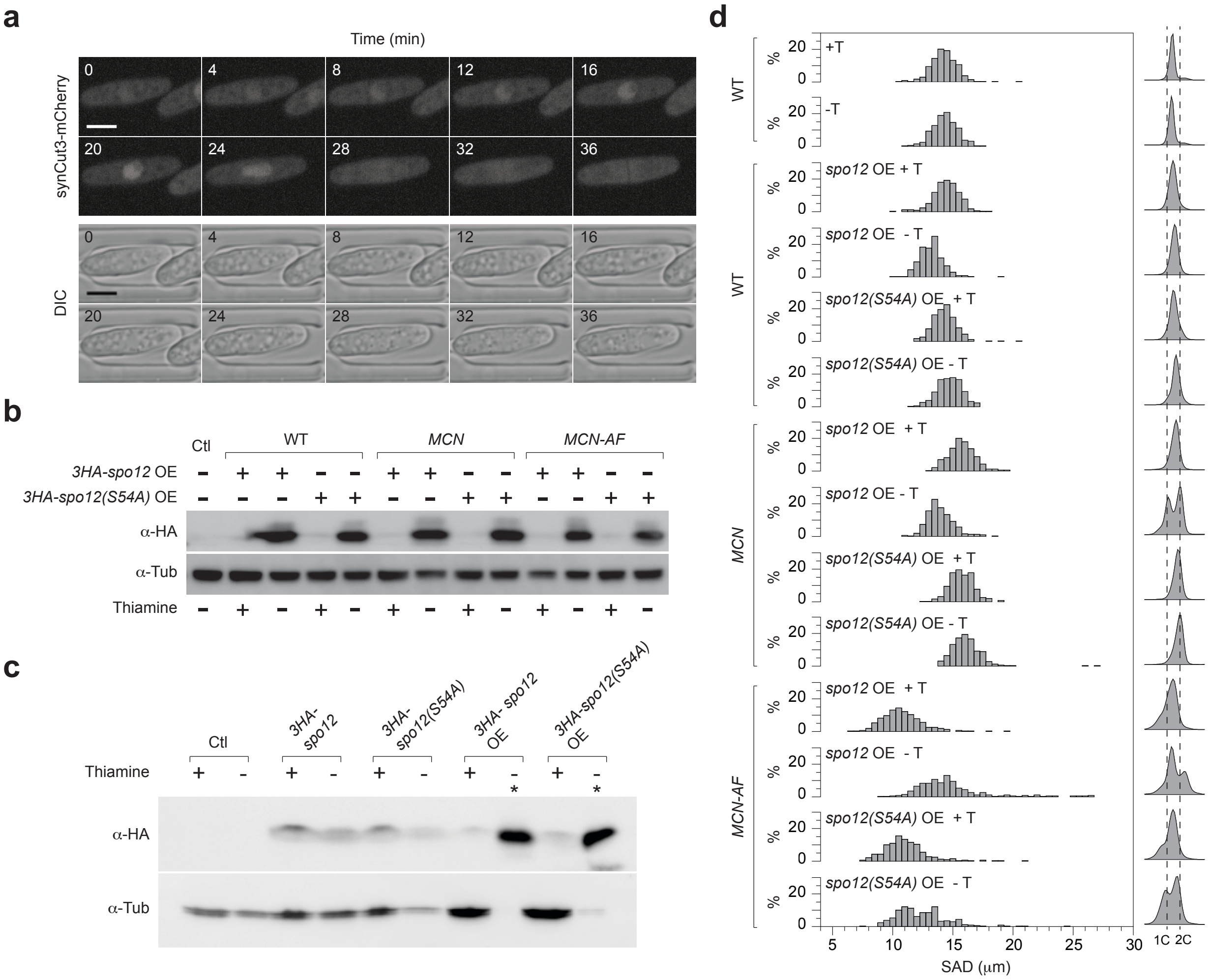

### Extended data Fig. 5

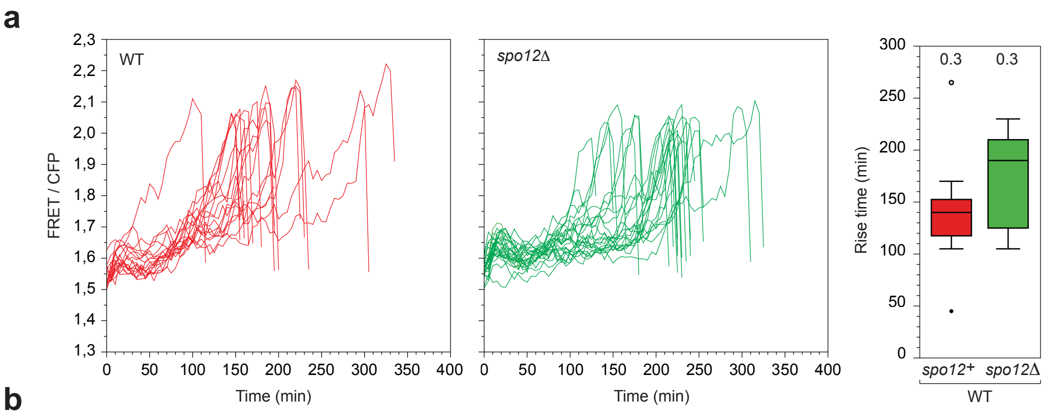


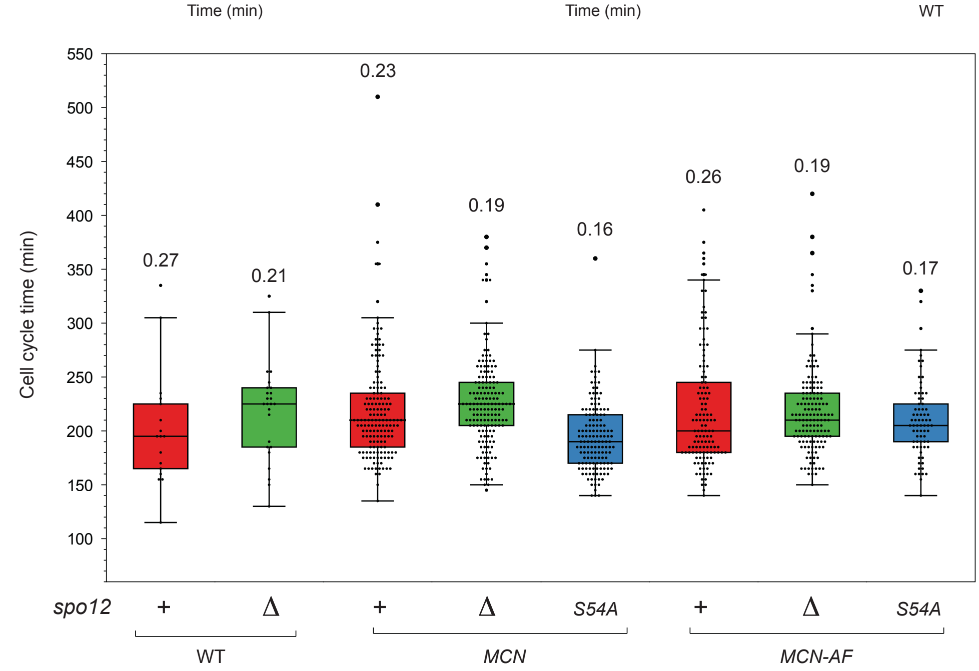

### Extended data Fig. 6

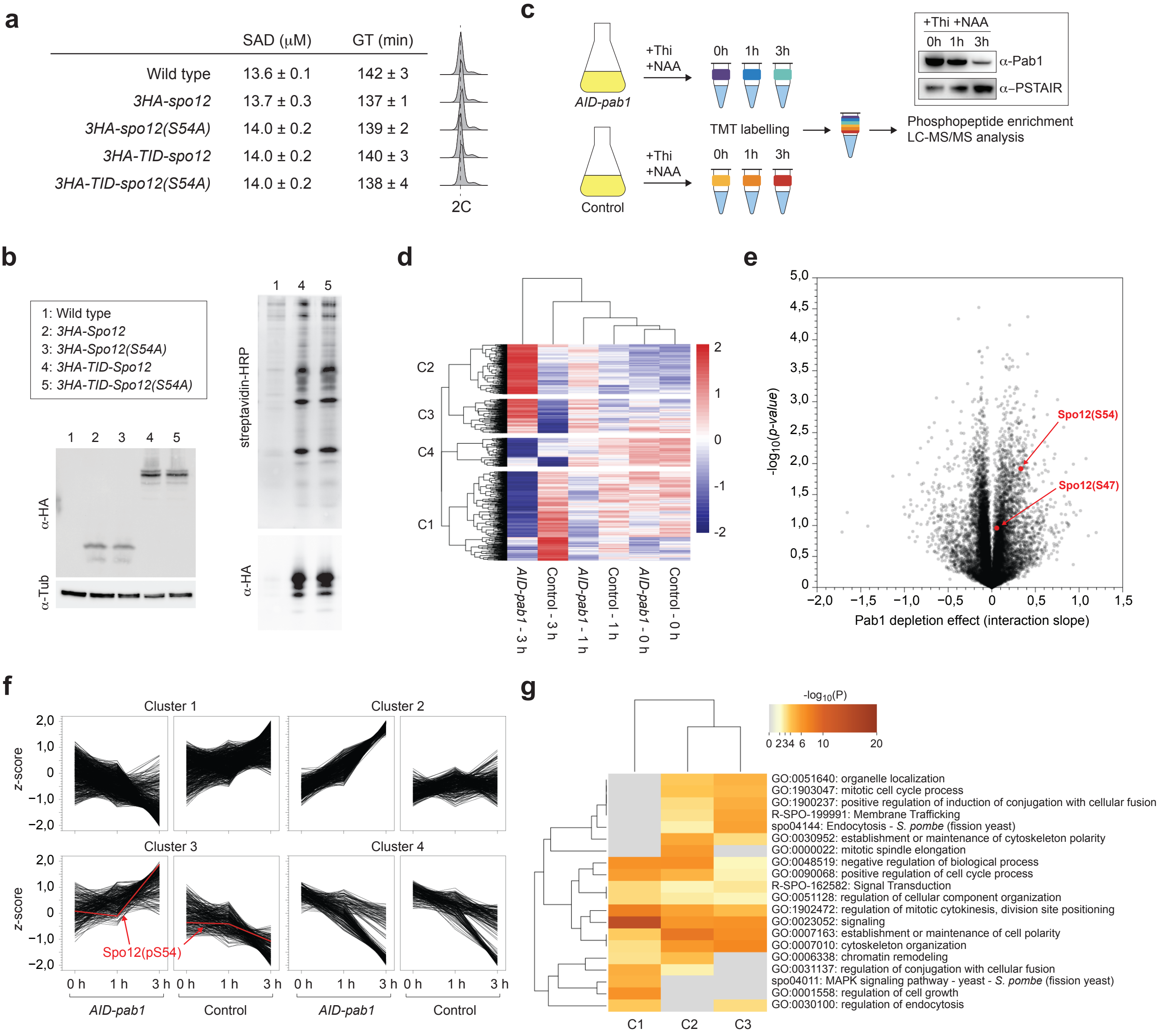

#### Extended data Fig. 7

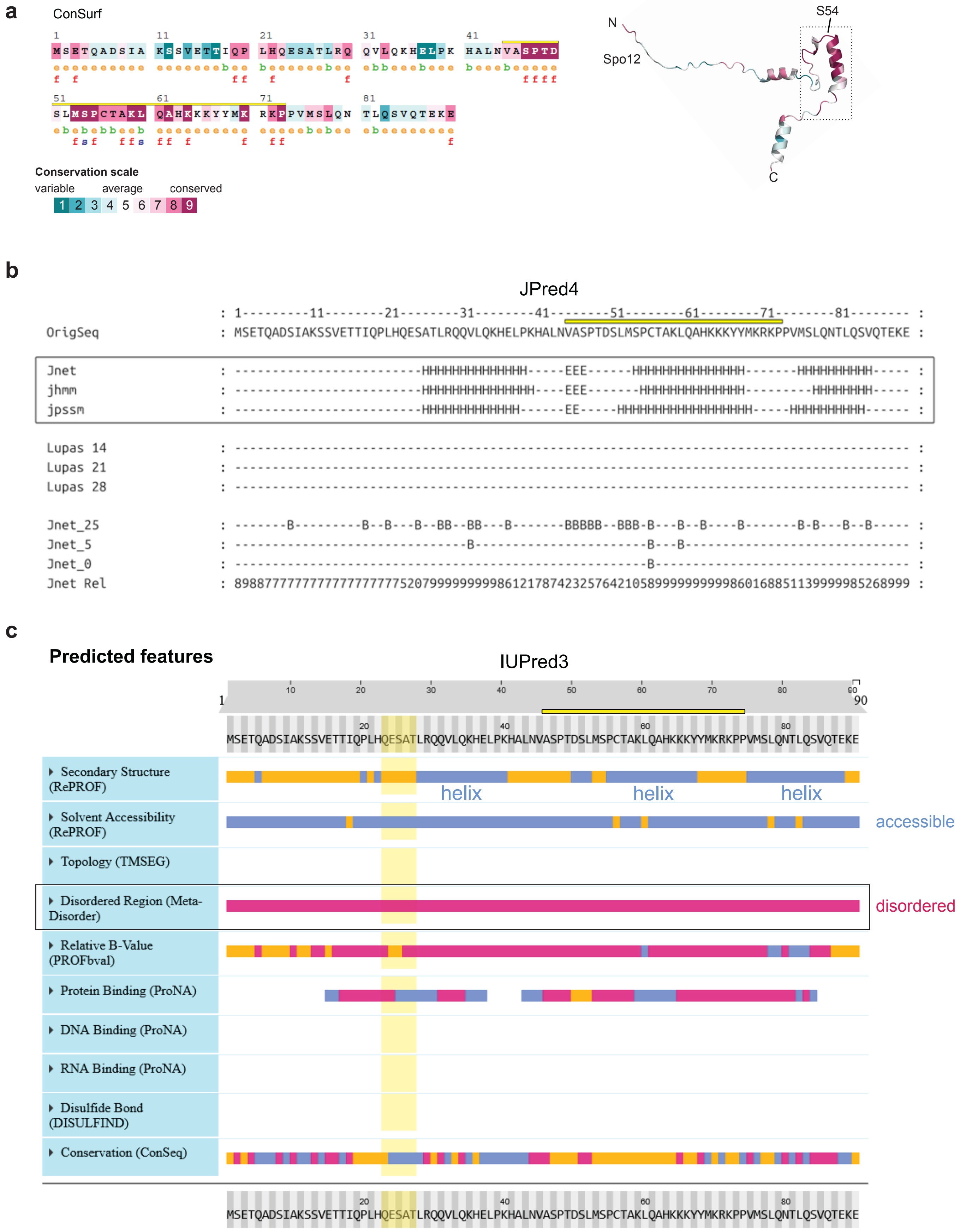

### Extended data Fig. 8

**a**

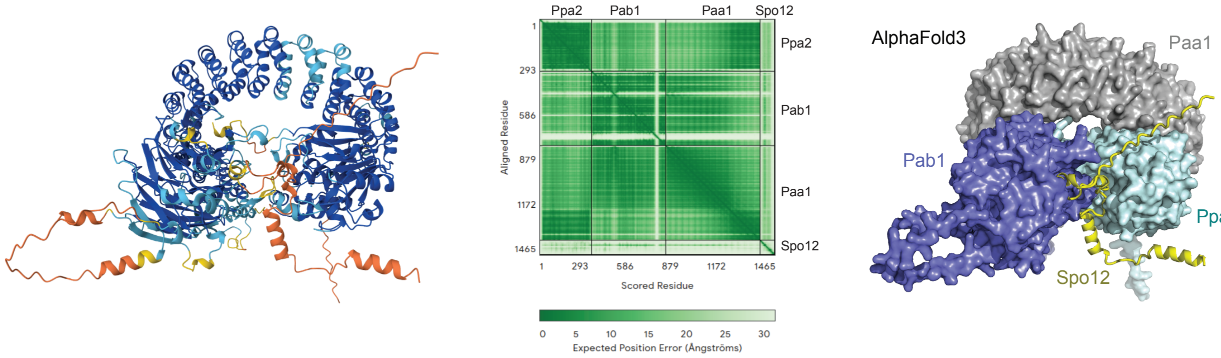


**b**

| Complex qTM | Complex tTM | Chain paring | Description | Scientific Name | Prob. | Seq. Id. | E-Value | Position in query | Alignment |
|---|---|---|---|---|---|---|---|---|---|
| 0.83 | 0.88 | A → 8ttb-assembly1_C | Cryo-EM structure of the PP2A:B55-ARPP19 c... | Homo sapiens | 1.00 | 79.4 | 7.01e-58 | 20 – 321 | |
| | | B → 8ttb-assembly1_B | Cryo-EM structure of the PP2A:B55-ARPP19 c... | Homo sapiens | 1.00 | 53.5 | 2.05e-56 | 10 – 463 | |
| | | C → 8ttb-assembly1_A | Cryo-EM structure of the PP2A:B55-ARPP19 c... | Homo sapiens | 1.00 | 53.1 | 8.02e-39 | 10 – 586 | |
| | | D → 8ttb-assembly1_D | Cryo-EM structure of the PP2A:B55-ARPP19 c... | Homo sapiens | 0.00 | 42.8 | 2.62e+2 | 71 – 77 | |
| 0.82 | 0.89 | A → 8so0-assembly1_C | Cryo-EM structure of the PP2A:B55-FAM122A ... | Homo sapiens | 1.00 | 79.4 | 4.33e-58 | 20 – 321 | |
| | | B → 8so0-assembly1_B | Cryo-EM structure of the PP2A:B55-FAM122A ... | Homo sapiens | 1.00 | 53.5 | 3.40e-55 | 10 – 461 | |
| | | C → 8so0-assembly1_A | Cryo-EM structure of the PP2A:B55-FAM122A ... | Homo sapiens | 1.00 | 53.1 | 4.27e-39 | 10 – 586 | |

**c**

| Complex qTM | Complex tTM | Chain paring | Description | Scientific Name | Prob. | Seq. Id. | E-Value | Position in query | Alignment |
|---|---|---|---|---|---|---|---|---|---|
| 0.86 | 0.93 | A → 8so0-assembly1_C | Cryo-EM structure of the PP2A:B55-FAM122A ... | Homo sapiens | 1.00 | 78.1 | 3.86e-58 | 2 – 308 | |
| | | B → 8so0-assembly1_B | Cryo-EM structure of the PP2A:B55-FAM122A ... | Homo sapiens | 1.00 | 55.6 | 1.03e-56 | 10 – 461 | |
| | | C → 8so0-assembly1_A | Cryo-EM structure of the PP2A:B55-FAM122A ... | Homo sapiens | 1.00 | 53.1 | 7.18e-40 | 10 – 586 | |
| | | D → 8so0-assembly1_D | Cryo-EM structure of the PP2A:B55-FAM122A ... | Homo sapiens | 0.00 | 3.1 | 3.21e+2 | 58 – 89 | |
| 0.85 | 0.90 | A → 8ttb-assembly1_C | Cryo-EM structure of the PP2A:B55-ARPP19 c... | Homo sapiens | 1.00 | 78.1 | 2.45e-57 | 2 – 308 | |
| | | B → 8ttb-assembly1_B | Cryo-EM structure of the PP2A:B55-ARPP19 c... | Homo sapiens | 1.00 | 54 | 5.36e-57 | 10 – 463 | |
| | | C → 8ttb-assembly1_A | Cryo-EM structure of the PP2A:B55-ARPP19 c... | Homo sapiens | 1.00 | 53.1 | 2.05e-39 | 10 – 586 | |
| | | D → 8ttb-assembly1_D | Cryo-EM structure of the PP2A:B55-ARPP19 c... | Homo sapiens | 0.00 | 20 | 2.81e+2 | 57 – 76 | |

**d**

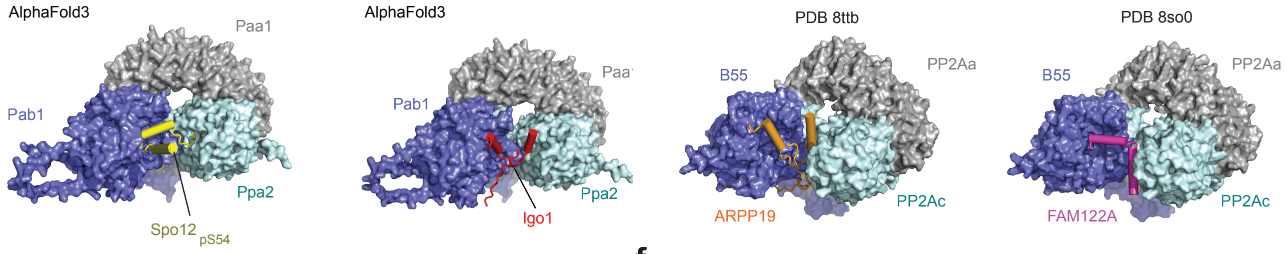


**e**

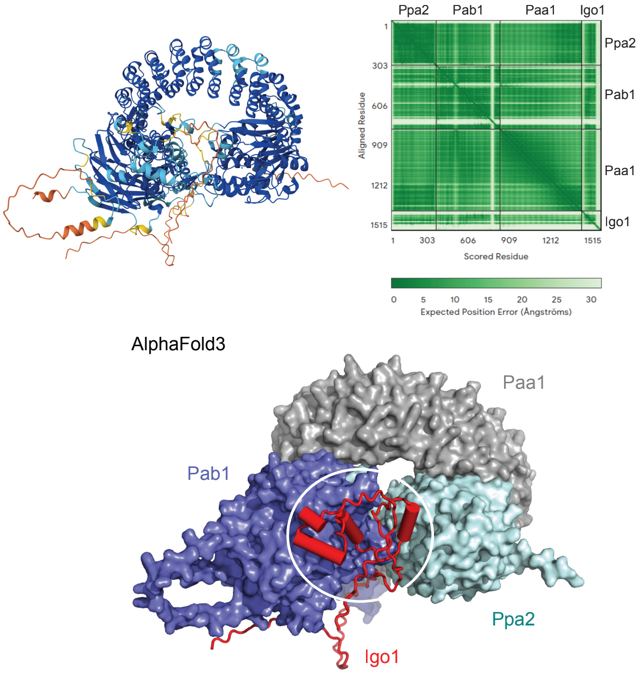


**f**

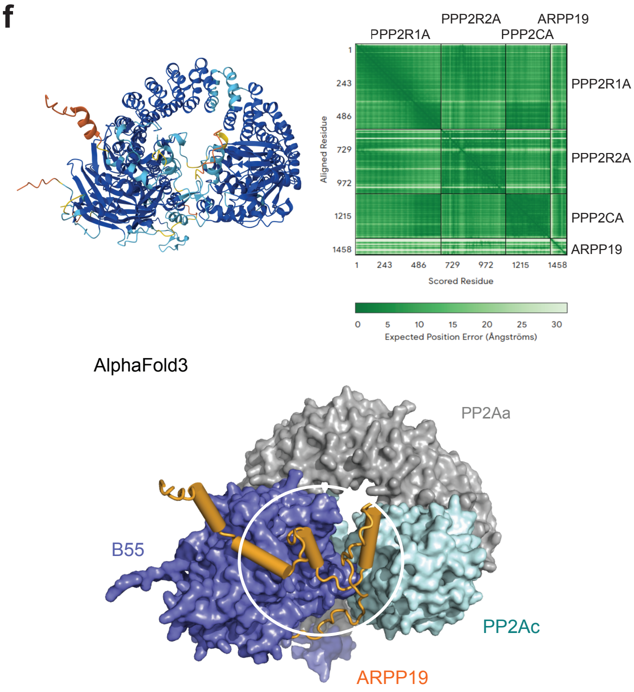

### Extended data Fig. 9

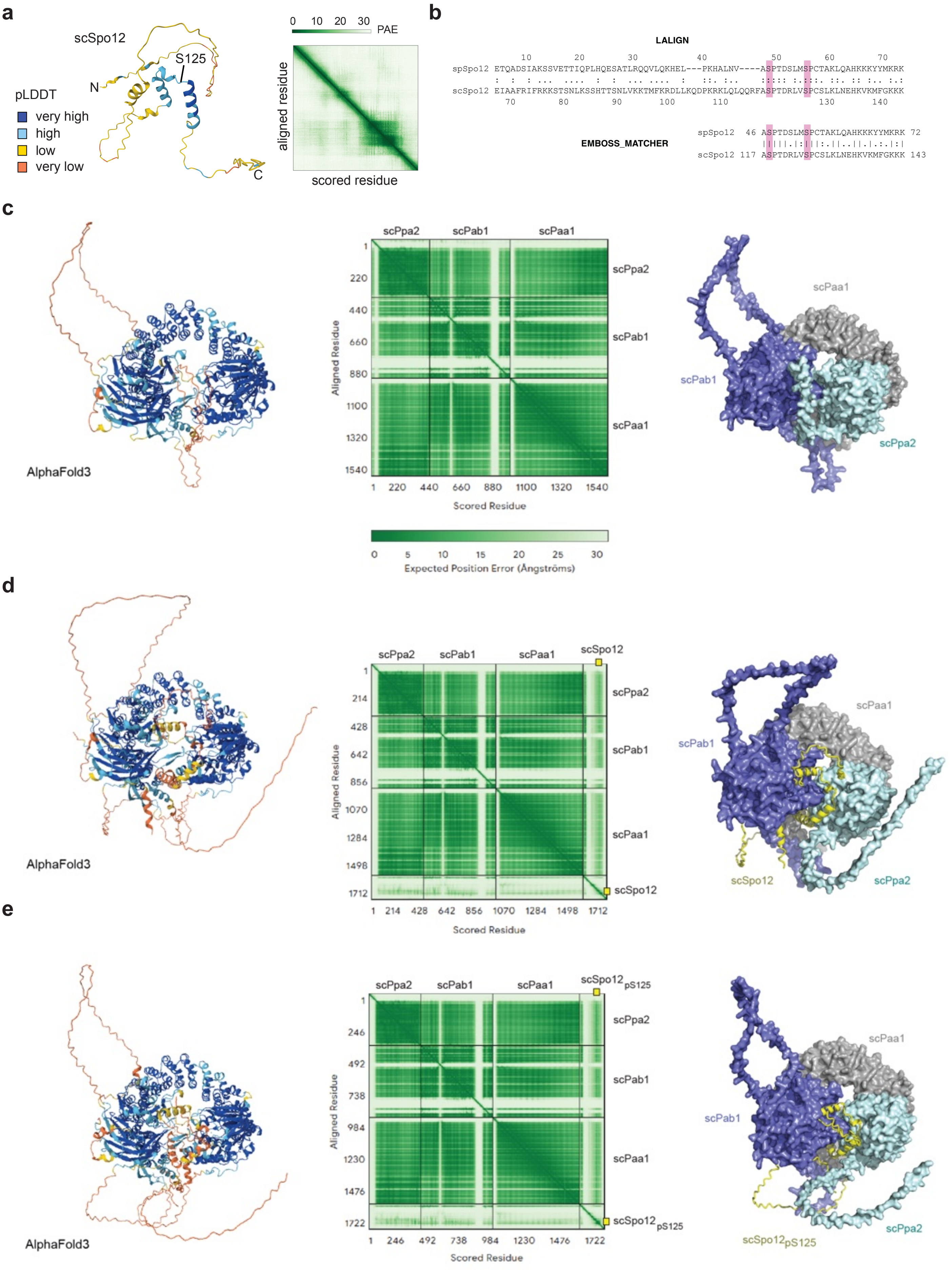

#### Extended data Fig. 10

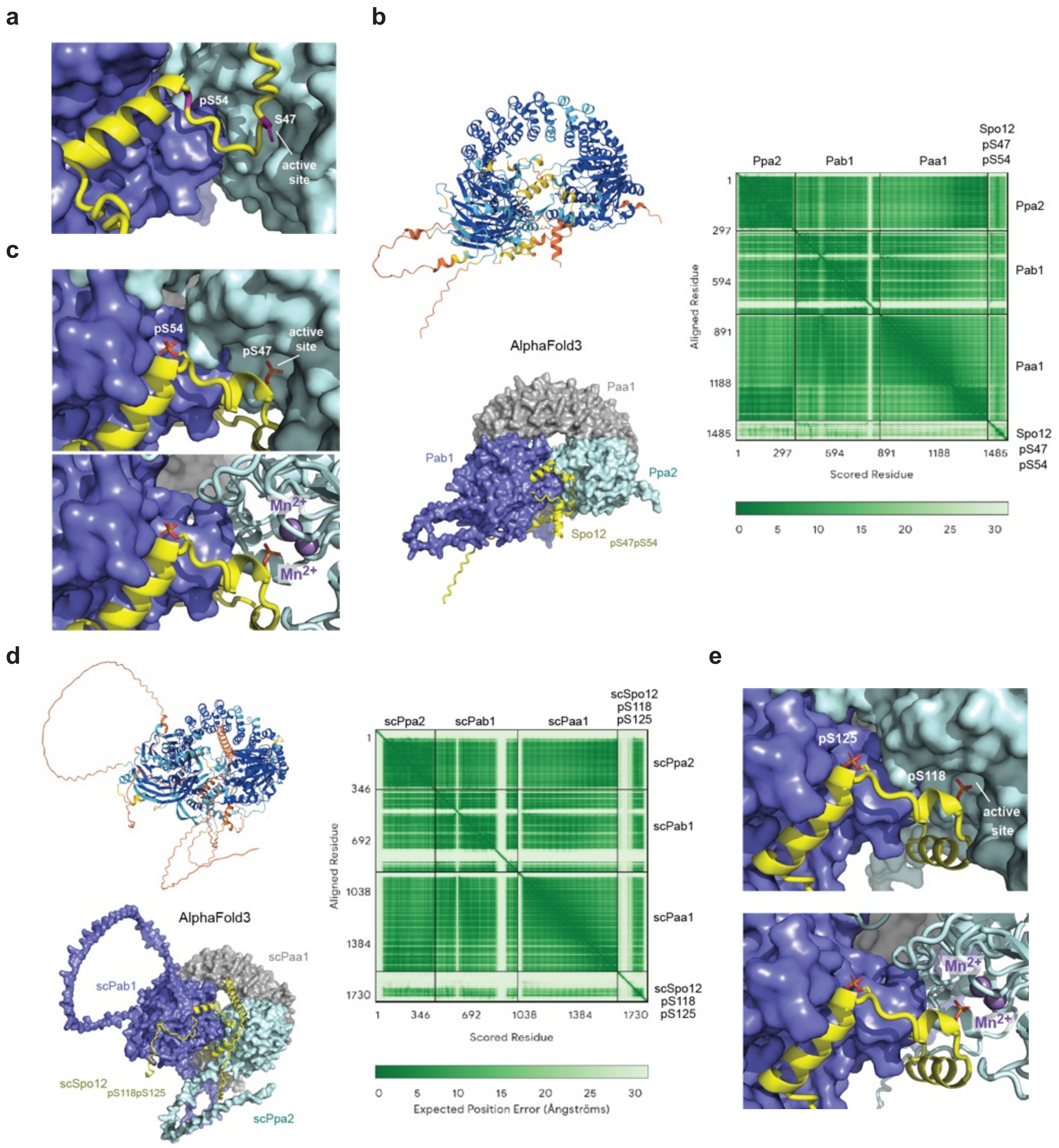

### Extended data Fig. 11

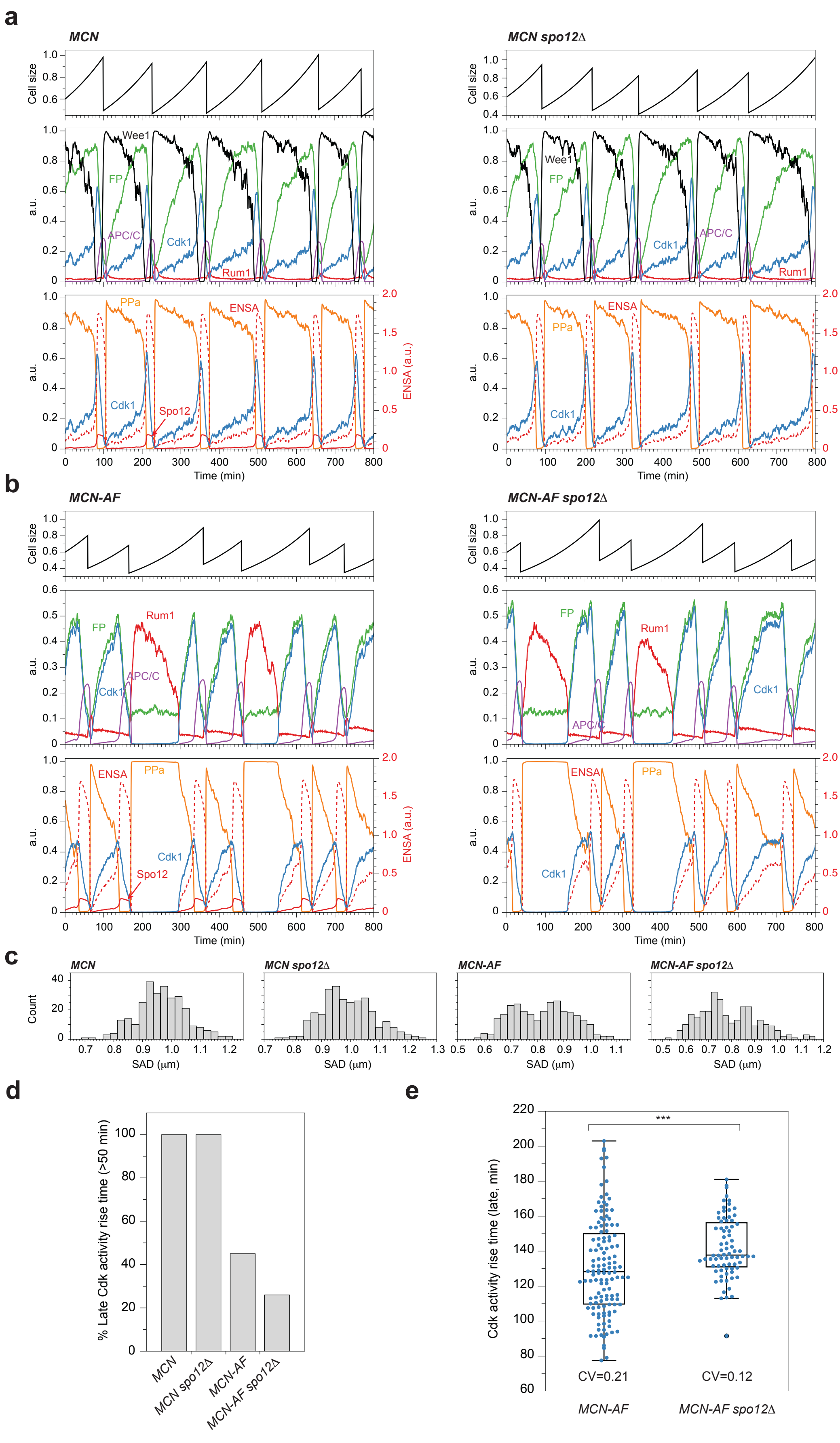

### Extended data Fig. 12

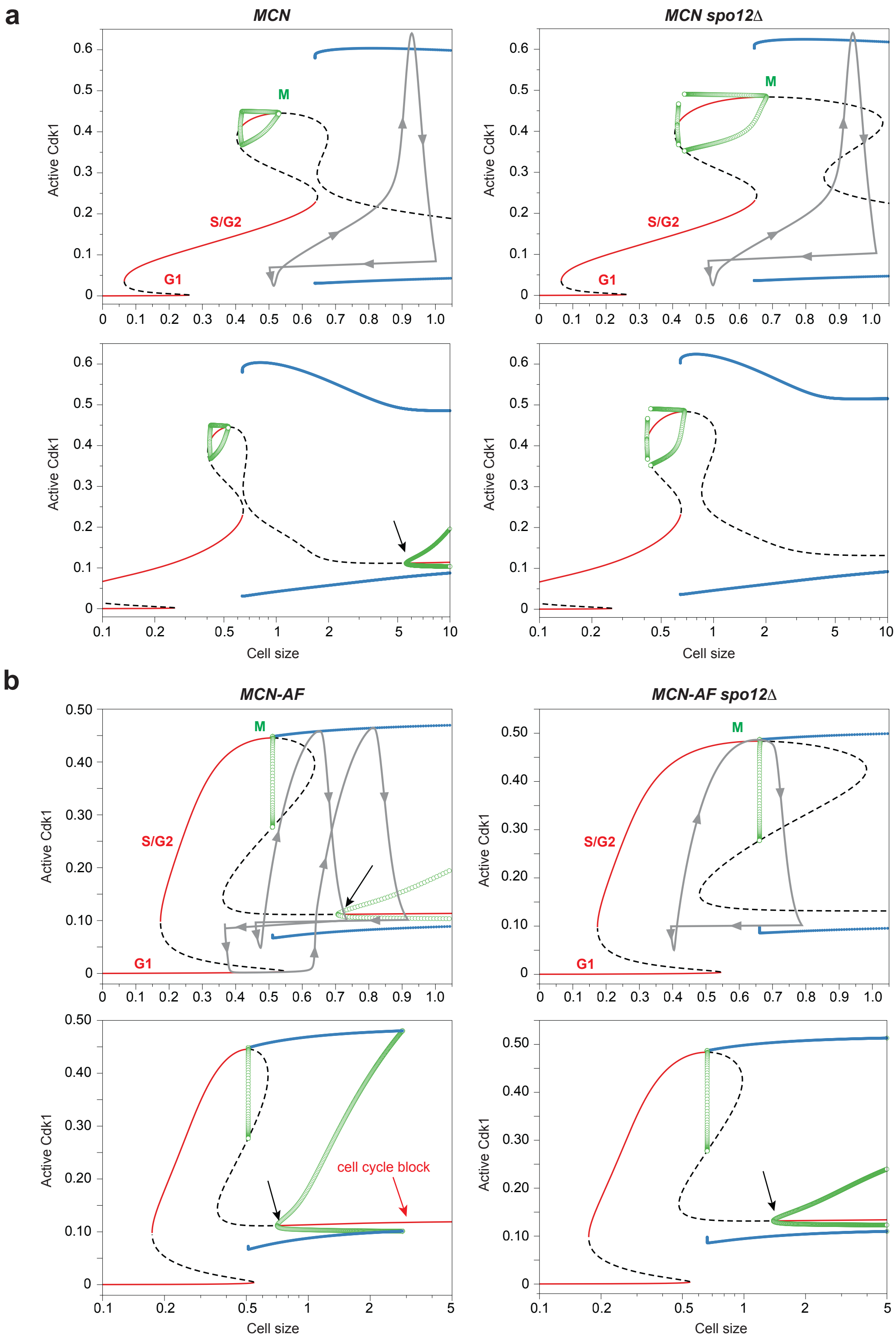

### Extended data Fig. 13

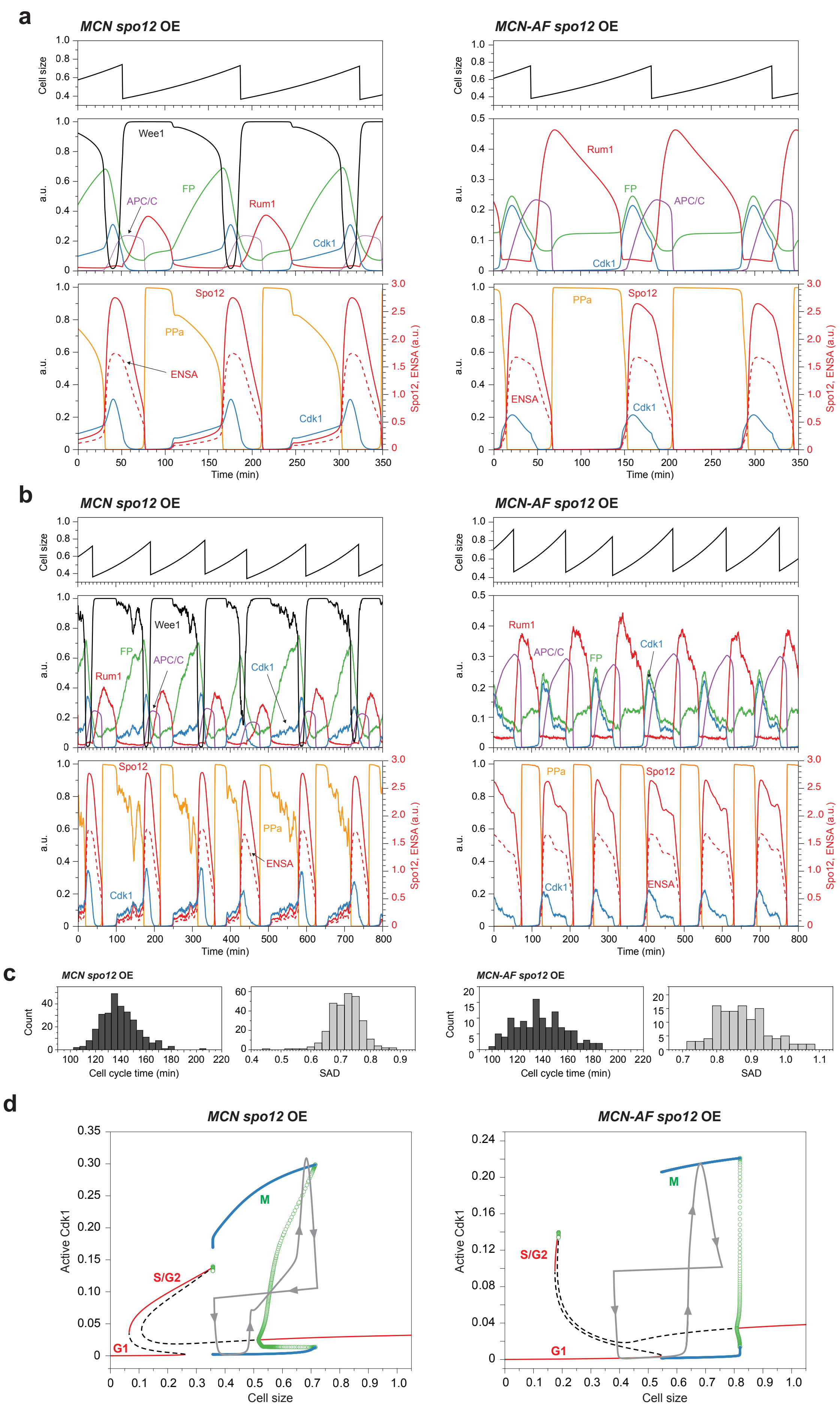

#### Extended data

**Extended data Fig. 1. Experimental evolution of *MCN-AF* populations.**
**a.** Schematic of the laboratory evolution experiment. A single clone from the ancestral strain of each genotype was grown and the culture was split in separate replicates ($M_{\#}$). All individual cultures were then maintained in vegetative growth in a turbidostat system for 70 days[25]. Populations and individual clones were characterized (check marks) and/or sequenced (DNA cartoons) at the indicated times. *: only 2 clones of the wild-type (WT) M7 culture were sequenced at Day 70. **b.** Doubling times for three individual clones of each of the indicated cultures isolated after 35 and 70 days of experimental evolution. At each time point, samples of the cultures were plated on solid medium, and three individual colonies were isolated for characterization. Generation times were determined for each clone in batch cultures. All isolated *MCN-AF* evolved clones showed reduced generation times. **c.** DNA content analysis of the indicated populations after 21 (D21) and 65 (D65) days of experimental evolution. All evolved *MCN-AF* populations showed a reduction in G1 duration, as shown by the reduction in the 1C peak. **d.** DNA content analysis of individual clones of the indicated populations as in *b*. Consistent with the population analyses in *c*, all *MCN-AF* clones showed a reduction in G1 duration. **e.** Size at division (SAD) and coefficient of variation (CV SAD) for the different evolving cultures at the indicated times (as in *c*). SAD was determined from batch cultures that were inoculated with samples from the evolving populations. Box and whiskers plots indicate the min, Q1, Q3 and max, with outliers determined the 1.5 IQR (interquartile range). $n$>100 for each measurement. In all evolving *MCN-AF* cultures, average SAD decreased, with a reduction in CV SAD in M1 and M3. **f.** SAD and CV SAD for three individual clones of the indicated populations as in *b*, *d*. Data are consistent with the population analyses in *e*. *b-f*: Par.: parental strain of the indicated genotypes, prior to evolution. **g.** Muller diagram showing the temporal clonal dynamics in five evolved populations (see Methods). Different clones are color-coded, with clonal specific mutations of interest further indicated. **h.** Sanger sequencing of *spo12* for the different colonies from a representative tetrad obtained by crossing a clone from the final M3 culture (day 70) with the parental *MCN-AF* strain. The population generation time (GT) for each of the four colonies was determined in batch cultures and a 2:2 segregation of the evolved allele (evo) was observed. ref: reference wild-type allele.

**Extended data Fig. 2. Population and single-cell characterization of strains lacking Spo12 function.**
**a.** Blankophor images of WT, *MCN* and *MCN-AF* cells expressing the indicated *spo12* alleles. Scale bar = 10 μm. **b.** Distribution of cell size at division for the indicated genotypes as in Fig. 1e. **c.** Evaluation of meiotic progression in *pat1-114 vs. pat1-114 spo12Δ* diploid cells. The *pat1-114* allele allows for synchronous entry in meiosis at the restrictive temperature of 34 °C[26] (see Methods). nuc.: nucleus; defect: aberrant nuclear divisions. Averages of three independent replicates with standard errors are shown ($n$=100 for each replicate). Loss of Spo12 had no effect on meiotic progression. **d.** Spot assay (serial dilution 1/5) of the indicated genotypes on rich medium (YE4S) and rich medium containing 3 mM hydroxyurea (HU) or 2.5 μM camptothecin (CPT). Loss of Spo12 (*spo12Δ* or *spo12(S54A)*) partially rescues the hypersensitivity of *MCN-AF* cells to replication stress[27]. *rad3Δ* cells are checkpoint-defective and used as a control. **e.** Distribution of cell size at division for the indicated genotypes from single-cell analyses as in Fig. 1g. **f.** Single-cell analyses as in Fig. 1g for the duration of mitosis (i), residency time of Plo1-GFP at the spindle pole body (SPB, ii) and duration of chromosome segregation (iii). The duration of mitosis was defined as the time between the initial recruitment of Plo1 at the SPB and the full segregation of the chromosomes. The residency time of Plo1 at the SPB was defined as the time between its initial recruitment and its disappearance from the duplicated SPBs. The duration of chromosome segregation was defined as the time between the loss of enriched Plo1 signal at the duplicated SPBs and the complete separation of the DNA into two masses (monitored using Nhp6-mCherry). Pooled datasets from three independent replicates. $n$>30 for each replicate. **g.** Western blot analysis of 3HA-tagged (N-terminal) Spo12 and Spo12(S54A) in the indicated genetic backgrounds. Total protein samples were prepared from asynchronous cultures. Tubulin was used as a loading control. Changes in the intensity of the higher molecular weight band between the Spo12 and Spo12(S54A) samples suggest that it comprises the S54-phosphorylated form. The

persistence of a very weak band in the S54A context indicates that Spo12 may be modified on other residues.

**Extended data Fig. 3. Cdc2-dependent phosphorylation of Spo12 regulates Spo12 localization.**
**a.** Localization of msGPF2-Spo12 or msGFP2-Spo12(S54A) in asynchronous cultures of WT, *MCN* and *MCN-AF* cells. An enrichment in msGFP2-Spo12 but not msGFP2-Spo12(S54A) was detected in the nucleus of long cells prior to their division (white arrowheads). Scale bar = 10 μm. **b.** Synchronization protocol for *MCN*-derived cells. Exponentially growing cells were treated with 1 μM of the non-hydrolysable ATP analog 3-MBPP1 for 2 h 45 min to inhibit the MCN fusion protein and block cells in G2[27]. At T=0, cultures were washed three times by filtration to trigger synchronous re-entry into the cell cycle. **c, d.** The synchronization was evaluated using binucleated count (*c*, DAPI staining of heat fixed samples) and DNA content analysis (*d*). In both strains, a strong level of synchrony was observed, with similar peaks in binucleated count and timing of S phase onset (widening of the DNA profiles at T=30 min). B: G2 block. Async: control asynchronous cultures. **e.** Western blot analysis of msGPF2-Spo12 or msGFP2-Spo12(S54A) levels in synchronized cultures (see *b-d*). Ctl: control cells expressing neither of the GFP-tagged versions of Spo12. B: G2 block. Tubulin (Tub) was used as a loading control. An enrichment of the high molecular weight band was noticeable for msGFP2-Spo12 starting at T=15 min, concomitant with M phase (see *c*). This is consistent with the Cdc2-dependent phosphorylation of Spo12 on S54. In *d* and *e*, *spo12*$^{+}$ and *spo12(S54A)* correspond to *MCN msGFP2-spo12* and *MCN msGFP2-spo12(S54A)*, respectively. **f.** *Top panels*: localization of msGPF2-Spo12 or msGFP2-Spo12(S54A) in the synchronous *MCN* populations as in *b-e* (full time course for Fig. 2a). Scale bar = 10 μm. Spo12 but not Spo12(S54A) significantly accumulates in the nucleus at G2/M and becomes diffuse again after mitotic exit. *Bottom panels*: detailed images for cells at T=10 min. A faint signal for msGFP2-Spo12(S54A) was detected at the mitotic spindle (black arrows).

**Extended data Fig. 4. Single-cell analyses of Cdc2 activity and Spo12 overexpression.**
**a.** Representative images of the time-lapse data used to measure the peak nuclear/cytoplasmic signal ratio (N/C) of the synCut3-mCherry biosensor in Fig. 2b. Cells were grown in microfluidic devices with single-cell traps under constant medium flow to allow for optimal growth (see Methods). *Top panel*: for each cell, the nuclear synCut3-mCherry signal was monitored to determine the time of highest intensity. This time point was used to evaluate the peak N/C ratio. *Bottom panel*: DIC images. Scale bars = 10 μm. **b.** Evaluation of *spo12* and *spo12(S54A)* overexpression (OE) in the indicated genetic backgrounds by Western blot analysis. In both cases, an N-terminal 3HA tag allowed for detection of the proteins. Ctl: control strain expressing non-tagged endogenous *spo12*. Tubulin (Tub) was used as a loading control. **c.** Western blot analysis comparing *spo12* and *spo12(S54A)* overexpression as in *b* with their respective endogenous levels. For endogenous protein levels, an N-terminal 3HA tag was integrated at the *spo12* locus in both *spo12* and *spo12(S54A)* strains. Ctl: control strain expressing non-tagged endogenous *spo12*. Tubulin (Tub) was used as a loading control. *: due to the high levels of Spo12 and Spo12(S54A) upon overexpression, 1/10 of the total amount of protein compared to the other samples was loaded on the gel. **d.** Distribution of cell size at division (SAD) and DNA content analysis of cells overexpressing *spo12* or *spo12(S54A)* in the indicated genotypes. Cells were grown as in Fig. 2c in medium with (+T) or without (-T) thiamine. Pooled datasets of three independent replicates ($n$>100 for each replicate). Note that for DNA content analysis, the 1C and 2C dashed lines were aligned to the *MCN* profiles. The WT profiles are shifted to the left due to the smaller size of the cells. The strong cell cycle and morphological defects in *MCN-AF* cells overexpressing *spo12* make it difficult to interpret the corresponding profiles. The difference in the *MCN-AF* profiles in the presence of thiamine compared to Fig. 1f may result from the presence of the vitamin, which is known to enhance growth in fission yeast (compare *MCN-AF spo12(S54A)* OE in the presence and absence of thiamine).

**Extended data Fig. 5. Analysis of Cdc2 activity using a FRET biosensor.**
**a.** Single-cell profiles of FRET/CFP signal and Cdk activity rise times for the indicated genotypes as in Fig. 3a, b. All traces are aligned to the previous division for each cell (T=0 min). WT: $n$=19; *spo12Δ*: $n$=27. Analyses include all measurements, even for cells that did not undergo a full cell cycle over the

duration of the time-lapse experiments. For the activity rise time, a higher threshold of 1.75 was used compared to Fig. 3b due to the difference in the low activity baseline (T=0 to ~50 min). Box and whiskers plots indicate the min, Q1, Q3 and max, with outliers determined by 1.5 IQR (interquartile range). Coefficients of variation are indicated. **b.** Analysis of the single-cell cycle times in the FRET experiments as in *a* and Fig. 3a for the indicated genotypes. Cycle times were determined using successive nuclear divisions as start and end points. For these analyses, only cells that completed a full cell cycle over the duration of the experiments were integrated (WT: *n*=17; *spo12Δ*: *n*=25; *MCN*: *n*=158, *MCN spo12Δ*: *n*=162; *MCN spo12(S54A)*: *n*=142; *MCN-AF*: *n*=125; *MCN-AF spo12Δ*: *n*=143; *MCN-AF spo12(S54A)*: *n*=73). Thus, cells that were either not dividing (Fig. 1h) or that showed highly extended cycle times are not accounted for in this plot. Therefore, the presented averaged cell cycle times do not represent the population doubling times as shown in Fig. 1e. Also note that these experiments were performed at 30 °C, in contrast to the data presented in Figs. 1, 2, 5 and Extended data Figs. 2, 3, 4 and 6 (32 °C), leading to slower cell cycle times. Numbers are coefficients of variation. Box and whiskers plots indicate the min, Q1, Q3 and max, with outliers determined by 1.5 IQR (interquartile range).

**Extended data Fig. 6. Proteomic analysis for Spo12 interactors and phosphoproteomics upon Pab1 depletion.**
**a.** Cell size at division (SAD), population generation time (GT), and DNA content analyses comparing wild-type cells expressing different alleles of *spo12* and their associated N-terminal TurboID (TID) fusions. Spo12 is 3HA-tagged in all strains. Averages of three independent experiments with standard errors. For SAD, *n*>100 for each replicate. Standard deviations of the pooled datasets for SAD: WT, 1.0; *3HA-spo12*, 1.1; *3HA-spo12(S54A)*, 1.1; *3HA-TID-spo12*, 1.1; *3HA-TID-spo12(S54A)*, 1.1. **b.** *Inset panel*: strain numbering for the bottom and right panels. *Bottom left panel*: Western blot analysis of Spo12 in whole cell extracts from the indicated strains. Tubulin (Tub) was used as a loading control. *Right panel*: representative samples used for mass spectrometry (4 and 5) in the TurboID assay, with wild type as a control (all samples were processed using the protocol for mass spectrometry; see Methods). One third of each sample was analyzed by Western blot. Biotinylated proteins were detected using streptavidin-HRP (top blot). The membrane was then stripped and blotted with an anti-HA antibody. Expression of TurboID fusions significantly increased the pull-down of biotinylated proteins. **c.** Schematic of the experimental setup used for Fig. 5a. *AID-pab1* and control strains were treated with 15 µM thiamine (Thi) and 0.5 mM NAA for the indicated times. Following protein extraction and digestion, samples were labeled with isobaric mass tags (TMTplex), each represented by a different color. After mixing, phosphopeptides were enriched and analyzed by liquid chromatography-tandem mass spectrometry (LC-MS/MS). *Inset panel*: depletion of Pab1 was confirmed by Western blot against its V5 tag. Cdc2 (PSTAIR) was used as a loading control. **d.** Heatmap of the phosphopeptides ($n$ = 1479) that showed significant changes upon depletion of Pab1 (using the AID-Pab1 system) compared to the control strain (see *c*), as determined by linear regression ($p<0.05$; see Supplementary Table 3). Phosphopeptide intensities were normalized by row-wise $z$-scoring and grouped into four clusters (C1 to C4) using hierarchical clustering. Color scale represents relative abundance (red = higher, blue = lower). Columns represent conditions (time in the presence of NAA and thiamine) in control or *AID-pab1* strains. **e.** Volcano plot showing phosphopeptides identified in the experiment in *c*, *d*. Phosphopeptide intensities at each timepoint in *AID-pab1* and control strains were $\log_2$-transformed and median-centered. Significant changes were determined by linear regression and comparison of the slopes for each phosphopeptide in the two strains. The $x$-axis represents the effect of Pab1 depletion (interaction slope) and the $y$-axis shows the statistical significance ($-\log_{10}(p\text{-value})$). Also see Supplementary Table 3. Spo12 pS54 and pS47 are highlighted. Pab1 depletion only significantly alters the phosphorylation status of S54. **f.** Kinetic profiles for all the clusters as in *d* (also see Fig. 5a for Cluster 3), as identified by phosphoproteomics upon Pab1 depletion (see *c*). Individual phosphopeptides are shown as black lines, with Spo12(pS54) highlighted in red. The $x$-axis represents time (h) after thiamine + NAA treatment in control or *AID-pab1* strains. **g.** Heatmap of GO terms and other enriched pathways for each of the relevant clusters in *d*. Terms within each cluster are listed in Supplementary Table 3.

**Extended data Fig. 7. Prediction of Spo12 structure.**
**a.** Analysis of Spo12 structure related to Fig. 4b. The conservation of residues was calculated by using the ConSurf online server (https://consurf.tau.ac.il), visualized on the sequence and predicted tertiary structure (AlphaFold Protein Structure Database, accession number AF-Q10189-F1-v6). The highest stretch of conserved residues is indicated by a yellow line above the protein sequence. The location of the phosphorylated serine (S54) is indicated on the predicted structure (right panel). **b, c.** Secondary structure prediction using JPred4 (http://www.compbio.dundee.ac.uk/jpred4)[28] and IUPred3 (https://iupred3.elte.hu/)[29]. The stretch of conserved residues is indicated by a yellow line above the protein sequence. In *b*, the box highlights the predictions of helical (H) and strand (E) residues using three approaches (Jnet, Jhhm, Jpssm). Other results indicate predicted buried residues (B) and overall confidence (from 0 to 9). In *c*, the top line again shows predicted helical regions. The second line illustrates an overall prediction of solvent accessible residues. The boxed region highlights an overall prediction that full-length *S. pombe* Spo12 is disordered.

**Extended data Fig. 8. Structural prediction of the Spo12:PP2A complex.**
**a.** AlphaFold3 structure prediction as in Fig. 4c of the PP2A phosphatase module in complex with unmodified Spo12. **b, c.** Output from the FoldSeek server[30] using the multimer option (https://search.foldseek.com/multimer) in order to find the closest multi-subunit structural homologs from the PDB. The initial search using the AlphaFold3 model of the PP2A phosphatase module in complex with unmodified Spo12 (*b*) identified PDB entries 8ttb (ARPP19) and 8so0 (FAM122A) as the closest structures, but with only very minor similarity for Spo12. When the predicted model included S54-phosphorylated Spo12 (*c*), there was a significant increase in the extent of similarity, with PDB entry 8so0 being the most similar. **d, e.** Structural comparison of the PP2A module in complex with Spo12(pS54) or Igo1 (Alphafold3 models) with PDB entries 8so0, 8ttb[31]. Despite some similarities, the structure of the Spo12:PP2A complex displays some differences with those integrating FAM122A (PDB 8so0) or ARPP19 (PDB 8ttb) (*d*). The AlphaFold3 model of the *S. pombe* PP2A phosphatase complex with IgoI (*e*) clearly belongs to the ARPP19 family based on similarity to PDB 8ttb. **f.** The similarity between phosphatase-bound human ARPP19 and *S. pombe* Igo1 is further supported by using an AlphaFold3 predicted model of the full-length ARPP19. The white circles in *e* and *f* highlight a group of three helical regions and connecting loops of Igo1 and ARPP19 that have a similar conformation in both complexes.

**Extended data Fig. 9. Structural prediction of the *S. cerevisiae* Spo12:PP2A complex.**
**a.** Similar to *S. pombe* Spo12 (Fig. 4b), the AlphaFold2 model of *S. cerevisiae* Spo12 (AlphaFold Protein Structure Database, accession number AF-P17123-F1-v6) is predicted to be intrinsically disordered, with helical regions predicted in the proximity of the S125 phosphosite. **b**. Pairwise sequence alignment between *S. pombe* and *S. cerevisiae* Spo12 using LALIGN within the EMBL-EBI Job Dispatcher[32] shows moderate similarity in the Spo12 sequences, although the two S-P phosphorylation motifs are fully conserved (highlighted in pink). The more stringent EMBOSS_MATCHER identifies a shorter region of similarity between the two Spo12 sequences, but nevertheless includes the two S-P motifs (in pink) and in total correlates with the *S. pombe* Spo12 conserved region (yellow line in Extended data Fig. 7). **c-e.** AlphaFold3 structure predictions of the *S. cerevisiae* PP2A phosphatase module (*c*), in complex with unmodified Spo12 (*d*), and with S125-phosphorylated Spo12 (*e*). For simplicity, Cdc55, Pph22 and Tpd3 (see Fig. 4e) are referred to as scPab1, scPpa2 and scPaa1, respectively. Each panel includes the pLDDT scores (colored as in *a*), the predicted aligned error (PAE), and a surface model of the predicted complex with Spo12 shown as a yellow cartoon. The conserved Spo12 region is shown as a yellow box on the PAE panels.

**Extended data Fig. 10. Role of Spo12 phosphorylation in the Spo12:PP2A complex.**
**a.** From the AlphaFold3 prediction of the phosphatase-bound *S. pombe* Spo12, the negative charge of the main Cdk-phosphorylated serine (pS54) is predicted to counter the positive charges of the nearby surfaces of the Ppa2 and Pab1 phosphatase subunits. Interestingly, the second putative phosphorylation site (S47) is precisely located in the vicinity of the Ppa2 phosphatase active site. **b.** An AlphaFold3 prediction of the phosphatase-bound doubly phosphorylated Spo12 (pS47, pS54) shows an overall

increased confidence in the ipTM and pTM scores (see Supplementary Table 5) and a specific increase in confidence for Spo12 within the PAE plot. **c.** The phosphate of pS47 in the doubly phosphorylated Spo12 is located within the active site of Ppa2, in close proximity to the catalytic $Mn^{2+}$ ions. **d.** AlphaFold3 was also used to predict the structure of *S. cerevisiae* phosphatase-bound Spo12 with double phosphorylation on S118 and S125. For simplicity, Cdc55, Pph22 and Tpd3 (see Fig. 4e) are referred to as scPab1, scPpa2 and scPaa1, respectively. **e**. As in fission yeast (*c*), this second phosphorylated serine in *S. cerevisiae* Spo12, seven residues N-terminal to the main phosphorylation site, is placed within the active site, beside the catalytic $Mn^{2+}$ ions. The second phosphorylation could therefore help in Spo12 binding, but would also be directly dephosphorylated by the phosphatase subunit.

**Extended data Fig. 11. Stochastic simulations of a model cell cycle network integrating Spo12 function.**
**a, b.** Stochastic simulations of the model in Fig. 6a, corresponding to the deterministic simulations in Fig. 6b, c. Noise was introduced to the ODEs describing the accumulation of the fusion protein (Cdk1 and cyclin B) as well as that of the G1 inhibitor Rum1. See Supplementary Model, part C. a.u.: arbitrary unit. **c.** Distribution of cell size at division (SAD), corresponding to the cell cycle time distribution in Fig. 6d, for the indicated genotypes using stochastic simulations as in *a*. $n$>100 for each genotype. **d.** Percentages of cells that show a late Cdk activity rise time (>50 min) in the simulated single-cell profiles in Fig. 6e. The model predictions are consistent with the FRET observations, although the reduction in the fraction of cells with a long G1 is more pronounced in the experimental setup (compare Fig. 3a with Fig. 6e). **e.** Simulated Cdk activity rise times in single cells from the data in Fig. 6e, restricting the analysis to *MCN-AF* and *MCN-AF spo12Δ* cells that show a long G1 (late rise time in *d;* $n$=128 for *MCN-AF* and $n$=76 for *MCN-AF spo12Δ*). CV: coefficient of variation. Even when only considering cells with a long G1, the model predicts an increase in cell-to-cell homogeneity in Cdk activity profiles upon loss of Spo12 function. Box and whiskers plots indicate the min, Q1, Q3 and max, with outliers determined by 1.5 IQR (interquartile range). ***: $p$<0.001 (two-tailed independent T-test for normal distribution).

**Extended data Fig. 12. Bifurcation diagrams of the cell cycle model.**
**a, b.** Bifurcation diagrams for the deterministic model of the indicated genotypes (see Supplementary Model, part B) as in Fig. 6b, c, using cell size as a bifurcation parameter (with *MCN* cells dividing at a size of 1). Red: stable steady-states (G1, S/G2 and M, resulting from the bistable switches of the model governed by Rum1 and Wee+Cdc25); dashed black: unstable steady-states; blue: amplitude (minima and maxima) of stable oscillations; green: amplitude (minima and maxima) of unstable oscillations; grey lines: overlay of the cell cycle trajectories from the numerical simulations of the deterministic model. Black arrows indicate the onset of a non-canonical steady-state of permanent cell cycle arrest. In the *MCN-AF* strain, this steady-state first appears in cells of size ≈ 0.7 a.u., compared to ≈ 6 a.u. in *MCN* cells. Note that in contrast to *MCN*, the stable limit cycles end at size ≈ 3 in *MCN-AF* cells, leading to cell being attracted to the arrested steady-state (red arrow, cell cycle block). *a,b*: bottom panels are similar to the top panels but extended over a broader range of cell sizes (log scale *x*-axis).

**Extended data Fig. 13. Simulations of the model cell cycle network upon *spo12* overexpression.**
**a, b.** Deterministic (*a*) and stochastic (*b*) simulations based on the model in Fig. 6a, in *MCN* or *MCN-AF* cells overexpressing *spo12* (OE). For the stochastic simulations, noise was introduced as in Extended data Fig. 11. a.u.: arbitrary unit. **c.** Distribution of cell cycle time and size at division (SAD) for the indicated genotypes from the stochastic simulations in *b*. $n$>100 for each genotype. **d.** Bifurcation diagrams as in Extended data Fig. 12 for the *MCN* and *MCN-AF* cells overexpressing *spo12*.
