## Supplementary Model for "Regulation of cell proliferation by a novel feedback system on Cdk function"

### A. Deterministic Model

The mathematical model of Spo12 function in *MCN* cells is based on the proposed molecular mechanism in Fig. SM1 (also see Fig. 6a). The variables, parameters and other definitions used to define the model are introduced in Table SM1. Using standard principles of biochemical kinetics, we define the deterministic model by the set of nonlinear ordinary differential equations (ODEs) provided in Table SM2. We simulate *MCN* cells using the parameter values specified in Table SM3.

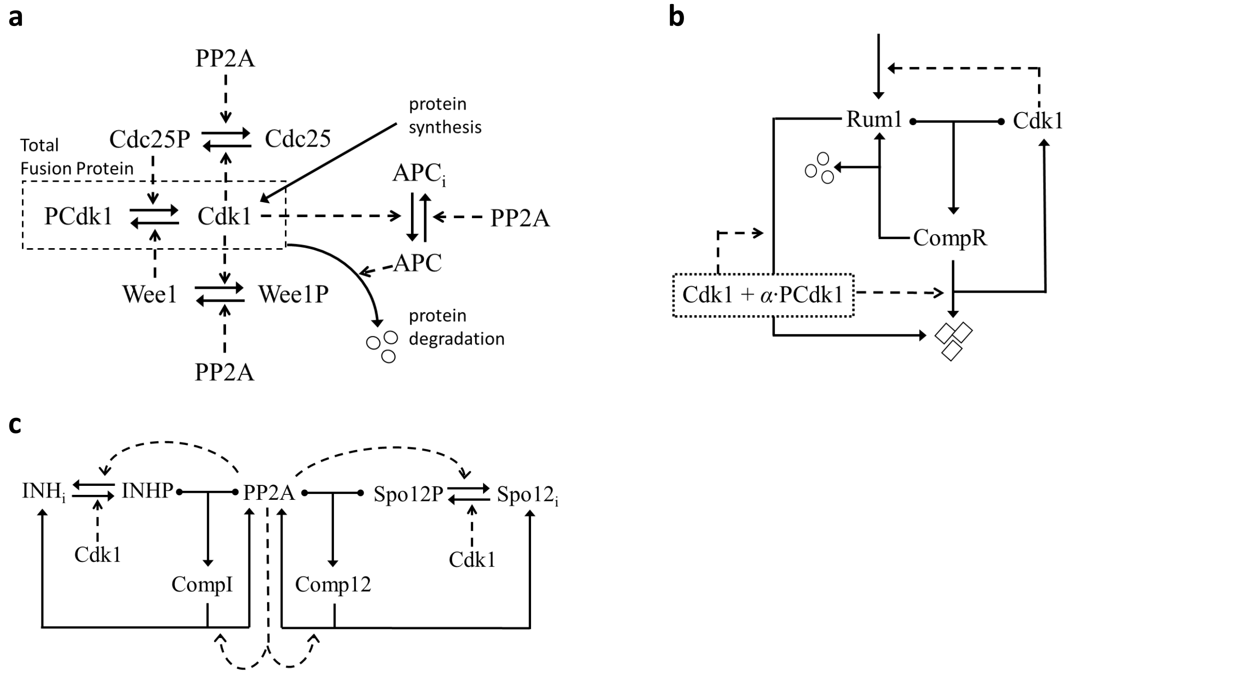


**Fig. SM1.** Proposed mechanism for cell cycle regulation in *MCN* fission yeast cells. **a.** Regulation of the fusion protein, Cdk1-Cdc13, which (we assume) is synthesized at a constant rate. The fusion protein exists in two forms: unphosphorylated active 'Cdk1' and tyrosine-phosphorylated inactive 'PCdk1'. The tyrosine kinase, Wee1, is inactivated by phosphorylation by Cdk1, whereas the tyrosine phosphatase, Cdc25, is activated by phosphorylation by Cdk1. We assume that PP2A is the counteracting phosphatase. Meanwhile, Cdk1 activates the Anaphase Promoting Complex/Cyclosome, 'APC', which ubiquitinates the fusion protein, labelling it for degradation by proteasomes. **b.** Regulation of the fusion protein by a stoichiometric inhibitor, Rum1. Rum1 binds to active Cdk1-Cdc13 to form an inactive complex, 'CompR'. We assume that Rum1 does not bind to PCdk1-Cdc13). Degradation of the fusion protein from CompR releases free Rum1. Rum1 phosphorylation by Cdk1 (and to a lesser extent by PCdk1) labels Rum1 for ubiquitination and proteolysis, releasing active Cdk1 from CompR. Active Cdk1 is assumed to upregulate Rum1 synthesis. **c.** Regulation of PP2A by stoichiometric inhibitors. 'INH' represents Igo1/ENSA and any other unspecified inhibitors. INHP and Spo12P are the phosphorylated, active forms of the inhibitors.

**Table SM1.** Definitions of Variables, Parameters and Other Quantities Governed by the Model. FP = fusion protein.

| | | | |
|---|---|---|---|
| Variables: | Cdk1 | = | kinase activity of FP (unphosphorylated Cdk1-Cdc13) |
| | PCdk1 | = | tyrosine-phosphorylated FP (PCdk1-Cdc13) |
| | Wee1, Cdc25 | = | tyrosine kinase and phosphatase (unphosphoryl'd forms) |
| | Rum1 | = | stoichiometric inhibitor of FP |
| | APC | = | Anaphase Promoting Complex (E3 ubiquitin ligase) |
| | PP2A | = | Cdk1-counteracting phosphatase (Ppa1/2:Pab1) |
| | Spo12, INH | = | stoichiometric inhibitors of PP2A |
| | CompR | = | Rum1:Cdk1-Cdc13 complex |
| | CompI, Comp12 | = | INHP:PP2A, Spo12P:PP2A complexes |
| | Size | = | Cell size (volume, mass, protein content, etc.) |
| Parameters: | $k_{s...}$, $k_{d...}$ | = | rate const for synthesis, degradation of protein … |
| | $k_{as...}$, $k_{di...}$ | = | rate const for association, dissociation of complex … |
| | $k_{ph...}$, $k_{dp...}$ | = | rate const for phosphorylation, dephos'n of protein … |
| | $K_{mapc}$ | = | Michaelis const for phos'n and dephos'n of APC |
| | $J$ | = | threshold for Cdk1 activation of Rum1 synthesis |
| | $PP2A_T$, $Spo12_T$, $INH_T$ | = | const total concentrations of PP2A, Spo12 and INH |
| | $Wee1_T$, $Cdc25_T$, $APC_T$ | = | const total concentrations of Wee1, Cdc25 and APC |
| | $\mu$ | = | specific growth rate of cells |
| Composite Variables: | $CycB_T$ | = | Cdk1 + PCdk1 + CompR (total Cdc13) |
| | $Rum1_T$ | = | Rum1 + CompR (total Rum1) |
| | $INH_{aT}$ | = | INHP + CompI (total phosphorylated INH) |
| | $Spo12_{aT}$ | = | Spo12P + Comp12 (total phosphorylated Spo12) |
| Derived Variables: | CompR | = | $CycB_T$ – Cdk1 – PCdk1 |
| | CompI | = | $PP2A_T$ – PP2A – Comp12 |
| | $INH_i$ | = | $INH_T$ – $INH_{aT}$ $Spo12_i$ = $Spo12_T$ – $Spo12_{aT}$ |
| | Wee1P | = | $Wee1_T$ – Wee1 Cdc25P = $Cdc25_T$ – Cdc25 |
| | $APC_i$ | = | $APC_T$ – APC |

**Table SM2.** Defining Equations of the Deterministic Model.

$$\frac{d}{dt}CycB_{\mathrm{T}} = k_{\mathrm{sfp}} - \left(k'_{\mathrm{dfp}} + k_{\mathrm{dfp}} * APC\right) * CycB_{\mathrm{T}} - k''_{\mathrm{dfp}} * CompR$$

$$\frac{d}{dt}Cdk1 = k_{\mathrm{sfp}} - \left(k'_{\mathrm{dfp}} + k_{\mathrm{dfp}} * APC\right) * Cdk1 - k_{\mathrm{asR}} * (Rum1_{\mathrm{T}} - CompR) * Cdk1 + k_{\mathrm{diR}} * CompR$$
$$+ \left(k'_{\mathrm{drum}} + k_{\mathrm{drum}} * (Cdk1 + \alpha * PCdk1)\right) * CompR - V_{\mathrm{wee}} * Cdk1 + V_{\mathrm{c25}} * PCdk1$$

$$\frac{d}{dt}PCdk1 = V_{\mathrm{wee}} * Cdk1 - V_{\mathrm{c25}} * PCdk1 - \left(k'_{\mathrm{dfp}} + k_{\mathrm{dfp}} * APC\right) * PCdk1$$

$$\frac{d}{dt}Rum1_{\mathrm{T}} = V_{\mathrm{srum}} - \left(k'_{\mathrm{drum}} + k_{\mathrm{drum}} * (Cdk1 + \alpha * PCdk1)\right) * Rum1_{\mathrm{T}}$$

$$\frac{d}{dt}APC = \frac{k_{\mathrm{phapc}} * Cdk1 * (APC_{\mathrm{T}} - APC)}{K_{\mathrm{mapc}} + APC_{\mathrm{T}} - APC} - \frac{k_{\mathrm{dpapc}} * PP2A * APC}{K_{\mathrm{mapc}} + APC}$$

$$\frac{d}{dt}PP2A = -k_{\mathrm{asI}} * (INH_{\mathrm{aT}} - CompI) * PP2A + k_{\mathrm{diI}} * CompI - k_{\mathrm{asS}} * (Spo12_{\mathrm{at}} - CompS) * PP2A$$
$$+ k_{\mathrm{diS}} * CompS + \left(k'_{\mathrm{dpinh}} + k_{\mathrm{dpinh}} * PP2A\right) * CompI + \left(k'_{\mathrm{dpspo}} + k_{\mathrm{dpspo}} * PP2A\right)$$
$$* CompS$$

$$\frac{d}{dt}INH_{\mathrm{aT}} = k_{\mathrm{phinh}} * Cdk1 * (INH_{\mathrm{T}} - INH_{\mathrm{aT}}) - \left(k'_{\mathrm{dpinh}} + k_{\mathrm{dpinh}} * PP2A\right) * INH_{\mathrm{aT}}$$

$$\frac{d}{dt}Spo12_{\mathrm{aT}} = k_{\mathrm{phspo}} * Cdk1 * (Spo12_{\mathrm{T}} - Spo12_{\mathrm{aT}}) - \left(k'_{\mathrm{dpspo}} + k_{\mathrm{dpspo}} * PP2A\right) * Spo12_{\mathrm{aT}}$$

$$\frac{d}{dt}CompS = k_{\mathrm{asS}} * (Spo12_{\mathrm{aT}} - CompS) * PP2A - k_{\mathrm{diS}} * CompS$$
$$- \left(k'_{\mathrm{dpspo}} + k_{\mathrm{dpspo}} * PP2A\right) * Comp12$$

$\frac{d}{dt}Size = \mu * Size,$ *Size* → *Size*/2 when *Cdk1* drops below *MET*, 'mitotic exit threshold'

Definitions: $V_{\mathrm{wee}} = k'_{\mathrm{wee}} * Wee1P + k_{wee} * Wee1$

$$V_{\mathrm{c25}} = (k'_{\mathrm{c25}} * Cdc25 + k_{c25} * Cdc25P) \cdot Size$$

$$V_{\mathrm{srum}} = \left(k'_{\mathrm{srum}} + k_{\mathrm{srum}} \frac{Cdk1^r}{J^r + Cdk1^r}\right) \cdot \frac{1}{Size}$$

$$\frac{Wee1}{Wee1_{\mathrm{T}}} = \frac{1 - \left(\frac{Cdk1}{k' + k_{\mathrm{dpwee}} * PP2A}\right)^{1+Qw}}{1 - \left(\frac{Cdk1}{k' + k_{\mathrm{dpwee}} * PP2A}\right)^{1+Nw}} \; ; \; \frac{Cdc25}{Cdc25_{\mathrm{T}}} = \frac{1 - \left(\frac{Cdk1}{k' + k_{\mathrm{dpc25}} * PP2A}\right)^{1+Q25}}{1 - \left(\frac{Cdk1}{k' + k_{\mathrm{dpc25}} * PP2A}\right)^{1+N25}}$$

**Table SM3.** Parameter Values for Simulating *MCN* cell cycles.*

| | | | |
|---|---|---|---|
| $k_{sfp} = 0.02$ | $k'_{dfp} = 0.015$ | $k_{dfp} = 0.65$ | $k''_{dfp} = 0.15$ |
| $k_{asR} = 100$ | $k_{diR} = 0.1$ | $k'_{drum} = 0.125$ | $k_{drum} = 21$ |
| $k_{phapc} = 0.1$ | $k_{dpapc} = 3$ | $K_{mapc} = 1$ | $\alpha = 0.25$ |
| $k'_{wee} = 0.001$ | $k_{wee} = 0.26$ | $k'_{c25} = 0.05$ | $k_{c25} = 0.2$ |
| $k'_{srum} = 0.03$ | $k_{srum} = 0.4$ | $J = 0.4$ | $r = 2$ |
| $k_{dpwee} = 0.2$ | $Q_w = 1$ | $N_w = 10$ | $k' = 0.21$ |
| $k_{dpc25} = 0.02$ | $Q_{25} = 8$ | $N_{25} = 10$ | |
| $k_{asI} = 25$ | $k_{diI} = 0.025$ | $k_{phinh} = 2$ | $k'_{dpinh} = 0.05$ |
| $k_{asS} = 25$ | $k_{diS} = 0.025$ | $k_{phspo} = 2$ | $k'_{dpspo} = 0.05$ |
| $k_{dpinh} = 4$ | $Wee1_T = 1$ | $Cdc25_T = 1$ | $APC_T = 1$ |
| $k_{dpspo} = 4$ | $PP2A_T = 1$ | $Spo12_T = 0.1$ | $INH_T = 1.9$ |
| $\mu = 0.005$ | $MET = 0.1$ | | |

* Time in min, all concentrations in arbitrary units.

The differential equations in Table SM2 are derived directly from the molecular mechanism in Fig. SM1, except for the tyrosine-modifying enzymes, Wee1 and Cdc25. We assume that these enzymes are regulated by mechanisms of ordered, multisite phosphorylation, mediated by Cdk1. Provided these reactions are fast enough that the distribution of phosphorylated forms is always at pseudo-steady state, then the fraction of protein that is hypo-phosphorylated ($\leq Q$ phosphorylated sites) is given by the equation[1]:

$$\frac{X + XP + XP_2 + \cdots + XP_Q}{X_T} = \frac{1 - \left(\frac{k}{h}\right)^{1+Q}}{1 - \left(\frac{k}{h}\right)^{1+N}}$$

where $k$ = activity of the kinase, $h$ = activity of the counteracting phosphatase, and $N$ = total number of phospho-sites on the protein. We assume that $N = 10$ for both Wee1 and Cdc25, that Wee1 and Wee1P are active but higher order phospho-forms are inactive, and that only Cdc25P$_9$ and Cdc25P$_{10}$ are active. Notice, as well, that cell size grows exponentially, with a mass-doubling time = $\ln 2/\mu$, and a mother cell divides precisely in half when Cdk1 activity drops below a threshold, *MET*, as the cell exits mitosis.

Parameter values are assigned (by hand) to provide a reasonable fit of model simulations to the observed behaviors of *MCN* cells and derived mutant genotypes. Notice, in particular, that the parameter values determining the dynamical properties of the PP2A inhibitors, INH and Spo12, are identical; this is our default assumption in the absence of any experimental evidence to the contrary. Notice also, that Spo12 represents only 5% of the total inhibitor concentration ($INH_T$ + $Spo12_T$) in *MCN* cells.

An 'ode' file, suitable for simulating this deterministic model in the freely available software XPP (XPP/XPPAUT Homepage), is provided in Code SM1. The output of this code, i.e., a simulation of the cell-cycle biochemistry in *MCN* cells, is provided in Fig. 6b of the main text. To simulate the mutant strains in Fig. 6b, c and in Extended data Fig. 13a, we made appropriate changes to specific parameter values and initial conditions:

| Genotype | Parameter Values | | | | | Initial Conditions | | |
|---|---|---|---|---|---|---|---|---|
| | $k'_{wee}$ | $k_{wee}$ | $k'_{c25}$ | $k_{c25}$ | $Spo12_T$ | $Spo12_{aT}$ | *CompS* | *PCdk1* |
| *MCN* | 0.001 | 0.26 | 0.05 | 0.2 | 0.1 | 0.005 | 0.0045 | 0.325 |
| *MCN spo12Δ* | 0.001 | 0.26 | 0.05 | 0.2 | 0 | 0 | 0 | 0.325 |
| *MCN spo12* OE | 0.001 | 0.26 | 0.05 | 0.2 | 3 | 0.15 | 0.12 | 0.325 |
| *MCN-AF* | 0 | 0 | 0 | 0 | 0.1 | 0.005 | 0.0045 | 0 |
| *MCN-AF spo12Δ* | 0 | 0 | 0 | 0 | 0 | 0 | 0 | 0 |
| *MCN-AF spo12* OE | 0 | 0 | 0 | 0 | 3 | 0.15 | 0.12 | 0 |

Notice that, in *spo12* OE strains, $Spo12_T = 3$, which is 30-fold larger than $Spo12_T$ in cells carrying the wild-type gene. This fold increase upon *spo12* OE is an approximation based on the protein levels provided for endogenous Spo12 and Nmt1 (whose promoter was used for *spo12* OE) in[2] (also see www.pombase.org). Furthermore, notice that, in *spo12* OE strains, the peak of Cdk1 activity is only ~0.25, which is less than half the peak activity in the other strains. Because PP2A activity is low in *spo12* overexpressing strains, these cells prematurely activate APC at much lower Cdk1 activity; hence, the accumulation of fusion protein is short-circuited and peak Cdk1 activity is reduced. Considering that the *MCN spo12* OE strain is viable, we conclude that much lower levels of Cdk1 activity are necessary to drive DNA synthesis and mitosis in this strain, presumably because its counteracting phosphatase activity is also much lower.

**B. Bifurcation Diagrams**

A bifurcation diagram illustrates how characteristic solutions of a nonlinear dynamical system (namely, steady states and limit cycle oscillations) depend on the value of a control parameter in the ODEs. In our case (*MCN* cells), because Cdk1 activity alone drives progression through S and M phases, we choose 'Cdk1' as the dynamic variable in the bifurcation diagram. Because growth appears to drive fission yeast through the phases (G1-S-G2-M) of the cell cycle, we choose 'Size' as the control parameter. To this end, we modify the 'deterministic.ode' code to remove Size as a variable and declare it as a parameter:

```
# dSize/dt = mu*Size
par Size=0.6
```

In Extended data Fig. 12 and 13d, we present bifurcation diagrams for each of the six genotypes under consideration. Concentrating on Extended data Fig. 12a, for *MCN* cells, we see a locus of steady state solutions (red solid lines = stable steady states, black dashed lines = unstable steady states) that twists its way from stable steady states of small size and low Cdk1 activity (G1 cells in the lower left corner) to large cells (Size=10) in a stable steady state with Cdk1≈0.12 (right boundary). Along the way, the steady state loses stability at saddle-node (SN) bifurcations at Size≈0.25 and 0.6, and also at Hopf bifurcations (HBs) at Size≈0.4, 0.5 and 6. At each HB there bifurcates a locus of unstable limit cycle oscillations (green open circles). More relevant to us is the locus of stable limit cycles (solid blue curves, which delineate the maximum and minimum excursions of Cdk1 activity during each oscillation). The stable limit cycles arise at a SNIC bifurcation ('saddle-node on an invariant circle') at Size≈0.6, where they are 'born' with large amplitude ($0.25 < Cdk1(t) < 0.57$) and very long (infinite) period. The stable limit cycles persist to Size≫10, where they disappear by coalescing with the locus of unstable limit cycles at a cyclic fold (CF) bifurcation.

The gray curve in Extended data Fig. 12a is the trajectory of size and Cdk1 activity of a growing and dividing fission yeast cell. A newly divided cell, born at Size≈0.5 and Cdk1≈0.05, is attracted to the only stable steady state (the red S/G2 state), but it never gets there because it is steadily increasing in size. Nonetheless, rising Cdk1 activity drives the cell through S phase and into G2. When Size surpasses 0.6, the S/G2 steady state disappears, and the trajectory is attracted to the stable limit cycle oscillation, but the period of the limit cycle is so long (at first) that the trajectory cannot latch onto the limit cycle until Size>0.85. The abrupt rise in Cdk1 activity drives the cell into mitosis, activates APC, and the subsequent degradation of fusion protein causes Cdk1 activity to drop below the threshold for mitotic exit (0.08 in this simulation). The cell divides at Size=1 to produce to newborn cells at Size=0.5, to complete the cycle.

The cell cycle trajectory of *MCN* cells is extremely robust to stochastic fluctuations, but the bifurcation diagram for *MCN-AF* cells (Extended data Fig. 12b) tells a different story. First of all, the deterministic cell-cycle trajectory (the gray curve) exhibits a 'period doubling' bifurcation: a short cycle ($0.45<\text{Size}<0.72$; period = $(140/0.301)\times\log(0.72/0.45) = 95$ min) alternates with a long cycle ($0.36<\text{Size}<0.9$; period = … = 185 min). Second, the rise in Cdk1 activity (max≈0.45) is not so large as for *MCN* cells (max≈0.65). So, if cells fail to enter (or complete) mitosis, they may continue to grow, and if Size exceeds ~3, they will drop off the limit cycle and into the 'cell cycle block' state, with 'indecisive' Cdk1 activity (~0.15) and APC activity (~0.2, not shown). Furthermore, Extended data Figs. 12 and 13d suggest that *MCN-AF spo12Δ* cells should be less prone than *MCN-AF* cells to exiting the cell division cycle, and *MCN-AF spo12* OE cells should be more prone. To estimate just how prevalent these aberrations may be requires a realistic stochastic model of cell cycle progression in *MCN* strains (see C).

**Code SM1.** Spo12model_deterministic.ode

```
# Cdk1 P'n & Rum1 binding are exclusive
# CycBT = total fusion protein = Cdk1 + PCdk1 + Rum1:Cdk1
# Rum1T = Rum1 + Rum1:Cdk1
# APC = active APC, APCtotal = 1
# PP2A = active PP2A, not bound to INH or Spo12
# INHaT = INH + INH:PP2A
# INHT = INHaT + INHi = total INH (adjustable parameter)
# Spo12aT = Spo12 + Spo12:PP2A
# Spo12T = Spo12aT + Spo12i = total Spo12 (adjustable parameter)
# PP2AT = PP2A + INH:PP2A + Spo12:PP2A = 1
# Size = Cell volume, cell mass, total protein content, etc.

# Define ODEs
dCycBT/dt = ksfp - (kdfp'+kdfp*APC)*CycBT - kdfp"*CompR

dCdk1/dt = ksfp - (kdfp'+kdfp*APC)*Cdk1 - Vwee*Cdk1 + Vc25*PCdk1
+ kdiR*CompR - kasR*(Rum1T - CompR)*Cdk1 + (kdrum' +
kdrum*(Cdk1+alpha*PCdk1))*CompR

dPCdk1/dt = Vwee*Cdk1 - Vc25*PCdk1 - (kdfp'+kdfp*APC)*PCdk1

dRum1T/dt = Vsrum - (kdrum' + kdrum*(Cdk1+alpha*PCdk1))*Rum1T

dAPC/dt = kphapc*Cdk1*(1-APC)/(K_mapc+1-APC) -
kdpapc*PP2A*APC/(K_mapc+APC)

dPP2A/dt = -kasI*(INHaT-CompI)*PP2A + kdiI*CompI -
kasS*(Spo12aT-CompS)*PP2A + kdiS*CompS +
(kdpinh'+kdpinh*PP2A)*CompI + (kdpspo'+kdpspo*PP2A)*CompS

dINHaT/dt = kphinh*Cdk1*(INHT-INHaT) -
(kdpinh'+kdpinh*PP2A)*INHaT

dSpo12aT/dt = kphspo*Cdk1*(Spo12T-Spo12aT) -
(kdpspo'+kdpspo*PP2A)*Spo12aT

dCompS/dt = kasS*(Spo12aT-CompS)*PP2A - kdiS*CompS -
(kdpspo'+kdpspo*PP2A)*CompS

dSize/dt = mu*Size

# Activity functions for Wee1 and Cdc25
Wee1 =  (1-(Cdk1/Vawee)^(Qw+1))/(1-(Cdk1/Vawee)^(Nw+1))
Cdc25 = (1-(Cdk1/Vi25)^(Q25+1))/(1-(Cdk1/Vi25)^(N25+1))
```

```
# Cell growth down-regulates Rum1 synthesis and up-regulates
Cdc25 activity
Vsrum = (ksrum'+ksrum*Cdk1^r/(J^r+Cdk1^r))/Size
Vc25  = (kc25'*Cdc25 + kc25*(1-Cdc25))*Size

# Other definitions
CompR = CycBT - Cdk1 - PCdk1
CompI = PP2AT - PP2A  - CompS
Vawee = k' + kdpwee*PP2A
Vi25 =  k' + kdpc25*PP2A
Vwee =  kwee'*(1 - Wee1) + kwee*Wee1

init Size=0.6, CycBT=0.45, Cdk1=0.1, PCdk1=0.325
init Rum1T=0.023, APC=0.0018, PP2A=0.91, INHaT=0.098
init Spo12aT=0.005, CompS=0.0045

# Auxiliary variables for plotting purposes
aux Wee1pt = (1-(Cdk1/Vawee)^(Qw+1))/(1-(Cdk1/Vawee)^(Nw+1))
aux Cdc25P = 1-(1-(Cdk1/Vi25)^(Q25+1))/(1-(Cdk1/Vi25)^(N25+1))
aux  INHpt = INHaT/2
aux Spo12pt = Spo12aT/3

# Cell divides when Cdk1 activity drops below mitotic-exit
threshold
global -1 {Cdk1-MET} {Size=Size/2}

# Parameter values for MCN cell cycles
p ksfp=0.02,   kdfp'=0.015, kdfp=0.65,     kdfp"=0.15
p kasR=100,    kdiR=0.1,    kdrum'=0.125,  kdrum=21
p kphapc=0.1,  kdpapc=3,    K_mapc=1,      alpha=0.25
p kwee'=0.001, kwee=0.26,   kc25'=0.05,    kc25=0.2
p ksrum'=0.03, ksrum=0.4,   J=0.4,         r=2
p kdpwee=0.2,  Qw=1,        Nw=10,         k'=0.21
p kdpc25=0.2,  Q25=8,       N25=10
p kasI=25,     kdiI=0.025,  kphinh=2,      kdpinh'=0.05
p kasS=25,     kdiS=0.025   kphspo=2,      kdpspo'=0.05
p kdpinh=4,    PP2AT=1,     Spo12T=0.1,    INHT=1.9
p kdpspo=4,    mu=0.005,    MET=0.1
# Note:        APCT=1,      Wee1T=1,       Cdc25T=1

# Default settings for XPP
@ meth=stiff, total=1000, dt=1, maxstor=10000
@ xp=time, xlo=0, ylo=-0.03, xhi=1000, yhi=1.03
@ nplot=5, yp=Size, yp2=Rum1T, yp3=APC, yp4=Cdk1, yp5=CycBT
# Plotting variables for lower panel of simulations
#@ nplot=5, yp=Size, yp2=PP2A, yp3=Spo12aT, yp4=Cdk1, yp5=INHpt
Done
```

### C. Stochastic Model

The stochastic version of the model is a set of stochastic differential equations derived from the deterministic model by adding white noise to the ODEs describing the accumulation of fusion protein (Cdk1 and $CycB_T$) and its stoichiometric inhibitor ($Rum1_T$):

$$\frac{d}{dt}CycB_T = k_{sfp} - \left(k'_{dfp} + k_{dfp} * APC\right) * CycB_T - k''_{dfp} * CompR + \sigma\sqrt{CycB_T} \cdot WienB$$

$$\frac{d}{dt}Cdk1 = k_{sfp} - \left(k'_{dfp} + k_{dfp} * APC\right) * Cdk1 - k_{asR} * (Rum1_T - CompR) * Cdk1 + k_{diR} * CompR + \left(k'_{drum} + k_{drum} * (Cdk1 + \alpha * PCdk1)\right) * CompR - V_{wee} * Cdk1 + V_{c25} * PCdk1 + \sigma\sqrt{Cdk1} \cdot WienB$$

$$\frac{d}{dt}Rum1_T = V_{srum} - \left(k'_{drum} + k_{drum} * (Cdk1 + \alpha * PCdk1)\right) * Rum1_T + \sigma\sqrt{Rum1_T} \cdot WienR$$

where $WienB$ and $WienR$ are 'wiener variables', *i.e.*, normally distributed random numbers with mean = 0 and variance = $(dt)^{1/2}$, where $dt$ is the numerical time step in a fixed time-step integrator such as the methods of Euler or Runge-Kutta (see [3], page 98). The parameter $\sigma$ determines the magnitude of stochastic fluctuation; we choose $\sigma = 0.015$.

In addition, because the *Cdk1*($t$) variable is fluctuating, we must introduce a more subtle condition for cell division: first, *Cdk1*($t$) must increase above $M_{entry}$, indicating that the cell has entered mitosis, and then *Cdk1*($t$) must decrease below $M_{exit} < M_{entry}$, to signal cell division. For our six genotypes, we choose $M_{entry}$ and $M_{exit}$ as follows:

| Genotype | $M_{entry}$ | $M_{exit}$ | $Spo12_T$ | $Spo12_{aT}(0)$ | $CompS(0)$ | $PCdk1(0)$ |
|---|---|---|---|---|---|---|
| *MCN* | 0.4 | 0.1 | 0.1 | 0.006 | 0.005 | 0.5 |
| *MCN spo12Δ* | 0.4 | 0.1 | 0 | 0 | 0 | 0.5 |
| *MCN spo12* OE | 0.2 | 0.05 | 3 | 0.18 | 0.16 | 0.5 |
| *MCN-AF* | 0.4 | 0.1 | 0.1 | 0.006 | 0.005 | 0 |
| *MCN-AF spo12Δ* | 0.4 | 0.1 | 0 | 0 | 0 | 0 |
| *MCN-AF spo12* OE | 0.2 | 0.05 | 3 | 0.18 | 0.16 | 0 |

For *spo12* OE strains, $M_{entry}$ and $M_{exit}$ are reduced two-fold, because that activity of PP2A is also much lower in these strains (as previously noted).

The 'ode' file for the stochastic model is provided in Code SM2. Simulations using this code are shown in Extended data Figs. 11a, b and 13b.

**Code SM2.** Spo12model_stochastic.ode

```
# Define ‘wiener variables’ which are normally distributed
# random numbers with mean 0 and variance sqrt(dt)
wiener WienB, WienR

# Define SDEs
dCycBT/dt = ksfp - (kdfp'+kdfp*APC)*CycBT - kdfp"*CompR +
sigma*sqrt(CycBT)*WienB

dCdk1/dt = ksfp - (kdfp'+kdfp*APC)*Cdk1 - Vwee*Cdk1 + Vc25*PCdk1
+ kdiR*CompR - kasR*(Rum1T - CompR)*Cdk1 + (kdrum' +
kdrum*(Cdk1+alpha*PCdk1))*CompR + sigma*sqrt(Cdk1)*WienB

dPCdk1/dt = Vwee*Cdk1 - Vc25*PCdk1 - (kdfp'+kdfp*APC)*PCdk1

dRum1T/dt = Vsrum - (kdrum' + kdrum*(Cdk1+alpha*PCdk1))*Rum1T +
sigma*sqrt(Rum1T)*WienR

dAPC/dt = kphapc*Cdk1*(1-APC)/(K_mapc+1-APC) -
kdpapc*PP2A*APC/(K_mapc+APC)

dPP2A/dt = -kasI*(INHaT-CompI)*PP2A + kdiI*CompI -
kasS*(Spo12aT-CompS)*PP2A + kdiS*CompS +
(kdpinh'+kdpinh*PP2A)*CompI + (kdpspo'+kdpspo*PP2A)*CompS

dINHaT/dt = kphinh*Cdk1*(INHT-INHaT) -
(kdpinh'+kdpinh*PP2A)*INHaT

dSpo12aT/dt = kphspo*Cdk1*(Spo12T-Spo12aT) -
(kdpspo'+kdpspo*PP2A)*Spo12aT

dCompS/dt = kasS*(Spo12aT-CompS)*PP2A - kdiS*CompS -
(kdpspo'+kdpspo*PP2A)*CompS

dSize/dt = mu*Size

# Flag=0 during G1/S/G2; Flag=1 during M phase
dFlag/dt = 0

# Activity functions for Wee1 and Cdc25
Wee1 =  (1-(Cdk1/Vawee)^(Qw+1))/(1-(Cdk1/Vawee)^(Nw+1))
Cdc25 = (1-(Cdk1/Vi25)^(Q25+1))/(1-(Cdk1/Vi25)^(N25+1))

# Cell growth down-regulates Rum1 synthesis and up-regulates
Cdc25 activity
Vsrum = (ksrum'+ksrum*Cdk1^r/(J^r+Cdk1^r))/Size
```

```
Vc25  = (kc25'*Cdc25 + kc25*(1-Cdc25))*Size

# Other definitions
CompR = CycBT - Cdk1 - PCdk1
CompI = PP2AT - PP2A - CompS
Vawee = k' + kdpwee*PP2A
Vi25 =  k' + kdpc25*PP2A
Vwee =  kwee'*(1 - Wee1) + kwee*Wee1

init Size=0.60, CycBT=0.63, Cdk1=0.12, PCdk1=0.5
init Rum1T=0.02, APC=0.002, PP2A=0.89, INHaT=0.12
init Spo12aT=0.006, CompS=0.005, Flag=0

# Auxiliary variables for plotting purposes
aux   Wee1 = (1-(Cdk1/Vawee)^(Qw+1))/(1-(Cdk1/Vawee)^(Nw+1))
aux Cdc25P = 1-(1-(Cdk1/Vi25)^(Q25+1))/(1-(Cdk1/Vi25)^(N25+1))
aux  INHpt = INHaT/2

# Set Flag=1 when cell enters M phase
global +1 Cdk1-MEntry {Flag=1}

# Cell divides when Cdk1 drops below MExit; set Flag=0}
global -1 Flag*Cdk1-MExit {Size=Size/2; Flag=0}

# Parameter values for MCN cell cycles
p ksfp=0.02,   kdfp'=0.015, kdfp=0.65,     kdfp"=0.15
p kasR=100,    kdiR=0.1,    kdrum'=0.125,  kdrum=21
p kphapc=0.1,  kdpapc=3,    K_mapc=1,      alpha=0.25
p kwee'=0.001, kwee=0.26,   kc25'=0.05,    kc25=0.2
p ksrum'=0.03, ksrum=0.4,   J=0.4,         r=2
p kdpwee=0.2,  Qw=1,        Nw=10,         k'=0.21
p kdpc25=0.2,  Q25=8,       N25=10,        sigma=0.015
p kasI=25,     kdiI=0.025,  kphinh=2,      kdpinh'=0.05
p kasS=25,     kdiS=0.025   kphspo=2,      kdpspo'=0.05
p kdpinh=4,    PP2AT=1,     Spo12T=0.1,    INHT=1.9
p kdpspo=4,    mu=0.005,    Mentry=0.4,    Mexit=0.1
# Note:        APCT=1,      Wee1T=1,       Cdc25T=1

# Default settings for XPP (method = Runge-Kutta)
@ total=1000, dt=0.01, maxstor=10000
@ xp=time, xlo=0, ylo=-0.03, xhi=1000, yhi=1.03
@ nplot=5, yp=Size, yp2=Rum1T, yp3=APC, yp4=Cdk1, yp5=CycBT

# Plotting variables for lower panel of simulations
#@ nplot=5, yp=Size, yp2=PP2A, yp3=Spo12aT, yp4=Cdk1, yp5=INHpt
done
```

As Fig. 1e shows, the *MCN-AF* strains proliferate more slowly than *MCN* strains. The generation times (population doubling times) of these clones are ordered:

$$\mathrm{GT}_{MCN} < \mathrm{GT}_{MCN\text{-}AF\ spo12\Delta} < \mathrm{GT}_{MCN\text{-}AF} < \mathrm{GT}_{MCN\text{-}AF\ spo12\ \mathrm{OE}} \qquad (1)$$

Furthermore, slower proliferation is associated with decreased viability, as shown by Fig. 1h: $q$ = probability of non-division ≈ 0.16 for *MCN-AF* cells and ≈ 0.09 for *MCN-AF spo12Δ* cells. A simple model of the proliferation of partially viable cells predicts that cell number will increase according to $N(t) = [2(1-q)]^{t/GT_0}$, where $GT_0$ is the population doubling time for $q = 0$ (full viability). Hence, the population doubling time for $0 < q < 1$ is $GT_q = GT_0/[1 + \log(1-q)/0.301]$. For $GT_0 = 145$ min and $q = 0.16$, we estimate $GT_q = 190$ min, and for $q = 0.09$, $GT_q = 168$ min; these estimates agree quite well with the observed generation times (185 min and 165 min) of *MCN-AF* and *MCN-AF spo12Δ* cells. For *MCN-AF spo12* OE cells (Fig. 2c; $GT_q = 250$ min, $GT_0 = 120$ min), we estimate $q \approx 0.30$.

The ordering (1) is consistent with the bifurcation diagrams, but in order to confirm the model, at least semi-quantitatively, we need to introduce some additional fluctuations to the stochastic model. In particular, we assume that the thresholds for mitotic entry and exit, $M_{\mathrm{entry}}$ and $M_{\mathrm{exit}}$, are random variables uniformly distributed over intervals $[M_{\mathrm{en}}(1-\rho_{\mathrm{en}}), M_{\mathrm{en}}(1+\rho_{\mathrm{en}})]$ and $[M_{\mathrm{ex}}(1-\rho_{\mathrm{ex}}), M_{\mathrm{ex}}(1+\rho_{\mathrm{ex}})]$. To this end, we introduce some additional lines to the 'stochastic.ode' code:

```
# Flag=0 during G1/S/G2; Flag=1 during M phase
dFlag/dt = 0
dMentry/dt = 0
dMexit/dt = 0
init Flag=0, Mentry=Men, Mexit=Mex
…
# Cell divides when Cdk1 drops below MExit; set Flag=0
global -1 Flag*Cdk1-MExit {Flag=0; Size=Size/2;
Mentry=Men*(2*ran(Ren)+1-Ren); Mexit=Mex*(2*ran(Rex)+1-Rex)}
…
p Men=0.4, Ren=0.1, Mex=0.15, Rex=0.2
```

We repeat the stochastic simulations with the modified code and the following values for the parameters $M_{\mathrm{en}}, \rho_{\mathrm{en}}, M_{\mathrm{ex}}, \rho_{\mathrm{ex}}$. In the lower half of the table, we provide appropriate initial conditions for a cell of each genotype shortly after birth. Using these parameter values and initial conditions, we simulate a clone of cells until it undergoes ~14 divisions or ends with a non-dividing progeny. Repeating these simulations many times, we estimate the probability that a cell of each genotype exits the cell cycle and arrests permanently in the terminal steady state predicted by the bifurcation diagram. The probability of arrest is reported in the table. Some representative simulations are provided in Fig. SM2.

| Genotype: | *MCN* | *MCN spo12Δ* | *MCN spo12* OE | *MCN-AF* | *MCN-AF spo12Δ* | *MCN-AF spo12* OE |
|---|---|---|---|---|---|---|
| $M_{en}$ | 0.45 | 0.45 | 0.2 | 0.45 | 0.45 | 0.2 |
| $\rho_{en}$ | 0.25 | 0.25 | 0.3 | 0.2 | 0.25 | 0.3 |
| $M_{ex}$ | 0.15 | 0.15 | 0.1 | 0.15 | 0.15 | 0.1 |
| $\rho_{ex}$ | 0.2 | 0.2 | 0.2 | 0.2 | 0.2 | 0.2 |
| CycBT | 0.4 | 0.4 | 0.115 | 0.12 | 0.12 | 0.12 |
| Cdk1 | 0.07 | 0.07 | 0.0027 | 0.0011 | 0.0011 | 0.0013 |
| PCdk1 | 0.3 | 0.3 | 0.013 | 0 | 0 | 0 |
| Rum1T | 0.026 | 0.026 | 0.3 | 0.53 | 0.53 | 0.52 |
| APC | 0.001 | 0.001 | 0 | 0 | 0 | 0 |
| PP2A | 0.94 | 0.94 | 0.99 | 1 | 1 | 1 |
| INHaT | 0.07 | 0.07 | 0.0026 | 0.0011 | 0.0011 | 0.0011 |
| Spo12aT | 0.0036 | 0.0036 | 0.004 | 0 | 0 | 0.0019 |
| CompS | 0.0031 | 0.0031 | 0.0035 | 0 | 0 | 0.0017 |
| Size | 0.5 | 0.5 | 0.4 | 0.4 | 0.4 | 0.4 |
| Mentry | 0.45 | 0.45 | 0.2 | 0.45 | 0.45 | 0.2 |
| Mexit | 0.15 | 0.15 | 0.1 | 0.15 | 0.15 | 0.1 |
| Prob arrest (simulated) | 0 | 0 | 0.04 | 0.14 | 0.07 | 0.23 |
| Prob arrest (estimated) | 0 | 0 | 0 | 0.16 | 0.09 | 0.30 |

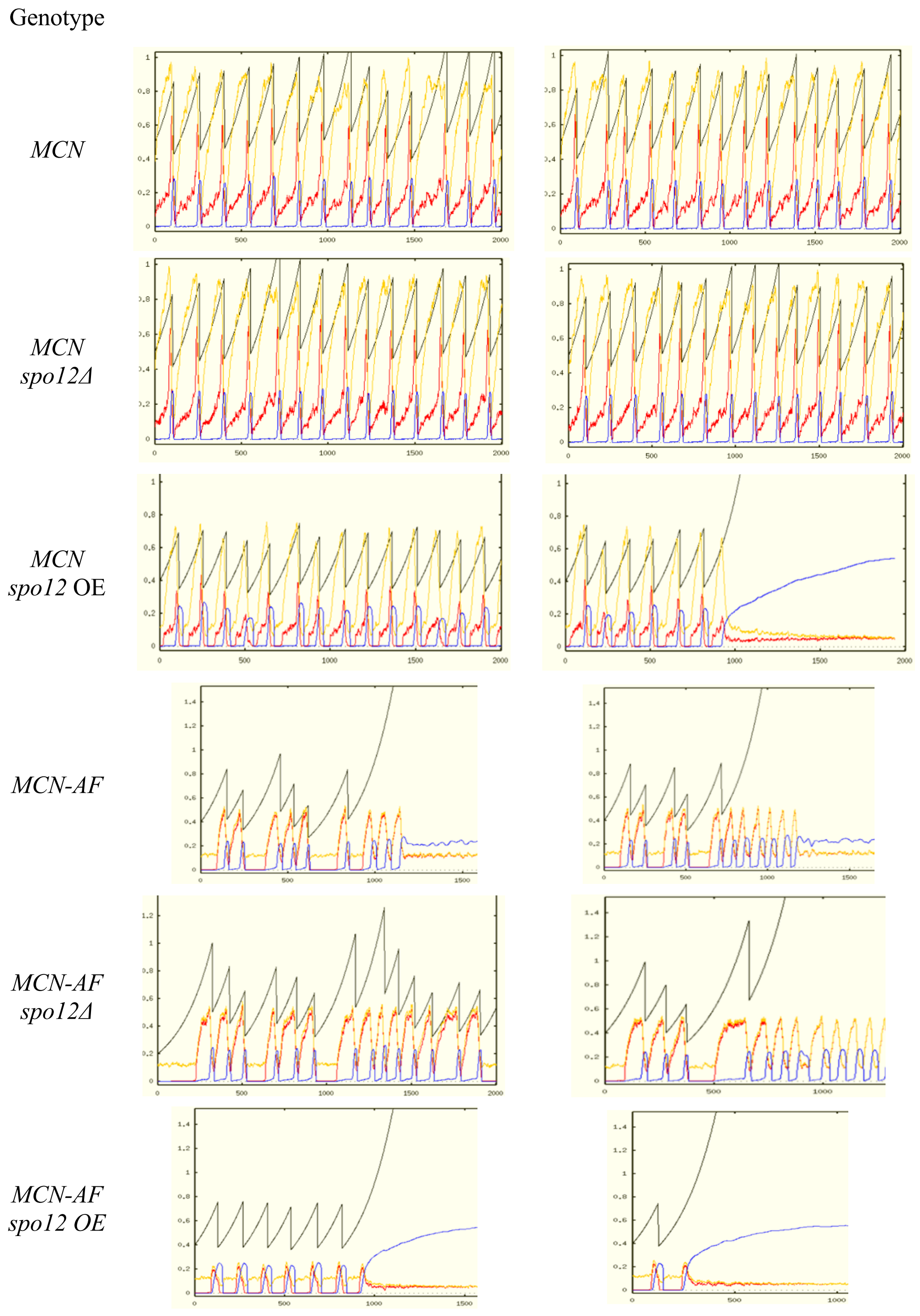


**Fig. SM2.** Representative stochastic simulations of cell clones for each of the six genotypes under consideration. Color code: black = size, yellow-orange = CycBT, red = Cdk1, blue = APC. For parameter values and initial conditions, see text.
